# A generative nonparametric Bayesian model for whole genomes

**DOI:** 10.1101/2021.05.30.446360

**Authors:** Alan N. Amin, Eli N. Weinstein, Debora S. Marks

**Affiliations:** Program in Systems, Synthetic and Quantitative Biology, Harvard Medical School, Boston, MA 02115; Program in Biophysics, Harvard University, Cambridge, MA 02138; Department of Systems Biology, Harvard Medical School, Boston, MA 02115

## Abstract

Generative probabilistic modeling of biological sequences has widespread existing and potential use across biology and biomedicine, particularly given advances in high-throughput sequencing, synthesis and editing. However, we still lack methods with nucleotide resolution that are tractable at the scale of whole genomes and that can achieve high predictive accuracy either in theory or practice. In this article we propose a new generative sequence model, the Bayesian embedded autoregressive (BEAR) model, which uses a parametric autoregressive model to specify a conjugate prior over a nonparametric Bayesian Markov model. We explore, theoretically and empirically, applications of BEAR models to a variety of statistical problems including density estimation, robust parameter estimation, goodness-of-fit tests, and two-sample tests. We prove rigorous asymptotic consistency results including nonparametric posterior concentration rates. We scale inference in BEAR models to datasets containing tens of billions of nucleotides. On genomic, transcriptomic, and metagenomic sequence data we show that BEAR models provide large increases in predictive performance as compared to parametric autoregressive models, among other results. BEAR models offer a flexible and scalable framework, with theoretical guarantees, for building and critiquing generative models at the whole genome scale.

## 1 Introduction

Measuring and making DNA is central to modern biology and biomedicine. Generative probabilistic modeling offers a framework for learning from sequencing data and forming experimentally testable predictions of unobserved or future sequences [18, 28, 54]. Existing approaches to genome modeling typically preprocess the data to build a matrix of genetic variants such as single nucleotide polymorphisms [23, 49]. However, most modes of sequence variation are more complex. Structural variation occurs widely within individuals (e.g. in cancer), between individuals (e.g. in domesticated plant populations) and between species (e.g. in the human microbiome), and methods for detecting and classifying structural variants are heuristic and designed only for predefined types of sequence variation such as repeats [12, 41, 45, 58, 67]. Ideally, we would be able to directly model genome sequencing data and/or assembled genome sequences. However, building generative models that work with raw nucleotides, not matrices of alleles, raises the extreme statistical challenges of having enough *flexibility* to account for genomic complexity, *interpretability* to reach scientific conclusions, and *scalability* to train on billions of nucleotides. Given the relevance of genetic analysis to human health, models should also possess strong *theoretical guarantees*.

Autoregressive (AR) models are a natural starting point for generative genome modeling, since they (1) have been successfully applied to protein sequences, as well as many other types of non-biological sequential data, (2) can be designed to have interpretable parameters, and (3) can be scaled to big datasets with very long sequences [56, 62]. However, since AR models are parametric models, they will in general suffer from misspecification; as we show empirically in Section 6, for genomic datasets misspecification can be a serious practical limitation not only for simple AR models but even for deep neural networks.

As an alternative strategy for building generative probabilistic models at the genome scale, we propose in Section 2 the nonparametric “Bayesian embedded autoregressive” (BEAR) model. BEAR models are Bayesian Markov models, with a prior on the lag and conjugate Dirichlet priors on the transition probabilities. The hyperparameters of the Dirichlet prior are controlled by an “embedded” AR model with parameters *θ* and an overall concentration hyperparameter *h*, both of which can be optimized via empirical Bayes. In Section 3 we show that BEAR models can capture arbitrary data-generating distributions, and establish asymptotic consistency guarantees and convergence rates for nonparametric density estimation. In Section 4, we show that the optimal *h* provides a diagnostic for whether or not the embedded AR model is misspecified and if so by how much, alerting the practitioner when the parameter estimates *θ* are untrustworthy. Besides estimation problems, BEAR models can also be used to construct goodness-of-fit tests and two-sample tests, thanks to their analytic marginal likelihoods, and we prove consistency results for these tests in Section 5. Finally we apply BEAR models at large scale, to genomic datasets with tens of billions of nucleotides, including whole genome, whole transcriptome, and metagenomic sequencing data; where comparable, BEAR models show greatly improved performance over AR models (Section 6).

Crucial to our the theoretical and empirical analysis is the statistical setting: we assume that the data *X*_1_,…, *X_N_* consists of finite but possibly variable length strings (with small alphabets) drawn i.i.d. from some underlying distribution *p*\*, and study the behavior of estimators and tests as *N* → ∞. This setup differs from common theoretical analyses of sequence models outside of biology, which typically consider the limit as the length of an individual sequence goes to infinity [24]. In biology, however, we observe finite sequences recorded from many individual species, organisms, cells, molecules, etc. and want to generalize to unseen sequences, making *N* → ∞ the appropriate large data limit.

## 2 Bayesian embedded autoregressive models

We first briefly review autoregressive (AR) models as applied to sequences of discrete characters. Let *f*(*θ*) denote an autoregressive function with parameter *θ* and let *L* denote the lag of the autoregressive model; then the AR model generates data as

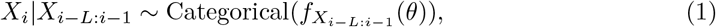

where *i* indexes position in the sequence *X* and *X*_*i−L*:*i−1*_ consists of the previous *L* letters in the sequence. Since sequence length as well as nucleotide or amino acid content is relevant to biological applications, we use a start symbol 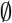 at the beginning and a stop symbol $ at the end of each sequence; letters *X_i_* are sampled sequentially starting from the start symbol and continuing until a stop symbol is drawn.

We propose the Bayesian embedded autoregressive (BEAR) model, a Bayesian Markov model that embeds an AR model into its prior. The BEAR model takes the form,

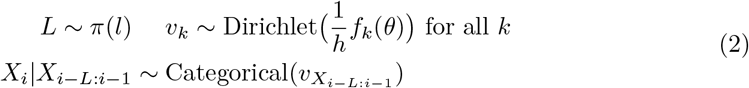

where *π*(*l*) is a prior on the lag with support up to infinity, *h* > 0 is a concentration hyperparameter, and *k* is a length *L* kmer. The BEAR model has three key properties (Fig. 1). First, the unrestricted transition parameter *υ* and lag *L* allow the model to capture exact conditional distributions of *p*\* to arbitrarily high order: *p*\*(*X_i_*|*X_i−1_*) at *L* =1, then *p*\*(*X_i_*|*X_i−2_, X_i–1_*) at *L* = 2, etc.. This property allows the BEAR model to be used for nonparametric density estimation (Section 3). Second, in the limit where *h* → 0, the BEAR model reduces to the embedded AR model (Eqn. 1). The optimal *h* provides a measurement of the amount of misspecification in the AR model (Section 4). Third, the choice of the conjugate Dirichlet prior allows the conditional marginals 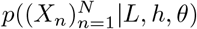 to be computed analytically, and (since *L* is one-dimensional) the total marginal likelihood 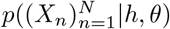 to be estimated tractably. This allows BEAR models to be used for hypothesis testing (Section 5).

**Figure 1:**
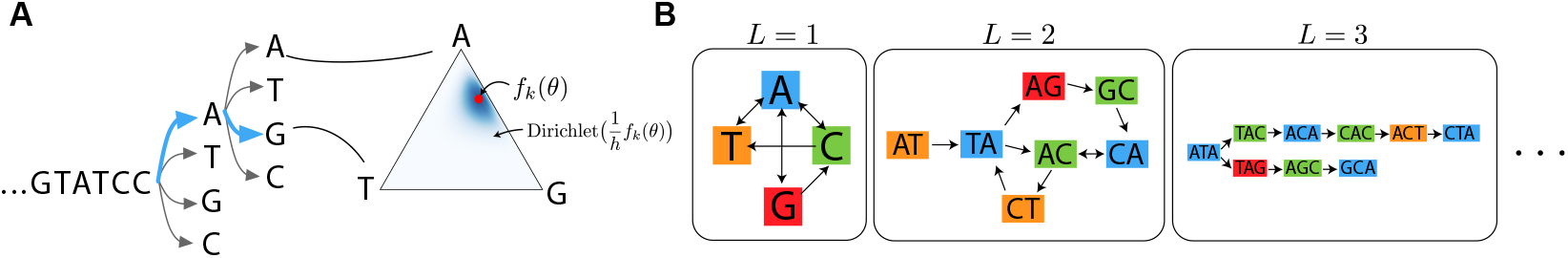
Overview of the BEAR model. (A) BEAR models employ a Dirichlet prior on Markov transition probabilities that is centered at the prediction of an AR model. (B) De Bruijn graphs showing BEAR transitions with non-zero probability under an example data-generating distribution. As the lag *L* increases, the model has higher resolution.

There are a variety of ways of performing inference in BEAR models, but for most applications we will focus on empirical Bayes methods that optimize point estimates of *L, h* and *θ*. Let #(*k, b*) denote the number of times the length *L* kmer *k* is seen followed by the letter or stop symbol *b* in the dataset 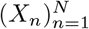. Using a high-performance kmer counter optimized for nucleotide data, KMC, we can compute the count matrix #(·, ·) for all observed kmers *k* in terabyte-scale datasets, even when the matrix does not fit in main memory (Section J.2) [35]. To optimize *h* and *θ*, we take advantage of the fact that the log conditional marginal likelihood can be written as a sum over observed kmers,

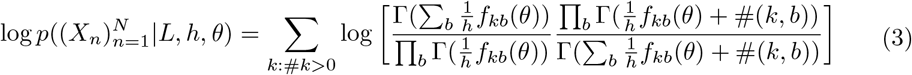

This decomposition lets us construct unbiased stochastic estimates of the gradient with respect to *h* and *θ* by subsampling rows of the count matrix (Section J.1). Empirical Bayes in the BEAR model therefore costs little extra time as compared to standard stochastic gradient-based optimization of the original AR model. Code is available at https://github.com/debbiemarkslab/BEAR.

### Toy example

We next briefly illustrate the properties and advantages of the BEAR model in simulation. We generated samples from an AR model in which *f_k_*(*θ*) depends on *k* linearly as a function of both individual positions and pairwise interactions between positions, with the strength of the pairwise interaction weighted by a parameter *β*\* (Section I.1). We first fit (using maximum likelihood) a linear AR model that lacks pairwise terms and is thus misspecified when *β*\* > 0. When the AR model is misspecified, it does not asymptotically approach the true data-generating distribution *p*\* (Fig. 2A, gray). We next computed the posterior of a vanilla BEAR model without the embedded AR in its prior, instead using the Jeffreys prior *V_k_* ~ *iid* Dirichlet(1/2,…, 1/2). The vanilla BEAR model asymptotically approaches the true data generating distribution since it is a nonparametric model; however, it underperforms the AR model in the low data regime (Fig. 2A, black). Finally, we applied our empirical Bayes inference procedure to a BEAR model with the misspecified linear AR model embedded. The BEAR model performs just as well as the AR model in the low data regime, just as well as the vanilla model in the high data regime, and better than both at intermediate values (Fig. 2A, orange). When the AR model is well-specified, the empirical Bayes estimates of AR parameters *θ* provided by the BEAR model match the maximum likelihood estimates of the AR model nearly exactly (Fig. S7). When the AR model is misspecified, however, the BEAR model provides a warning: the empirical Bayes estimate of *h* converges to a non-zero value, rather than zero (Fig. 2B). This warning emerges early: *h* converges well before the vanilla BEAR model starts outperforming the AR model.

**Figure 2:**
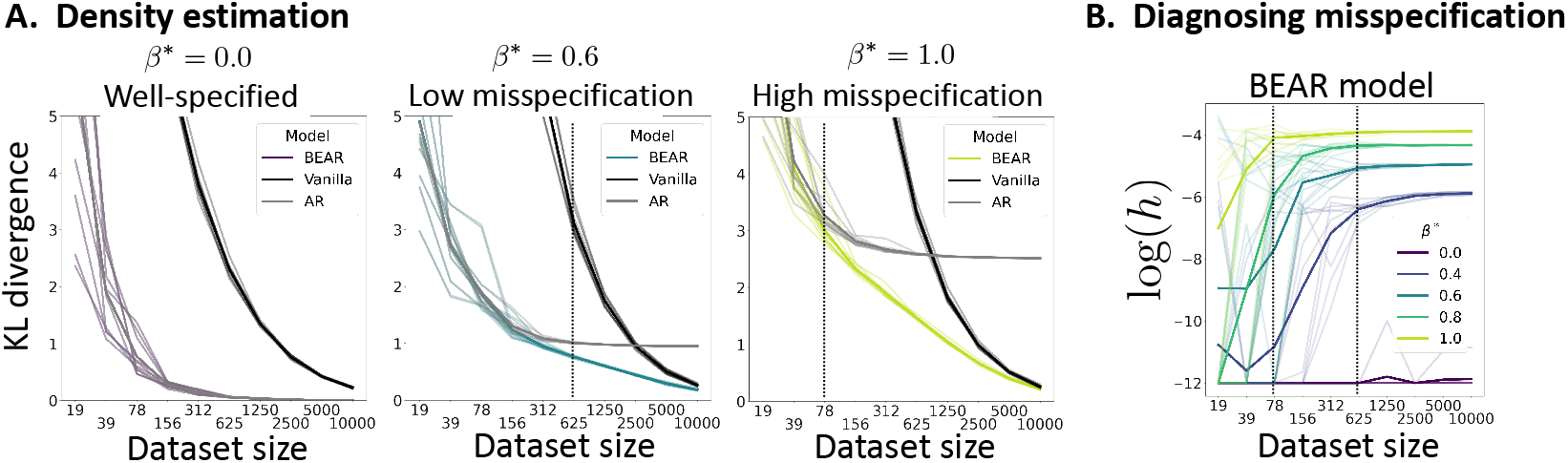
BEAR models detect and avoid misspecification without sacrificing small dataset performance. (A) Estimated KL divergence between simulated data-generating distribution *p*\* and model posterior predictive distribution, as a function of dataset size *N*. Five independent simulations were run; thin lines correspond to individual simulations, thick lines to the average across simulations. (B) The *h* misspecification diagnostic as a function of dataset size, for varying *β*\*. Dataset sizes at which *h* is close to convergence for *β*\* = 0.6 (right) and *β*\* = 1.0 (left) are marked with vertical lines.

### Related Work

The key idea behind BEAR models is to nonparametrically perturb a parametric model, following a similar strategy to the Polya tree method proposed by Berger and Guglielmi [6]. We too use independent Dirichlet priors centered at the parametric model’s predictions and use conjugacy to construct goodness-of-fit tests. BEAR models extend these ideas from one-dimensional continuous data to sequences of discrete characters.

BEAR models are closely linked to key non-generative genome analysis methods. Assembly algorithms and variant callers often analyze paths in the de Bruijn graph of a sequence dataset; in the limit *h* → ∞, samples from the posterior predictive distribution of the BEAR model, conditional on *L*, correspond to paths through the *L*-mer de Bruijn graph of the data [11, 30]. Comparisons between genomes and other sequences are often made on the basis of kmer counts; our two-sample test provides a generative perspective on this idea [3, 16, 67].

BEAR models are also connected to ideas in natural language processing, where kmers are referred to as ngrams. Under the vanilla BEAR model, the mean of the posterior predictive distribution conditional on *L* corresponds to an ngram additive smoothing model [9]. Comparisons between datasets using their ngram counts are also common in model evaluation metrics such as the BLEU score [47].

## 3 Density estimation

The density estimation problem is that of estimating *p*\* given data 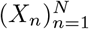. Density estimation is particularly crucial for biological sequence analysis due to its connections to fitness estimation [28]. State-of-the-art mutation effect prediction methods and clinical variant interpretation methods rely on density estimates of evolutionary sequence data, as do emerging techniques for protein design [19, 51, 54, 56]. Despite these applications, density estimation methods for biological sequences lack theoretical guarantees on their accuracy and are limited in their scale, being restricted to relatively short sequences [68]. Here, we show that the posterior distribution of the BEAR model is consistent and will concentrate on *p*\* as *N* → ∞, regardless of what *p*\* actually is, so long as *p*\* generates finite length sequences almost surely (a.s.).

We first study the expressiveness of BEAR models. Let 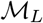 be the set of Markov models *p_υ_* with transition probabilities *υ* and lag *L* that generate finite length strings a.s.. Note that 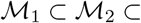…. Define the union 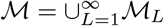. We can compare 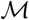 to the set of distributions over finite strings *S*, of which *p*\* is a member. In Section D we prove that,

#### Summary of Propositions 1-4

*Not all possible distributions over S are in* 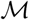. *However*, 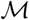 *is dense on the space of probability distributions over S with the total variation metric*. The implication of this result is that although BEAR models cannot exactly match arbitrary data-generating distributions, they can approximate *p*\* arbitrarily well as *L* increases. This makes asymptotic consistency possible.

We now show that the posterior of the BEAR will in fact asymptotically concentrate on the true *p*\*, i.e. it is consistent. For tractability, we assume in this section that the prior is fixed (we do not use empirical Bayes). The result relies on the tools for understanding convergence rates of posteriors developed in Ghosal et al. [21]. The most important assumption is that *p*\* is subexponential, meaning that for some *t* > 0, *E_p*_* exp(*t*|*X*|) < ∞ where |*X*| is the sequence length. Let 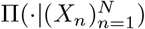 denote the posterior over sequence distributions. Let *B*(*p*,δ*) denote a ball of radius *δ* centered at *p*\*, using the Hellinger distance.

### Summary of Theorem 35

*Given M* > 0 *large enough and ϵ* ∈ (0, 1) *small enough, we have* 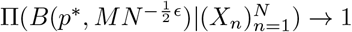 *in probability*.

A proof is in Section H and simulations in Section I.2. This result states that the posterior distribution of the model converges to a delta function at the true distribution *p*\* regardless of what *p*\* is. It also provides a rate of convergence: in a parametric model, the uncertainty would shrink as 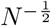, but here the rate is slower, 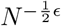, a price paid for the nonparametric model’s expressivity. The proof includes a variety of new theoretical constructions and algorithms that are used to approximate subexponential sequence distributions.

## 4 Robust parameter estimation

To derive a biological understanding of mutational processes, evolutionary history, functional constraints, etc. from sequence data, researchers must estimate model parameters (not just density). However, parameter estimates cannot in general be trusted when models are misspecified [31]. To reach robust scientific conclusions, therefore, parameter estimates should ideally come with a warning about whether or not the model is misspecified and some measurement of the degree of misspecification. Here, we study in BEAR models the asymptotic behavior of empirical Bayes estimates of the AR parameter *θ*, as well as the hyperparameter *h*, showing that *h* diagnoses misspecification in the embedded AR model.

Our analysis builds off the study of empirical Bayes consistency in Petrone et al. [48], which showed that empirical Bayes will, in general, maximize the prior probability of the true data-generating parameter value. Extending this theory to BEAR models is nontrivial, since in BEAR models the standard Laplace approximation to the marginal likelihood can fail. For theoretical tractability, as in many analyses of similar models, we fix *L* at some arbitrary and large value [27]. Define 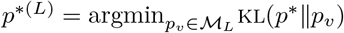 as the closest model in 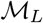 to *p*\*, and define *υ*\* such that *p_υ*_* = *p*^*(*L*)^ (note *p*^*(*L*)^ → *p*\* as *L* → ∞). We say that the AR model is misspecified “at resolution *L*” if *f* cannot approximate *p*^*(*L*)^, i.e. if there does not exist some sequence of parameter values 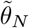 such that 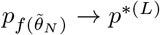; otherwise, the AR model is well-specified at resolution *L*. Now we can study empirical Bayes estimates of *h* and *θ*, denoted *h_N_* and *Θ_N_*.

### Summary of Propositions 15-20

*Let* 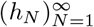 *and* 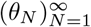 *be sequences maximizing the BEAR marginal likelihood* 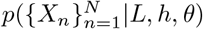 *for each N. If the model is well-specified at resolution L, then h_N_N^1/4−ϵ^* → 0 *for every ϵ* > 0 *and* 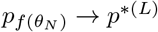 *in distribution, with both sequences converging in probability. On the other hand, if the model is misspecified at resolution L, then h_N_ is eventually bounded below by some positive (non-zero) number a.s*.. Proofs are in Section F and simulations in Section I.1. The implication of this result is that when the AR model is well-specified, *h_N_* converges to zero (at a rate that is a power of the dataset size) and *Θ_N_* converges to the parameter value *θ*\* at which the AR model matches the data (Corollary 16). On the other hand, when the AR model is misspecified, *h_N_* does not converge to zero; heuristically, we find instead that *h_N_* is approximately proportional to a divergence between *p*^*(*L*)^ and the AR model,

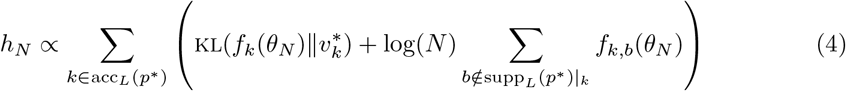

where acc_*L*_(*p*\*) = {*k* | *p*\*(#*k* > 0) > 0} is the set of kmers with non-zero probability and supp_*L*_(*p*\*)|_*k*_ = {*b* | *p*\*(#(*k, b*) > 0) > 0} is the set of transitions from *k* with non-zero probability. In summary: when fitting a BEAR model by empirical Bayes, you get, along with a parameter estimate *Θ_N_*, a value *h_N_* which tells you the amount (from zero to infinity) of misspecification in the AR model. If *h_N_* is close to zero, you can trust the estimate *Θ_N_*.

## 5 Hypothesis testing

### Goodness-of-fit test

Building generative models based on natural sequences that are accurate enough to produce novel functional sequences is a major outstanding challenge. A crucial component of the problem is model evaluation: while relative model performance may be compared on the basis of likelihood, absolute performance – whether or not the model in fact provides an accurate description of the data – is usually addressed solely on the basis of limited numbers of summary statistics, such as average amino acid hydrophobicity or sequence length [54, 56]. Given a dataset 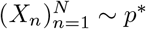 i.i.d., a goodness-of-fit test asks whether or not the data distribution *p*\* matches a model distribution 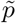. It takes into account all possible distributions *p*\* including those that differ from 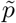 in a manner that cannot be captured by finitely many summary statistics. Our test compares the null hypothesis 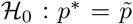 to the alternative 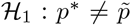 using the Bayes factor 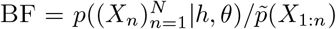, where 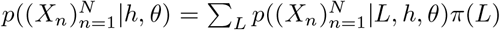 is the marginal likelihood under the BEAR model. Note that practically, the sum over *L* is straightforward to approximate, and that the test can be computed in time linear in the amount of data.

We now prove consistency. As in comparable theoretical analyses of tests based on Polya trees, for theoretical tractability we truncate the prior, setting *π*(*L*) = 0 for *L* larger than some arbitrary 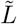 but *π*(*L*) > 0 for 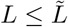 [27]. We treat *θ* and *h* > 0 as fixed.

#### Summary of Proposition 21

*If* 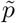 *is at least as close to p* as p^*(*L*)^ is, as measured by* KL(*p*\*||·), *then* BF → 0 *in probability as N* → ∞. *On the other hand, if p^*(*L*)^ is closer than* 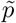, *then* BF → ∞ *in probability*. A proof is in Section G.1 and simulations in Section I.3.

An important practical limitation on nonparametric hypothesis testing is low power: since so many alternative distributions must be considered, the null hypothesis can rarely be rejected. However, Proposition 21 holds for the Bayes factor 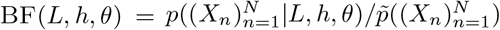 with any choice of *L, h* > 0, and *θ*. Thus in practice to increase power we can maximize the value of BF(*L, h, θ*) as a function of *L, h*, and/or *θ* (note that this approach is heuristic, since we have not proven the consistency of the maximized Bayes factor). Berger and Guglielmi [6] provide extensive methodological guidance on using analogous tests constructed with Polya trees. Based on their recommendations, we suggest first choosing *θ* such that *p_f(θ)_* is as close as possible to 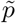, then plotting the Bayes factor as a function of *h* and/or *L* to identify the maximum value and confirm that any conclusion is robust to changes in *h* and/or *L*. Another challenge in nonparametric hypothesis testing is that it can be difficult to understand how exactly a test reached its conclusion. To identify which sequences provided the most evidence for or against the null hypothesis, we can examine the Bayes factor for each individual sequence conditional on the rest of the dataset, in analogy to the witness function used in kernel-based tests [40, 59].

### Two-sample test

A two-sample test asks whether or not two datasets 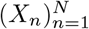 and 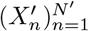 are drawn from the same distribution. Efforts to compare different sequence datasets are widespread in biology: for instance, researchers often wish to determine whether two microbiome samples, taken under different conditions or at different timepoints, are the same up to sampling noise [41]. Two-sample tests can also be used to evaluate generative sequence models that lack tractable likelihoods (for which the goodness-of-fit test proposed above does not apply) such as energy-based models or implicit models like GANs and biophysical simulators [25, 39, 44]. Assume 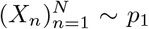 and 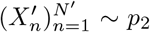 iid. Our test compares the null hypothesis 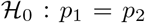 to the alternative 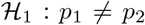 using the Bayes factor 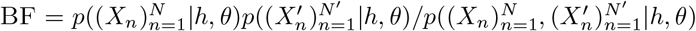. As in the goodness-of-fit case, the test can be computed approximately in time linear in the amount of data, and the same advice on increasing power and identifying important sequences holds here too.

We next prove consistency, again truncating the prior at 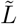 and fixing *h* and *θ*.

#### Summary of Proposition 22

*If* 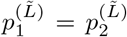, *then* BF → 0 *as N* → ∞ *in probability. Otherwise, if* 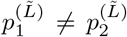, *then* BF → ∞ *in probability*. A proof is in Section G.2 and simulations in Section I.3.

## 6 Results

### Predicting sequences

We sought to evaluate BEAR models as compared to AR models on the task of predicting real nucleotide (nt) sequences. We considered eleven datasets of four different types: whole genome sequencing read data, single cell RNA sequencing read data (including from patient tumors), metagenomic sequencing read data (including from patient fecal samples) and full bacterial genomes from across the tree of life (Section K). Datasets ranged in total size from ~ 10^7^ – 10^10^ nt and in individual sequence length from ~ 10^2^ – 10^6^ nt (Table S1). 25% of data was randomly held out for testing, in the form of entire sequences (reads, genomes, etc., see Table S2); our goal was to evaluate BEAR models as density estimators, so we did not use masking (a common holdout strategy in natural language processing). We considered a linear AR model and a deep convolutional neural network (CNN) AR model with > 10× more parameters; we also designed a biologically-structured AR model which makes predictions based on a reference genome and a Jukes-Cantor mutation model (Section L.1). We then embedded each AR model to create a corresponding BEAR model. The BEAR models improve over the AR models in nucleotide prediction according to both perplexity (Table 1) and accuracy (Table S3) in all datasets, even when the model lag *L* is held fixed for comparison (Section L.3).

**Table 1:**
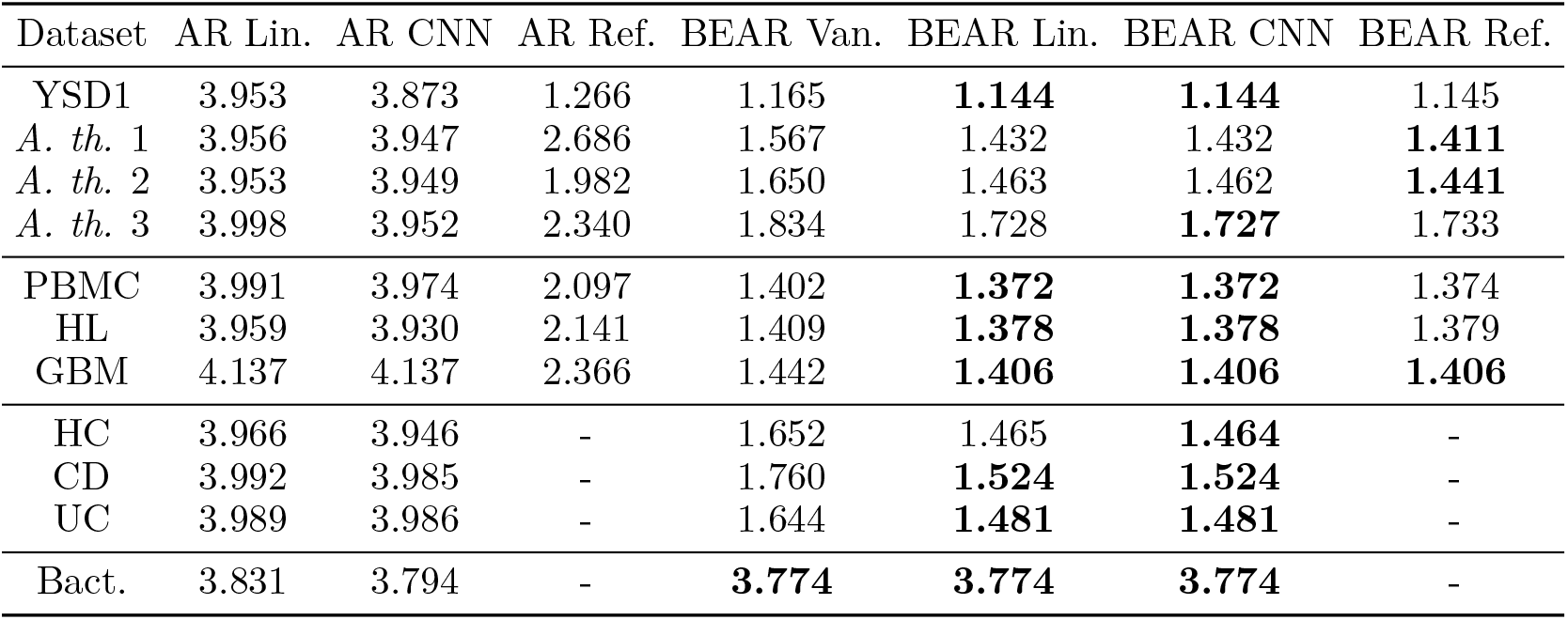
Heldout perplexity. *Whole genome sequencing data*: YSD1: A Salmonella phage. *A. th.: Arabidopsis thaliana*, a plant (datasets represent different individuals). *Single cell RNA sequencing data*: PBMC: peripheral blood mononuclear cells, taken from a healthy donor. HL: Hodgkin’s lymphoma tumor cells. GBM: glioblastoma tumor cells. *Metagenomic sequencing data*: HC: non-CD and non-UC controls. CD: Crohn’s disease. UC: ulcerative colitis. *Full assembled genomes*: Bact.: Bacteria. *Models* Van.: Vanilla (Jeffreys prior). Lin.: Linear. CNN: convolutional neural network. Ref.: reference genome/transcriptome model.

In 10 out of 11 datasets, BEAR models increase nucleotide prediction accuracy from near chance values of 30 – 35% (in the case of the linear and CNN models) to 78 – 95%, bringing genome-scale models into the realm of potential practical use (Table S3). The training time for BEAR models is essentially identical to that of AR models, aside from the time required to build the transition count matrix, which need only be done once before training all models (Fig. S13). Remarkably, the optimal lag *L* chosen by empirical Bayes is often quite short, less than 20 nt (Table S4). The improvements offered by BEAR models that use an embedded AR model over the vanilla BEAR model are modest for datasets of this size; however, sequencing experiments are often designed to collect enough data for downstream analyses. We found in an example that, if sequencing coverage was 3× instead of 100×, the improvement in prediction accuracy would have been greater than 10 percentage points instead of 0.1 (Section L.4; Fig. S14).

### Measuring misspecification

When conventional deep neural network methods fail to provide strong predictive performance, popular wisdom often ascribes the failure to too much model flexibility or not enough training data, especially in scientific applications. Examining the *h* misspecification diagnostic in the BEAR models described above, we see that this is not the case here (Table 2). The large values of *h* suggest that where the CNN fails it is not because of too much flexibility but because of too little: it suffers from misspecification. Meanwhile, the reference-based model has only two learned parameters, but is less misspecified than the CNN in all but one dataset. This too runs counter to popular wisdom in machine learning, which often assumes that when principled, low-flexibility scientific models outperform deep neural networks it is thanks to their low variance in the small data regime.

**Table 2:** Diagnostic *h*. Abbreviations as in Table 1.

| Dataset | Lin. | CNN | Ref. |
| --- | --- | --- | --- |
| YSD1 | 5.528 | 5.461 | 4.183 |
| *A. th.* 1 | 2.765 | 2.756 | 2.990 |
| *A. th.* 2 | 2.643 | 2.633 | 2.326 |
| *A. th.* 3 | 3.969 | 3.964 | 1.598 |
| PBMC | 4.167 | 4.145 | 3.762 |
| HL | 4.050 | 4.038 | 3.581 |
| GBM | 4.172 | 4.154 | 3.238 |
| HC | 4.668 | 4.651 | - |
| CD | 3.096 | 3.094 | - |
| UC | 3.843 | 3.835 | - |
| Bact. | 0.010 | 0.003 | - |

### Generating samples

BEAR models are generative and can be used to sample new sequences. We sampled extrapolations from the end of a read sequence recorded in a plant (*A. thaliana*) whole genome sequencing experiment, and compared to an alternative non-probabilistic extrapolation method that is widely used in biology, local assembly (Fig. 3A; Section M). In this example the assembly algorithm SPAdes returns four possible assemblies, a relatively large number compared to other reads in the dataset (Fig. 3A stars) [5]. Samples from the BEAR model include these four possibilities, but also many more, some with higher probability. The distribution over possible nucleotide choices under the BEAR model is much wider than the number of assemblies would suggest: it has a perplexity of 1.4 per position (on average across samples) at the beginning of the extrapolation, and a perplexity of 2.7 at 50 nucleotides (Fig. 3B). These observations suggest that SPAdes, which does not provide a measurement of uncertainty, may not be capturing the full range of possible sequences.

**Figure 3:**
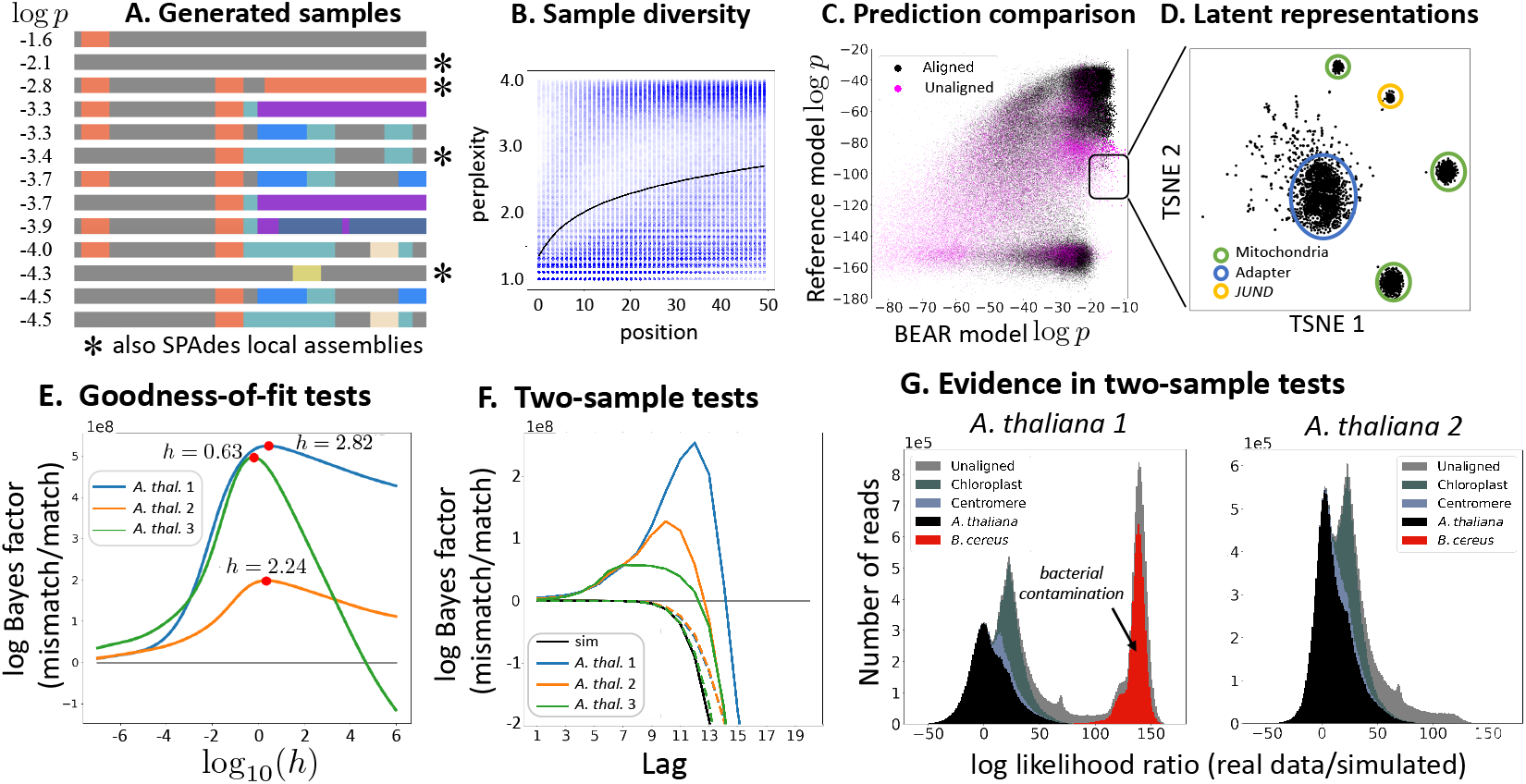
Generation, visualization and testing. (A) Sample extrapolations, colored to denote distinct paths through the *L*-mer de Bruijn graph. (B) Perplexity of the next Markov transition under the BEAR model, for each position of each sampled extrapolation, with the average across samples shown in black (Section M). (C) Log probability of each read in the HL dataset under the BEAR model and a model built from the reference transcriptome. Reads are colored by whether or not they map to the reference. (D) Latent representations of the reads highlighted in C, visualized using tSNE, with clusters annotated as likely coming from mitochondria, the sequencing adapter, or transcripts of the gene *JUND* (Section N). (E) Goodness-of-fit test Bayes factor as a function of hyperparameter *h*. (F) Two-sample test Bayes factor as a function of lag *L*. Black line compares simulated data to simulated data; dashed lines compare subsampled real data to subsampled real data; solid lines compare real data to simulated data. (G) Log probability of each read under the real data BEAR model minus the log probability under the simulated data BEAR model (Section O).

### Visualizing data

Methods for learning local representations or features of biological sequences can be powerful tools for visualization and semisupervised learning [7]. One approach to extracting such representations is to learn a generative model *q*(*X*_1_,…, *X_L+1_*) of kmers, for instance using a variational autoencoder. While such models are not autoregressive, the small size of the DNA alphabet makes it tractable to estimate the conditional *q*(*X_L+1_*|*X_1:L_*) by Bayes rule, and this conditional can then be embedded into a BEAR model. We applied this strategy to probabilistic PCA. We visualized in low dimensions the inferred latent representation for a model trained on a single cell RNA sequencing dataset (HL), and were able to assign annotations to clusters, including those containing unmapped reads (Fig. 3CD; Section N). The BEAR model however raises the warning that the model is misspecified (*h* = 4.836), suggesting there may be richer latent structure yet to discover.

### Testing hypotheses

The question of when and how microbiomes change is widespread, but has in the past relied on summary statistics of sequencing datasets [41]. Schreiber et al. [55] studied changes in patient urine microbiomes before and after kidney transplant, and performed both unbiased metagenomic sequencing and diagnostic quantitative polymerase chain reaction (qPCR) for a specific virus associated with complications (JC polyomavirus). They found evidence of donor-to-recipient viral transmission in 5 cases out of 14. We applied the BEAR two-sample test to patients’ metagenomic sequencing data before and after transplantation, using the vanilla Jeffreys prior and integrating over lags, in order to detect changes; the test rejects the null hypothesis in all 5 cases where there was transmission, and accepts the null hypothesis in all but one of the remaining 9 cases (Table S6; Section O.1). These results show, in a small example, that the two-sample test has sufficient power to detect microbiome changes in real data, and can be consistent with more specific tests.

We next applied BEAR hypothesis tests to evaluate generative models. We evaluated the reference-based AR model described above using the BEAR goodness-of-fit test. The test identifies considerable evidence (log Bayes factor > 10^8^) for misspecification in each *A. thaliana* whole genome sequencing dataset, and this conclusion is robust to a wide range of *h* values (Fig. 3E; Section O.2). Next, we evaluated a detailed simulation model (ART) that is intended to generate likely reads of a given reference genome [29]. The model lacks tractable likelihoods, so we use a two-sample test. When integrating over all lags, the test accepts the null hypothesis, but if we examine the test results for individual lags *L* to increase power, we see evidence of differences (Fig. 3F; Section O.2). To understand the source of these differences, we examined the conditional Bayes factor for individual reads, discovering clusters of reads that are poorly explained by the simulation model (Fig. 3G). One group mapped to chloroplasts, an organelle with its own genome that is variable in copy number; reads mapping to centromeres, an area of the plant genome for which the reference genome is considered unreliable, were also poorly explained by the simulation model. In one dataset we found a cluster of outliers that did not map to *A. thaliana* at all, and instead mapped to a common soil bacteria, *Bacillus cereus*, presumably a contaminant in the experiment (Fig. 3G, left). These results illustrate how BEAR hypothesis tests can be used not only for testing but also for detailed model criticism.

## 7 Discussion

In this article we proposed the nonparametric BEAR model, studied its theoretical properties, and developed algorithms and implementations for terabyte-scale inference. BEAR models substantially outperform standard AR models on a variety of datasets, and come with extensive theoretical guarantees, including for density estimation, misspecification detection, and hypothesis testing. BEAR models are closely connected to non-probabilistic genome analysis methods, such as de Bruijn graph assembly, but provide an alternative that is uncertainty-aware. Note, however, that BEAR models do not explicitly account for paired-end read information, or other sources of long-distance information; this is an important area for future work. While there has been little previous empirical or theoretical work in the machine learning literature on generative models of full genomic, transcriptomic or metagenomic sequences, we hope BEAR models provide a useful starting point.

## Acknowledgments and Disclosure of Funding

We thank Jean Disset for a small scale version of the kmer counting code and Rob Patro for crucial advice on large scale kmer counting. We thank Tessa Green, Chris Sander, and Elizabeth Wood for discussion and advice. We thank Winnie Wang for their illustrations used in the theory section of the supplementary materials. We thank members of the Marks Lab for discussion and ideas. E.N.W. is supported by the Fannie and John Hertz Foundation. D.S.M. is supported by the Chan Zuckerberg Initiative.

## A Broader impact statement

The social impact and bioethics of genetics is a complex and well-studied subject. Here, we briefly highlight some of the ways the methods presented in this article intersect with major issues. Health: statistical genetics methods have been crucial in diagnosing disease and dissecting its mechanisms, and we hope that the methods presented here will further this research. On the other hand, a detailed understanding of personal genetic information can be used as the basis for discrimination. Technology: as generative models, BEAR models may be unusually useful in designing new sequences such as therapeutics. Genetic engineering, however, is a dual-use technology. Culture: popular understanding of genetics is often bound up in the idea that genomes are relatively fixed and change only very slowly over time, which feeds into concerns over the “naturalness” of genetic modification and beliefs about the static nature of race and ethnicity. These public perceptions are likely partially a consequence of scientific models and methods that often simply ignore complex genetic variation and analyze individual genomic data by comparing it to “reference” individuals. BEAR models contribute theoretical and statistical methods for working with complex variants and without relying on references.

## B Overview of supplemental material

Sections C–H are our theoretical results. Section I describes our simulation experiments. Section J details how we implemented scalable inference for BEAR models. Sections K–O provide details on our empirical results based on real data. The Datasets.xlsx file contains information on all the datasets, including links or accession numbers for public databases. Code is available at https://github.com/debbiemarkslab/BEAR.

## C Theory Introduction

BEAR models can be used to address a variety of different estimation and testing problems, and the theoretical questions that arise in each case are related but distinct. One crucial, high-level distinction is between the “finite-lag case” (where we assume the model lag *L* is finite) and the “infinite-lag case” (where we allow the model lag *L* to approach infinity). In addressing nonparametric density estimation, it is crucial to consider the infinite lag case, since it is likely in practice that the true distribution can only be matched in the infinite *L* limit. On the other hand, when it comes to diagnosing misspecification or constructing hypothesis tests, the finite lag case is more acceptable since it is likely in practice that any differences between the model and the data, or between two datasets, will be reflected in finite marginals of the data distribution. The finite lag case is complicated by the fact that it is likely that many kmer-to-base transitions have extremely low probability in practice; even on massive datasets, we observe many transitions with no counts whatsoever. To deal with this case, we develop theoretical tools to accommodate the possibility that some transitions truly have probability zero under the data generating distribution.

An essential and innovative aspect of our formalism is the focus on “subexponential” sequence distributions that obey an exponential moment bound on their length. Our choice to consider sequence distributions that have no upper bound on the lengths of sequences they produce separates our theory from the theory of distributions on finite sets. On the other hand, moment bound assumptions separate our theory from the theory of distributions on countable sets.

The theory will be organized as follows. Section D describes basic theoretical properties finite-lag Markov sequence models, including their expressiveness and subexponentiality. Subexponential sequence models will be introduced in general here. Section E demonstrates consistency of inference with a fixed lag and in model selection between lags. A connection is established between effective model dimensions and topologies of de Bruijn graphs. Section F describes the behavior of the model when inference proceeds by empirical Bayes. The parameter *h* is established as a descriptor of misspecification. Section G describes theoretical guarantees on the behavior of goodness-of-fit and two-sample tests. Finally, section H demonstrates consistency in the infinite lag case. Later sections depend on definitions and results established in previous sections with the exception that section H may be read immediately after reading the definitions at the top of section E.

### C.1 Notation

We consider an alphabet 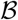 with more than one letter. Define 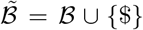 where $ is interpreted as the stop symbol, i.e. $ may only appear as the last letter of a sequence. Also define the set of strings of the alphabet 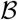 of length *L* that start with any number (including 0) of repeated 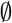 symbols, 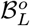. For a sequence *X* of letters in 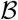, possibly terminated by $, we define |*X*| as its length, including the stop symbol $ but not any start symbols 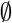. For two strings *X,X*′ define #*X*′(*X*) the number of occurrences of *X*′ as a substring in *X* and, if *X* is not terminated by $, define (*X, X*′) as the concatenated string. We also define the substring from index *i* to *j* (inclusive) of *X* as *X_i:j_*.

Define the set *S* of all finite sequences terminated by a stop symbol and give it the discrete topology. Note that *S* is countable. Say *p* is a distribution of *S*. We will use 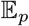, or 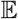 if there is an unambiguous data-generating distribution, to denote taking an expectation; for example, 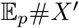 is the expected number of occurrences of the substring *X*′ in sequences drawn from *p*. For a sequence *Y* possibly not terminated by a stop symbol, we define *p*(*Y*…) = *p*({*X* ∈ *S* | *X_i_* = *Y_i_* ∀*i* ≤ |*Y*|}). We also define subexponential moment bounds, an assumption we will make great use of:

#### Definition 1

(Subexponential sequence distributions). We say a distribution *p* on *S* is subexponential if for a *t* > 0, 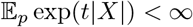.

For a random variable *Z* on a probability space with probability *P*, and a measurable set *A* in the sample space, we define

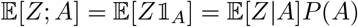

where 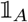 is the random variable with 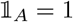 on *A* and 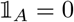 outside of *A*. As well, for two real sequences 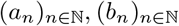, both possibly undefined for small *n*, we write *a_n_* ≲ *b_n_* to mean that there is a positive constant *C* such that eventually *a_n_* ≤ *Cb_n_*. We write *a_n_* ≲ *b_n_* when *a_n_* ≲ *b_n_* and *a_n_* ≳ *b_n_*. We define *a* ˄ *b* as the minimum of *a* and *b*, and *a* ˅ *b* as the maximum.

## D Finite-lag Markov models

In this section we define finite-lag Markov models, and then study the expressiveness of the model class. After defining finite-lag Markov models, this section will concern itself with the expressiveness of the model class. We first show that while there are sequence distributions over *S* that are not finite-lag Markov models, the set of finite-lag Markov models is nevertheless dense in the space of distributions over *S*. We then show that finite-lag Markov models are subexponential.

The class of finite-lag Markov models is defined to be

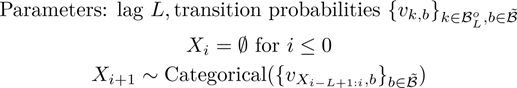

stopping generation when a $ symbol is drawn and with parameters picked so that |*X*| < ∞ a.s.. These models are equivalent to Markov processes on the set 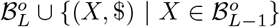. The requirement that generated sequences be finite length a.s. is equivalent to the requirement that for every 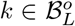 that is Markov-accessible, there is another 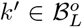 that is Markov-accessible from *k* such that 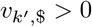. Call *p_υ_* a probability distribution generated this way with parameters *L, υ*. Call the set of such probability distributions with lag 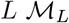. Define the set of all finite lag Markov models 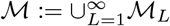 and note 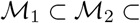…. Defining 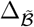 as the 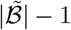-dimensional simplex with coordinates indexed by 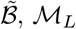 is parametrized by transition probabilities in 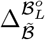. This parametrization is not defined everywhere on the boundary and is not injective as if an *L*-mer *k* is not Markov-accessible by a distribution *p_υ_*, the vector of probabilities *υ_k_* does not affect *p_υ_*’s distribution. This parametrization is continuous in the sense of the topology described by proposition 2.

We first give some examples of simple sequence distributions that are not finite-lag Markov.

### Proposition 1.

*Not all possible distributions over S are in* 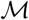.

*Proof*. Let 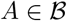 and *p*\* be a distribution over finite sequences that puts probability *a_i_* on the sequence *A* × *i*:= *A*…*A* of length *i* with 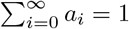. Assume 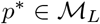 with transition probabilities 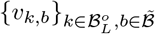.

For *i* ≤ *L*, define 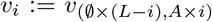, i.e. the vector of transition probabilities from the *L*-mer that is *L* – *i* 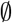 symbols followed by *i A* symbols. For *i* ≥ *L* call *υ_i_*: = *v_L_*.

Notice that for any *i*, the $-component of the vector *υ_i_* is 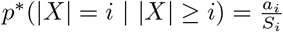 where 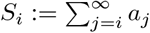. Thus the *A*-component is 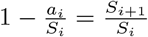. By the definition of the sequence 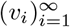, it is constant for *i* ≥ *L*. Call *α*: = *S_L+1_/S_L_* = *υ_L,A_* = *υ_i,A_* = *S_i+1_/S_i_* for all *i* ≥ *L*. Thus for all *i* > *L*, 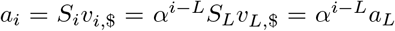. Thus the sequence *a_i_* eventually decays exponentially and, as examples, it is impossible that *a_i_* ~ 1/*i*! or *a_i_* ~ 1/*i*^2^.

Next we show that 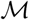 is dense in the set of probability distributions on *S*. To speak of density, we review the topology and types of convergence on the set of distributions of *S* in this next proposition.

### Proposition 2.

*The topology of convergence in total variation, convergence in distribution, and pointwise convergence of the probability of each X* ∈ *S are identical*.

*Proof*. Pointwise convergence of the probability of each *X* ∈ *S* implies convergence in total variation by Scheffé’s lemma. It is also known that the topology induced by the total variation metric is stronger than the topology of convergence in distribution. Finally, since for each *X* ∈ *S*, the set {*X*} is open and closed, so that the Portmanteau lemma shows that convergence in distribution implies pointwise convergence.

### Lemma 3.

*Say p is a distribution on S. There is a lag L Markov model, p_L_, such that for all X ∈ S, if* |*X*| ≤ *L, p_L_*(*X*) = *p*(*X*), *and if* |*X*| > *L*, 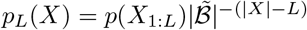.

**Figure S1:**
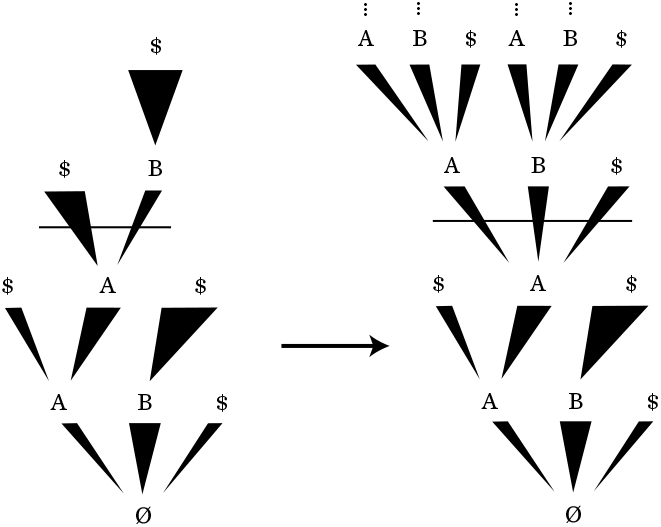
Example application of this construction to the distribution on the left. Transition probabilities for kmers smaller than *L* = 2 are those defined by the original distribution, while those for larger kmers are all 1/3. The thickness of each line denotes the probability of the transition.

*Proof*. For all 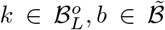, if there is a start symbol 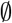 in *k*, define 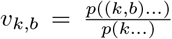, otherwise, define 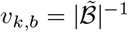. It is clear *p_υ_* satisfies the properties of *p_L_* (Fig S1).

### Corollary 4.

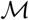 *is dense in the set of distributions of S*.

*Proof*. Define *p* to be a distribution on *S* with finite support, 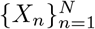. Pick an *L* > |*X_n_*| for all *n*, so that with the definition of lemma 3, *p_L_* = *p* and thus 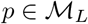. Now note that any distribution on *S* can be approximated at finitely many points in *S* arbitrarily well by distributions with finite support. The result follows from proposition 2.

### Proposition 5.

*Finite-lag Markov models are subexponential*.

*Proof*. Say 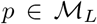 for some *L*, with transition probabilities *υ*. Every 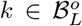 that is Markov-accessible by *p* has a 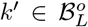 that is Markov-accessible from *k* in less than *s_k_* transitions such that *υ_k′,$_* > 0. Thus, 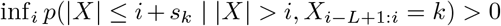. Define *s* = max_*k* accessible_ *s_k_, q* = inf_*i*_ *p*(|*X*| ≤ *i* + *s* ||*X*| > *i*) > 0. Now note, for any positive integer *m*, 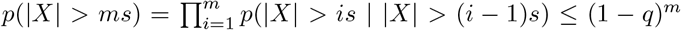. For a random variable *Z* ~ Geom(*q*),

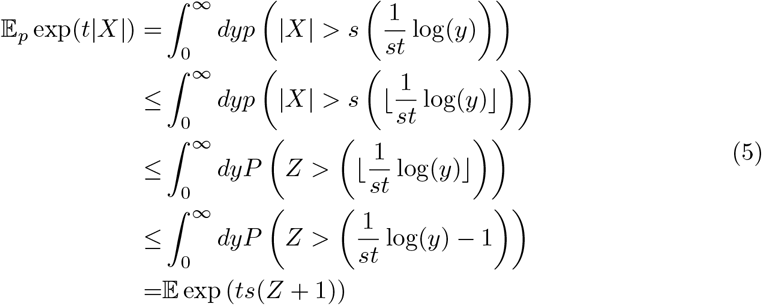

The integral is finite for some *t* > 0 as geometric random variables are sub-exponential.

## E Consistency in the finite *L* case

In this section we consider fitting to data BEAR models with fixed hyperparameters *h* and *θ* (that is, standard Bayesian Markov models). We first study the asymptotic behavior of the posterior over *υ*, the transition probability parameter, conditional on a particular lag *L*. We prove a Wald-type consistency result, showing that the posterior concentrates on a neighborhood of the true data-generating parameter value *υ*\*, if such a value exists; when *p*\* is not in the model class 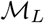, the posterior over *υ* concentrates at the point *υ*\* corresponding to the distribution 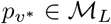 closest in KL divergence to *p*\*. We next study the asymptotic behavior of the posterior over the lag *L*, building on the theory of nested model selection since *L* is a discrete variable. We show that the posterior concentrates at the true data-generating value *L*\* when such a lag exists (i.e. when there is some *L*\* such that 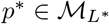), and otherwise diverges. At a high level, neither of these results are surprising, and they would be expected to hold in general for well-behaved Bayesian models. The details of the model’s asymptotic behavior, however, turn out to be somewhat unusual; as we shall see, the fact that some transitions from a particular kmer *k* to a base *b* have probability zero under the data-generating distribution *p*\* can complicate the normal story of Bayesian asymptotics.

To describe the possible kmer-base transitions, we define, for a distribution on *S*, *p*, and a lag *L*, the set of accessible kmers 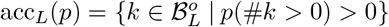 and transitions 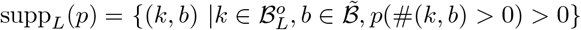. Define also, for any particular a 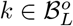, the set of allowed transitions 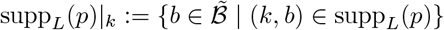. Define the restriction of the parameter space 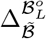 to the support of *p*\*, 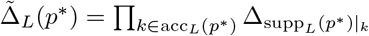. If 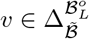, we will often use the abbreviation supp(*υ*) = supp_*L*_(*p_υ_*) for convenience.

Say *p*\* is a distribution on *S* and *L* is a lag. Define the transition probabilities *υ*\*, corresponding to the closest model in 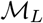 to *p*\* (as measured by KL), as

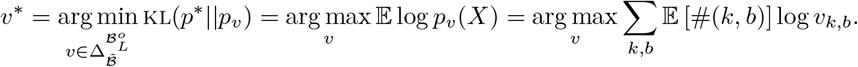

Unlike for many other statistical models studied in other contexts, here we can easily compute the closest model to the data-generating distribution: using Lagrange multipliers, one may see that for all 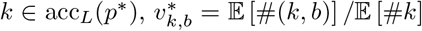. We then define 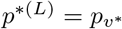 as the best approximation to *p*\* in 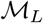. Note supp(*υ*\*) = supp_*L*_(*p*^*(*L*)^) = supp_*L*_(*p*\*).

We now ask whether Bayesian inference on 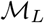 is consistent, i.e., whether the posterior converges to a point mass at *p*^*(*L*)^, even in the case where supp_*L*_(*p*\*) is not all of 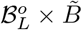. The result is a classic Wald-type argument, adapted from theorem 2.3 of Miller [43] and theorem 1.3.4 in Ghosh and Ramamoorthi [22]. The primary difficulty in the proof is that these previous theorems assume the true parameter value lies on the interior of the parameter space and rely on uniform convergence of the mean log likelihood in a neighborhood around the true value. In our case, we can have 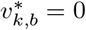, so that the true parameter value lies on the boundary of its space 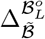 and the likelihood function diverges at this boundary point.

### Theorem 6.

*Say p* is a distribution on S with* 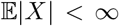. *Say* Π *is a prior on* 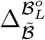 *that assigns probability 0 to the set of υ with* 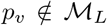. *Say X*_1_, *X*_2_,⋯ ~ *p*\* *iid. Call* 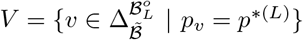 *and assume that it is not disjoint from the support of* Π. *Then for all open sets U containing V*,

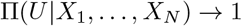

*a.s.. As a probability distribution on the space of measures on S*, 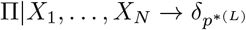.

*Proof*. Define *υ*\* as the transition probabilities of *p*^*(*L*)^. Define 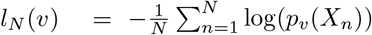, which is continuous in *υ* and 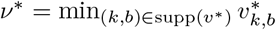. Note that

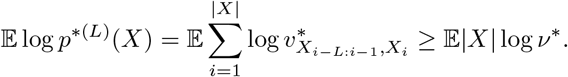

First we show that the likelihood of the data is eventually small in a neighborhood of the boundary. Pick an *η*_1_ > 0. Say (*k, b*) ∈ supp(*υ*\*) = supp_*L*_(*p*\*) and define *q_k,b_* = *p*\*(#(*k, b*) > 0) which is positive. Pick a positive

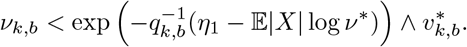

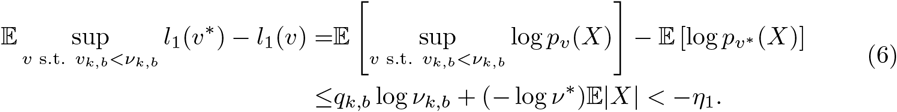

Thus defining 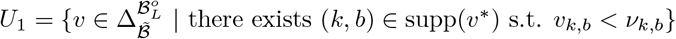, a.s., for large enough *N*, *l_N_*(*υ*\*) – *l_N_*(*υ*) < − *η*_1_ for all *υ* ∈ *U*_1_ by the SLLN.

Call the complement of *U*_1_ *K. K* is compact and for all *υ* ∈ *K*, supp(*υ*\*) ⊆ supp(*υ*). Note that *V* is compact and in the interior of *K*. Pick a positive *ν_K_* which has, for every (*k, b*) ∈ supp(*υ*\*), *ν_K_* – *ν_k,b_*. Then

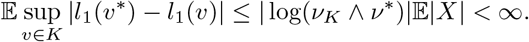

Then by theorem 1.3.3 in Ghosh and Ramamoorthi [22], a.s., *l_N_*(*υ*\*) – *l_N_*(*υ*) converges uniformly to KL(*p*\*||*p*^*(*L*)^) – KL(*p*\*||*p_υ_*) ≤ 0 on *K* (note, for the application of theorem 1.3.3 in Ghosh and Ramamoorthi [22], this quantity is well defined even if *p_υ_* is not a distribution over finite strings).

Now pick an open neighborhood *U* of *V*. By the continuity of *υ* ↦ KL(*p*\*||*p_υ_*), since *K* \ *U* is compact, inf_*υ∈K\U*_ KL(*p*\*||*p_υ_*) > KL(*p*\*||*p*^*(*L*)^) otherwise there would be a *υ* ∈ *V* \ *K*. Thus we can pick a positive KL(*p*\*||*p*^*(*L*)^) + *η*_2_ < inf_*υ∈K\U*_ KL(*p*\*||*p_υ_*). Since *υ* ↦ KL(*p_υ_*\* || *p_υ_*) is continuous and *K* is a neighborhood of *V*, there is an open *U*_2_ ⊂ *K* ∩ *U* containing *V* such that one can pick an *η*_3_ with sup_*υ∈U*_2__ KL(*p*\*||*p_υ_*) – KL(*p*\*||*p*^*(*L*)^) < *η*_3_ < *η*_1_ ˄ *η*_2_. Then a.s. eventually, *l_N_*(*υ*\*) – *l_N_*(*υ*) < −*η*_2_ for all *υ* ∈ *K* \ *U* and *l_N_*(*υ*\*) – *l_N_*(*υ*) > −*η*_3_ for all *υ* ∈ *U*_2_. Thus, a.s. for large enough *N*,

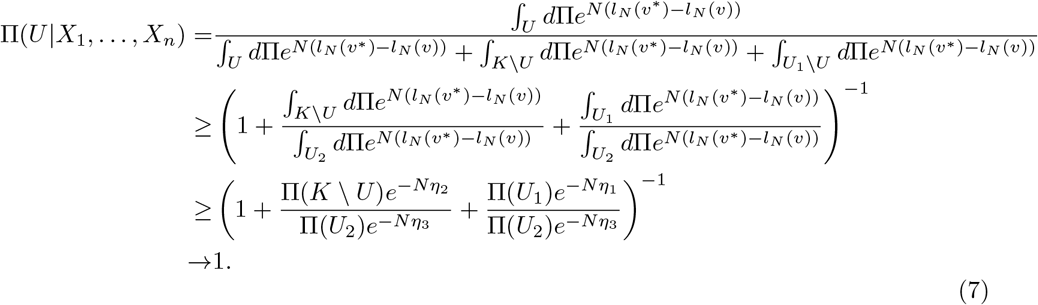

Finally, as a probability distribution on the space of measures on *S*, Π|*X*_1_,…, *X_n_* → *δ_p^*(*L*)^_*. This follows from the fact that the prior and thus posterior probability of 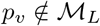 is 0 and so one may push forward the measure from 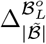 to the space of probability measures on *S*. The image of *V* is a point *p_υ*_*. Since this mapping is continuous, it preserves the weak convergence of the measure, in this case to a point mass.

Next we will study the posterior distribution of the BEAR model over the lag *L*, showing under general assumptions that the posterior concentrates on the true data-generating value *L*\* (when such a value exists). Our analysis builds off of standard asymptotic analyses of nested Bayesian model selection, since models with different lags are nested, i.e. 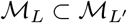 when *L*′ > *L*. Typically, when a simpler model (e.g. 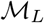) is nested inside a more complex model (e.g. 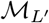), and the data-generating distribution *p*\* is in the simpler model, the log Bayes factor comparing the two models will asymptotically prefer the simpler model and scale as 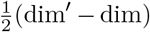 log *N* where dim′ is the dimension of the parameter space in the more complex model and dim is the dimension in the simpler model [14]. This *O*(log *N*) term, which is independent of the prior, can be thought of as originating from the Laplace approximation to the marginal likelihood; it is the basis of such widely used model-selection techniques as the Bayesian information criterion.

Somewhat surprisingly, the fact that some transitions may have probability zero 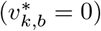 changes the asymptotic behavior of the log Bayes factor, in particular by altering the dimension factor dim′ – dim. In effect, dimensions of the parameter space corresponding to kmers that occur with probability zero under *p*\* do not contribute to the dimension count, while dimensions for which 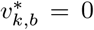 do not count as full dimensions; this leads to the notion of an “effective model dimension”, defined as 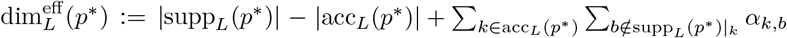 where *α_k,b_* is the concentration of the Dirichlet prior. This effective dimension depends the data-generating distribution *p*\* and on the prior hyperparameters, not just on *L*. Note that the unusual asymptotic behavior of BEAR models does not just come from their Markov structure; even in the everyday example of a Dirichlet-Categorical model, if some outcomes of the Categorical distribution have probability exactly zero under the true data-generating distribution, the standard Laplace approximation does not hold, and the Dirichlet prior contributes *O*(log *N*) terms to the log marginal likelihood [53].

### Theorem 7.

*Say p* is a distribution on S with* 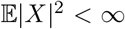 *and say X*_1_, *X*_2_, ⋯ ~ *p*\* *iid. Given L, consider a* 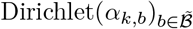 *prior on the simplex in* 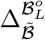 *corresponding to the L-mer k. For all L, assume α_k,b_* > 0 *for* (*k, b*) ∈ supp_*L*_(*p*\*) (*otherwise* 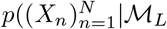) *is eventually* 0 *a.s*.).

*Define* 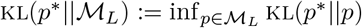. *Given L*_1_ ≠ *L*_2_, *if*^2^ 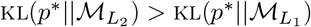,

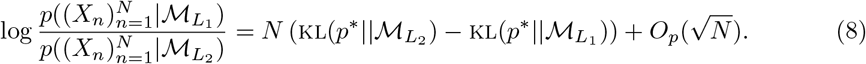

*Otherwise, if* 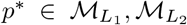 *and, defining, for a lag L*, 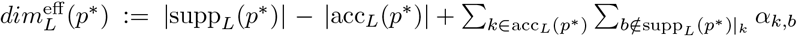,

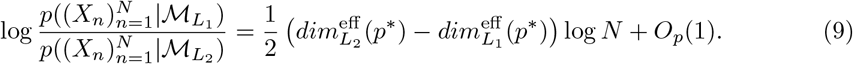

*Proof*. For a lag *L*, note 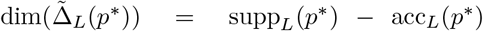. Put a 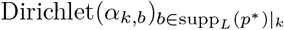 prior on each 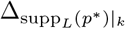. Call 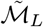 the set of probability distributions described by 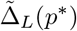. We will show that KL(*p*\*||*p*.) is maximized in the interior of 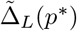 so that the asymptotics of the marginal likelihood 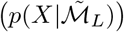 are well understood.

In 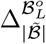 however, there are dimensions that correspond to k-mer - base transitions that are impossible under *p*\*. Using the symmetry of the Dirichlet prior, we can de-couple the asymptotics of these excess dimensions from the asymptotics of the much more “natural” space of 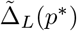:

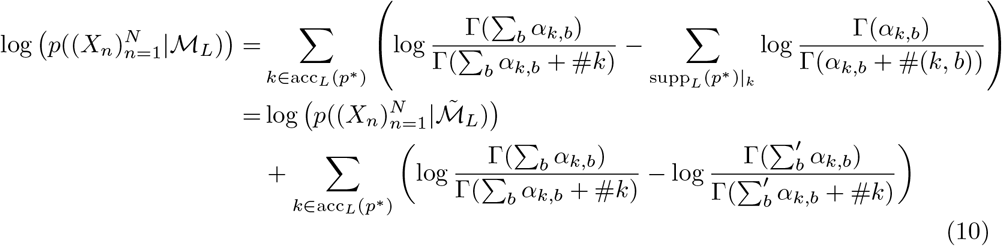

where 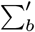 is a sum over the *b* ∈ supp_*L*_(*p*\*)|_*k*_, and where #*k* in this case is 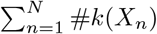 and #(*k, b*) is similar. We will deal with each of these terms in turn.

To analyze the first of these terms, we first check regularity conditions. For 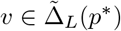 and strings *X*_1_,… *X_N_*, define

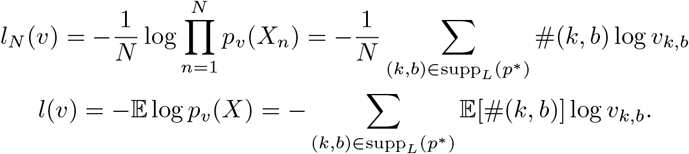

Call *υ_n_* the minimizer of *l_N_* and *υ*\* the minimizer of *l*. Note *υ*\* is also the minimizer of *υ* ↦ KL(*p*\*||*p_υ_*) for 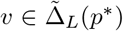 and has 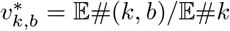. In particular *p_υ*_* = *p^*(L)^* so that 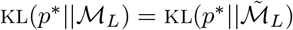. One may check that *l_N_* is *C*^∞^, and, by seeing that it is a sum of convex functions, convex. Calling *D^m^* the m-th derivative operator (*D*^0^ the identity), ||·|| some norm on 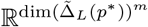, and *E* some set whose closure is in the interior of 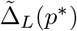

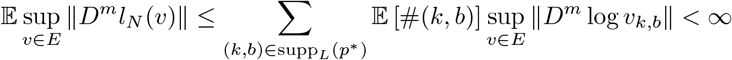

since *E* is relatively compact. Thus, by theorem 1.3.3 of Ghosh and Ramamoorthi [22], 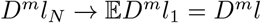 locally uniformly where the last equality is by Leibniz’s rule due to the local boundedness of all derivatives. In particular, *D*^3^*l_N_* are uniformly bounded across *N* on a neighborhood of *υ*\* and, sending 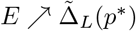, and noting *l_N_* is a.s. eventually −∞ on the boundary of 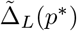, we see *l_N_* → *l* pointwise a.s..

As in the analysis of Dawid [14], we write

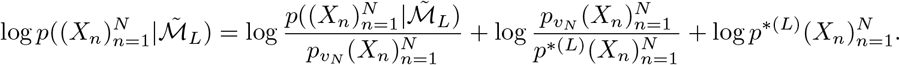

The above paragraph demonstrates that we satisfy conditions (2) of theorem 3.2 of Miller [43] and thus we can write

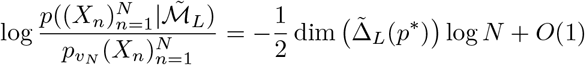

and say that *υ_N_* → *υ*\*. Now, using the mean value theorem,

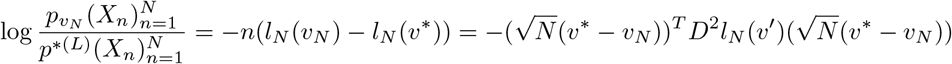

for some 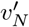 on the ray connecting *υ*\* and *υ_N_*. Call 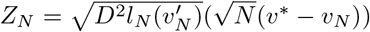. By local uniform convergence, since *υ_N_* → *υ*\*, 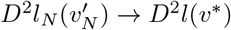. Satisfying the conditions on a neighborhood of *υ*\*, since *υ_N_* → *υ*\*, by theorem 5.41 in van der Vaart [64], 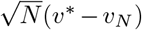 converges in distribution to *N*(0, *D*^2^*l*(*υ*\*)^−1^). Thus, by Slutsky’s theorem, *Z_n_* converges to *N*(0, 1), and by the continuous mapping theorem 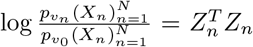 converges in distribution to 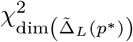; thus this term is *O_P_*(1). Recall from the remark in the last paragraph that 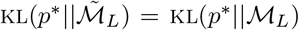 for all *L*; note in particular 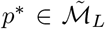 if and only if 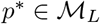. Then finally, by the analysis of Dawid [14], since 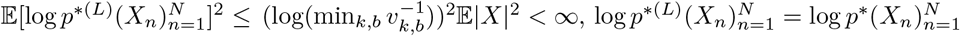 if 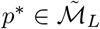 and

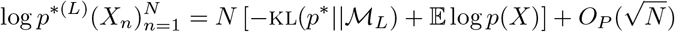

otherwise.

By our analysis above we can say that given *L*_1_ = *L*_2_, if 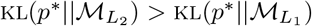,

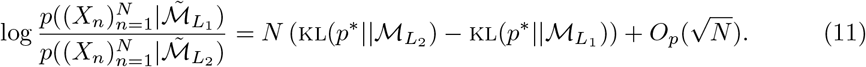

Otherwise, if 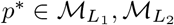,

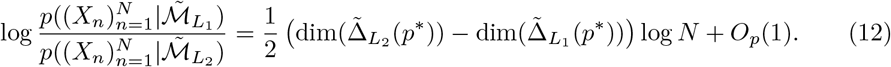

Moving to the second term, for a *k* ∈ supp(*υ*\*), by Stirling’s formula,

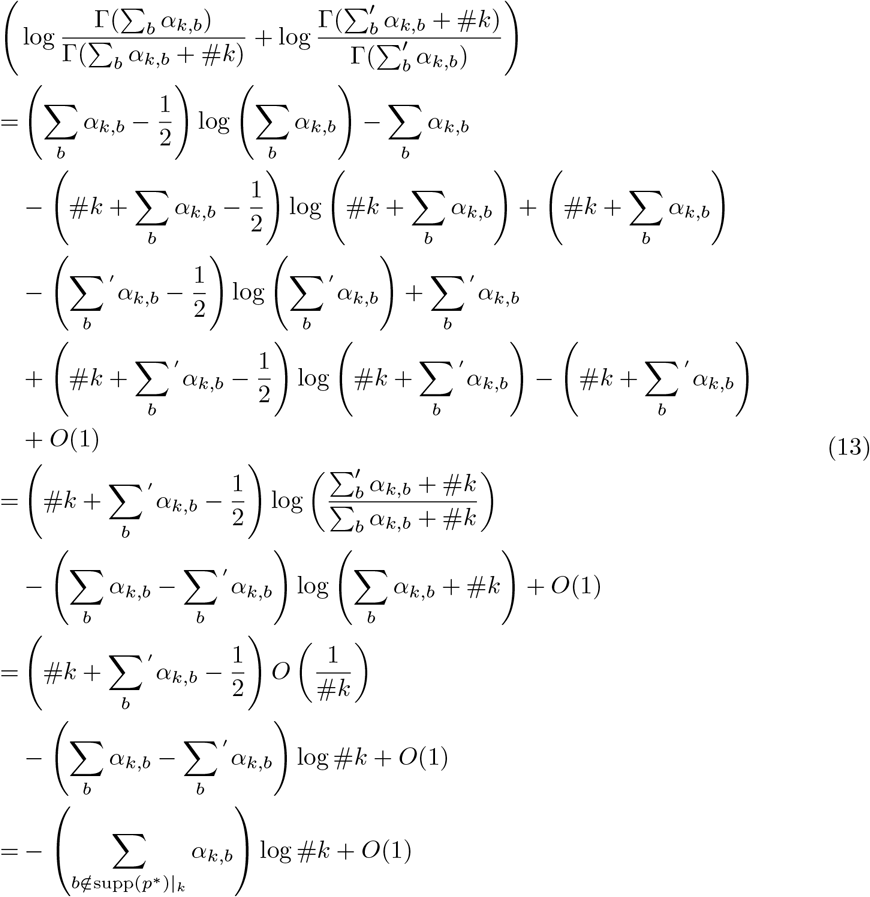

Now note 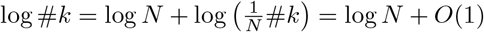 by the strong law of large numbers. Putting this together with 11, 12, 10, and 13 gives the result.

So far, we’ve studied pairwise comparisons between models with different lags; we now study the posterior over lags. We start with the case where there is no true data-generating lag, i.e. 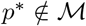. In this case, we can apply theorem 7 to show that the posterior over lags diverges to infinity.

### Corollary 8.

*Let π*(*L*) *denote a prior over lags, with π*(*L*) > 0 *for all L. Choose for each lag a Dirichlet prior on the simplex* 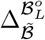 *that satisfies the conditions of Theorem 7. If p* is subexponential but* 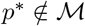, *the posterior diverges in the sense that for any choice of lag* 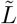, *we have* 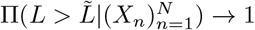 *a.s*..

*Proof*. It is shown in the proof of theorem 23 that as *L* → ∞, we have 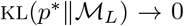. Say 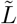 is a lag, so, since 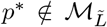, there exists some 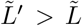 such that 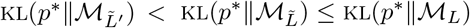 for all 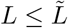. Note we have

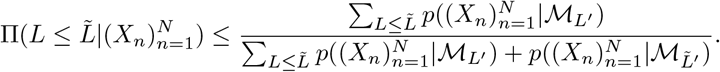

There are only finitely many *L*′ less than or equal to 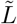, so we can apply theorem 7 and the conclusion follows.

We now consider the case where 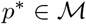. Pick *L*\* to be the minimum lag such that 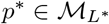. We will need to assume, for theoretical tractability, that the prior over lags has finite support. Then we can establish sufficient conditions for the posterior to concentrate on the true value *L*\*.

### Lemma 9.

*Let π*(*L*) *be a prior over lags with π*(*L*) > 0 *for all L less than some* 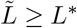, *and with π*(*L*) = 0 *for all* 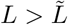. *Then* 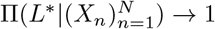 *in probability if* 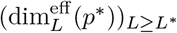 *is non-decreasing and* 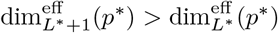.

*Proof*. Apply theorem 7.

If transition probabilities 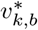 were always non-zero, the effective dimension of the model would simply be the dimension of the parameter space 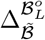, and thus the dimension would always increase with increasing lag, making lag selection consistent. Allowing for 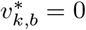 makes the situation more complicated, since in fact the effective dimension may not increase with increasing lag. If this is indeed the case, the posterior will no longer be guaranteed to determine the true *L*\* from data, even asymptotically. In order to describe how the effective dimension in fact scales with the lag, we will introduce the notion of a distribution’s de Bruijn graph: for a distribution *p* on *S*, the *L*-mer de Bruijn graph is the directed graph with nodes acc_*L*_(*p*) and a directed edge connecting *L*-mers *k* → *k*′ if *k*′ = (*k*_2:*L*_, *b*) for a *b* ∈ supp_*L*_(*p*)|_*k*_. (De Bruijn graphs are a common data analysis tool in biological sequence analysis, where they are typically constructed from an empirical distribution over observed sequences; here, we are in effect studying the asymptotic de Bruijn graph, i.e. the de Bruijn graph that we would have if an infinite amount of data were observed.) Call a de Bruijn graph a tree if every node has at most one parent (since sequences must start and end with start and stop symbols, there cannot be a loop where each kmer has just one parent). The next two results show that we can only consistently infer the true lag if the the *L*\*-mer de Bruijn graph of *p*\* is not a tree.

### Proposition 10.

*Say* 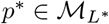 *and for each L, consider a* 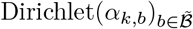 *prior on the simplex in* 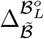 *corresponding to the L-mer k. Say for L ≥ L*, for all L-mers k and bases b*, 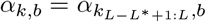 (*i.e. the prior concentration depends only on the last L* letters of the L-mer*). *There exists a* 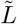 *(possibly infinity) such that for all L ≥ L*, the L-mer de Bruijn graph is a tree if and only if* 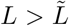. *Then* 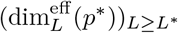 *is a non-decreasing sequence, strictly increasing until* 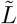, *and constant past* 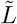.

*Proof*. Call *υ*\* the transition coefficients of *p*\*. Say *L* > *L*\*, *k* ∈ acc_*L*_(*p*\*). Call *k*′ ∈ acc_*L*\*_(*p*\*) the last *L*\* letters of *k*. If for some 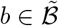, *p*\* (#(*k, b*) > 0) > 0 then clearly *p*\*(#(*k*′, *b*) > 0) thus supp_*L*_(*p*\*)|_*k*_ ⊆ supp_*L*\*_(*p*\*)|_*k′*_. On the other hand, say *b* ∈ supp_*L*\*_(*p*\*)|_*k′*_ = supp(*υ*\*)|_*k′*_ and *Y* is a string, not terminated with $, and with its last *L* characters equal to *k* and *p*\*(*Y*…). 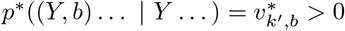 so, *p*\*(#(*k, b*) > 0) > 0. Thus supp_*L*_(*p*\*)|_*k*_ = supp_*L*\*_(*p*\*)|_*k*′_.

Now write

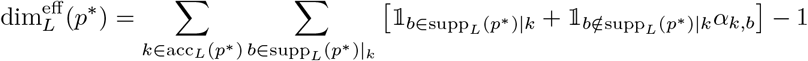

where, for a statement *A*, 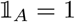 if *A* is true and 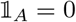 if *A* is false. Thus, since in this case supp_*L*_(*p*\*)|_*k*_ = supp_*L*\*_(*p*\*)|_*k*′_, and by the assumption on the prior coefficients,

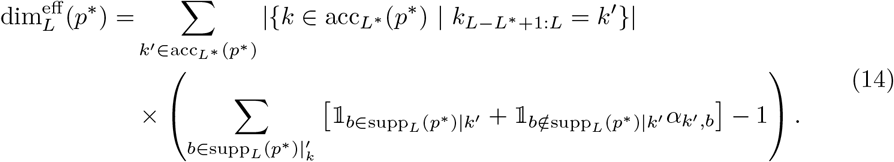

Since for each *k′* ∈ acc_*L*\*_(*p*\*) there is a *k* ∈ acc_*L*_(*p*\*) that has its last *L*\* letters equal to *k*′, 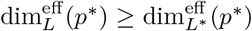. Since 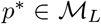 for all *L* ≥ *L*\* the argument may be repeated for all pairs *L*_1_ > *L*_2_ ≥ *L*\* to conclude 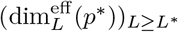 is non-decreasing.

Note if for *L*′ > *L*, 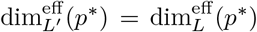 then for all *k*′ ∈ acc_*L*_(*p*\*) there is a unique *k* ∈ acc_*L*′_(*p*\*) with its last *L* letters equal to *k*. Thus if *X*_1_, *X*_2_ ∈ *S* with *p*\*(*X*_1_), *p*\*(*X*_2_) > 0 and *X*_1_, *X*_2_ end in the same last *L* letters (not including $), then *X*_1_, *X*_2_ end in the same last *L*′ letters. Looking at positions |*X_j_*| – *L*: |*X_j_*| – *L*′ + *L* − 1, one can also conclude that *X*_1_, *X*_2_ end in the same last *L*′ + (*L*′ – *L*) letters. Continuing, one may conclude *X*_1_ = *X*_2_. It can be seen that this is equivalent to the *L*-mer de Bruijn of *p*\* being a tree. On the other hand it is not difficult to see that if the *L*-mer de Bruijn of *p*\* is a tree then 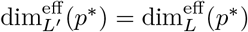 for all *L*′ > *L*.

### Corollary 11.

*Say* 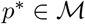 *and L*\* *is the minimum lag such that* 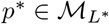. *Let π*(*L*) *be a prior over lags with π*(*L*) > 0 *for all L less than some* 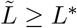, *and with π*(*L*) = 0 *for all* 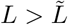. *For each L, consider a* 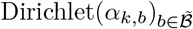 *prior on the simplex in* 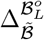 *corresponding to the L-mer k. Assume that for L* ≥ *L*\*, *for all L-mers k and bases b*, 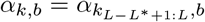. *Then lag selection is consistent if and only if the L*-mer de Bruijn graph of p* is not a tree*.

*Remark 1. If p*\*(*X*) > 0 for infinitely many *X* ∈ *S*, as is the case if the transition coefficients of *p*\* are all positive or there is a cycle in the *L*\*-mer de Bruijn graph of *p*\*, then no *L*-mer de Bruijn graph of *p*\* is a tree as sequences with *p*(*X*) > 0 cannot be identified by their last *L* letters. As another example, pick a particular sequence *X* ∈ *S* and say *X*′ is one letter away from *X*. For a 0 < *q* < 1, define *p* = *qδ_X_* + (1 – *q*)*δ_X′_*. Pick *L*\* the smallest lag such that 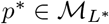. Then the *L*\*-mer de Bruijn graph splits into two paths at the position where *X* and *X*′ differ. These paths may rejoin after *L*\* nodes. Thus the *L*\*-mer de Bruijn graph is a tree if and only if the position at which *X* and *X*′ differ is less than *L*\* letters away from the end symbol $.

## F Misspecification detection

In this section, we turn from studying the parameter *υ* and lag *L* in the BEAR model to studying the hyperparameters *h* and *θ*. Intuitively, we expect the empirical Bayes estimate of *h* to behave as a diagnostic of misspecification, since *h* controls the extent to which the prior predictive distribution of the BEAR model is concentrated at the embedded AR model. Here we make this idea rigorous by examining the asymptotic behavior of the empirical Bayes estimates of *h* and *θ*.

We first briefly introduce the setup and some notation. We will assume *p*\* is subexponential. We will work with fixed lag *L*, though the results can be straightforwardly extended to the case of a prior over a finite number of lags. The function 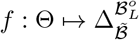 defines an autoregressive model, with parameter space Θ some set. For any *h* > 0, *θ* ∈ Θ, define a prior *π*(·|*h, θ*) on 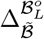 consisting of independent 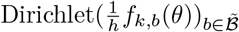 priors on each simplex corresponding to 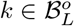. Define 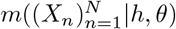 to be the marginal likelihood of the data 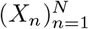 under the prior *π*(·|*h, θ*), that is 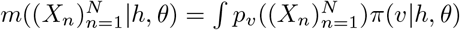. For our purposes we may assume *f_k,b_*(*θ*) > 0 for all (*k, b*) ∈ supp_*L*_(*p*\*); if this is not the case for some *θ* then the marginal likelihood at *θ*, for any choice of *h*, is a.s. eventually 0. We will study maximum marginal likelihood/empirical Bayes estimates 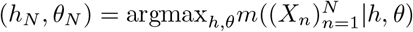.

Our starting point is the analysis of empirical Bayes presented in Petrone et al. [48]. Here is the (very heuristic) intuition behind their result: the Laplace approximation to the marginal likelihood is proportional to the probability of the true data-generating parameter under the prior, so asymptotically we expect 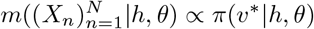. Then, roughly speaking, the empirical Bayes estimate will be (*h_N_, θ_N_*) ≈ argmax_*h, θ*_ *π*(*υ*\*|*h, θ*); in other words, the empirical Bayes estimate should asymptotically maximize the probability of the true parameter parameter value under the prior. Petrone et al. [48] give conditions under which this is indeed true, but BEAR models fail to meet them. There are two major problems: (1) in the limit as *h* → 0, the prior converges to a point mass, making the Laplace approximation invalid (the “degenerate” case mentioned by Petrone et al. [48]) and (2) when some transitions have probability zero, 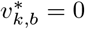, the standard Laplace approximation does not hold regardless of the value of *h*. Our analysis in this section adjusts for both these issues, and also provides more detailed insight such as convergence rates and intuitive approximations for the optimal *h*.

In analyzing extremum estimators, such as the maximum marginal likelihood estimator used in empirical Bayes, uniform convergence results are particularly powerful. Ideally, we might try to establish a Laplace-like approximation to the marginal likelihood that holds uniformly for all *h* and *θ*, but this is unavailable because of the degeneracy at *h* = 0. Our strategy will be to first demonstrate a uniform Laplace approximation over all *h, θ* with some caveats: (1) we ignore transitions that are not possible under *p*\* and analyze their contribution to the likelihood later; (2) if *h* → 0 we assume it does not decrease too fast; and (3) we assume similar control over the prior density at the “true” transition probabilities *υ*\*. In proposition 13 we prove that (3) must indeed hold for when *h_N_, θ_N_* are the maximizers of the marginal likelihood.

For any 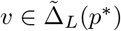, define the negative average log likelihood 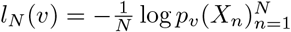, and let 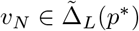 be the (a.s. eventually unique) maximizer of *l_N_*. Define a prior 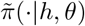 on 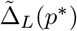 consisting of independent 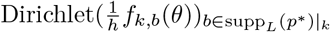 priors on each simplex corresponding to *k* ∈ acc_*L*_(*p*\*) (for a scalar *α*, Dirichlet(*α*) is just defined as the point mass on the 0-dimensional simplex {1}). Let 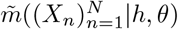 denote the marginal likelihood under the prior 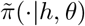 and define

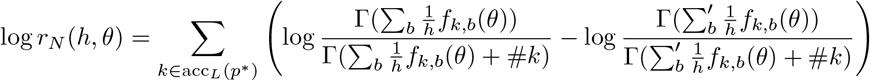

where 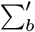 is a sum over the *b* ∈ supp_*L*_(*p*\*)|_*k*_. So, as shown in theorem 7, 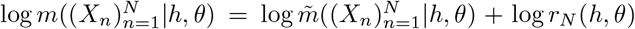. define *B*(*υ, η*) to be the ball of radius *η* around *υ* in some norm; finally, define 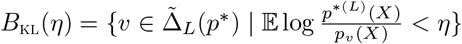 and, for convenience *B*(*η*) = *B*(*υ*, η*), for any *η* > 0.

### Theorem 12.

*With probability 1, for any sequence (h_N_)_N_ and (θ_N_)_N_, possibly dependent on the data, if h_N_N^1/4−ϵ^* → ∞ *for an* 1/4 > *ϵ* > 0 *and* 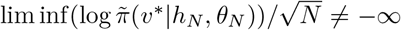, *then*

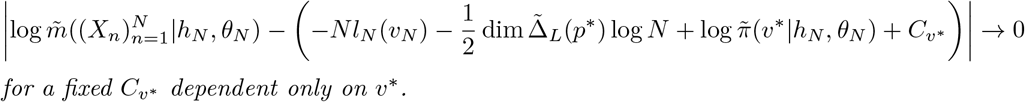

*for a fixed C_v_* dependent only on v*\*.

*Proof*. First note, calling *e_k,b_* the indicator vector at position *k, b* for some *k* ∈ acc_*L*_(*p*\*), *b, b*′ ∈ supp_*L*_(*p*\*)|_*k*_, the directional derivatives with respect to *υ*

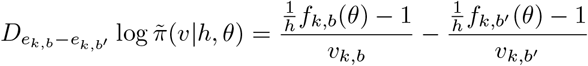

are bounded by *J/h*, for some *J* > 0 in a neighborhood of *υ*\* for all *θ*.

For an *η* > 0, define the KL ball

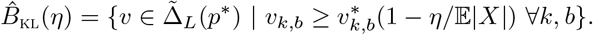

Note if 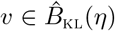, then the KL divergence is bounded,

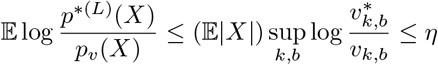

so *υ* ∈ *B*_KL_(*η*). Note

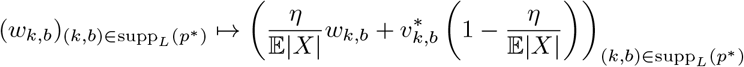

is a diffeomorphism from 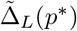 to 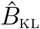 so by the change of variables theorem the volume of 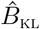 is 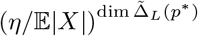 (which comes from the factor multiplying *w_k, b_*) times the volume of 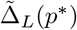. Finally note that by an application of the triangle inequality, 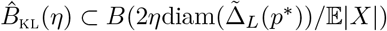.

Define the information matrix at 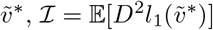, and an *ϵ*′ > 0 less than the smallest eigenvalue of 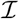 (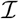 is positive definite by the strict convexity of *l*_0_ described in theorem 7). Also pick an 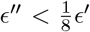 such that 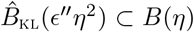 for all small *η*. Now define a sequence *η_N_* = *N*^−(1/4−*ϵ*)^ noting *η_N_/h_N_* → 0. Let 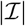 denote the determinant of the information matrix.

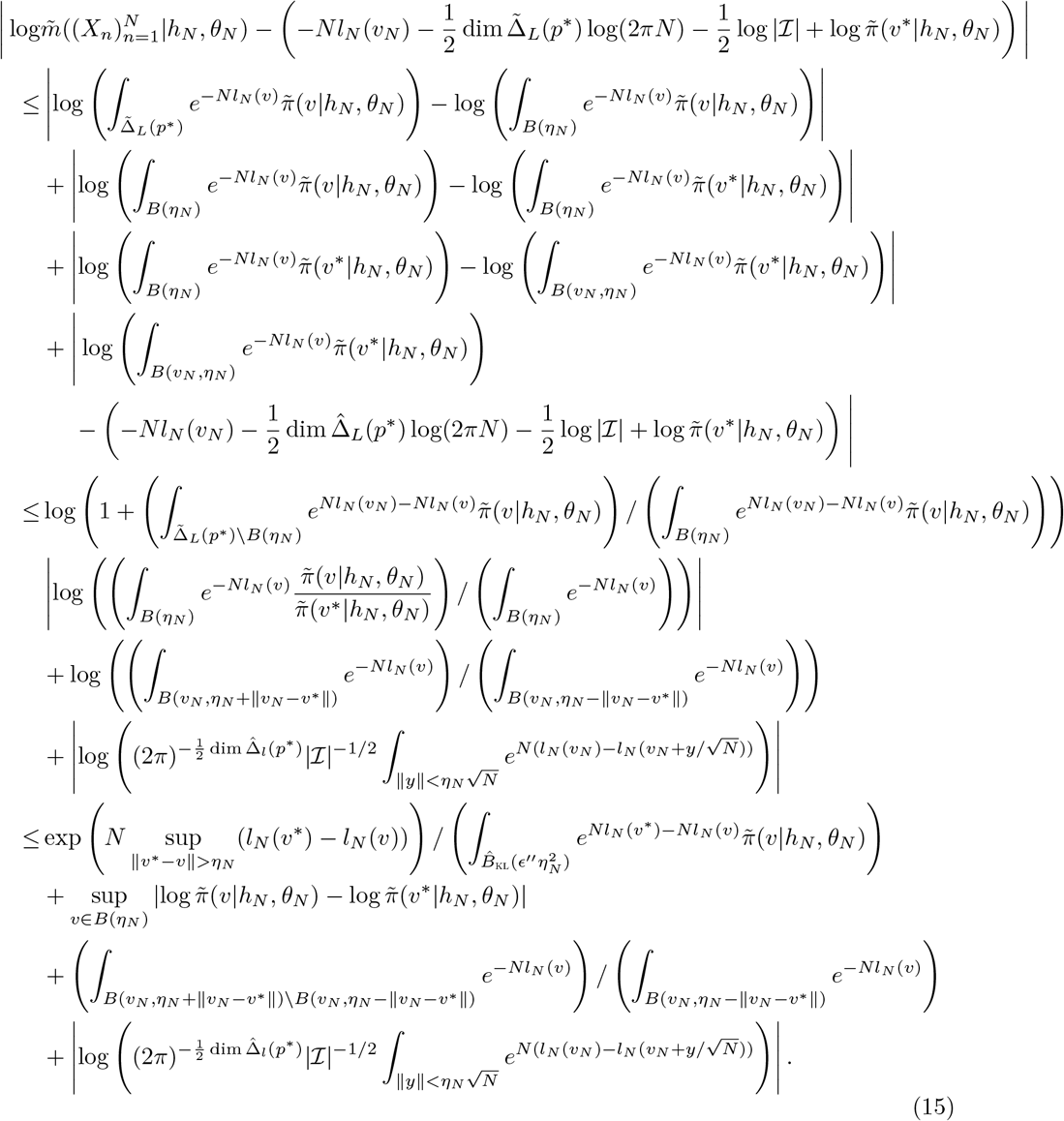

The third line in this inequality follows since *B*(*υ_N_, η_N_* – ||*υ_N_ – υ*\*||) ⊆ *B*(*υ_N_, η_N_*) ∩ *B*(*η_N_*) and *B*(*υ_N_, η_N_*)∪*B*(*η_N_*) ⊆ *B*(*υ_N_, η_N_* + ||*υ_N_* – *υ*\*||). First note that the second term is bounded by *Jη_N_/h_N_* and thus vanishes a.s.. We will show the rest of these terms also vanish a.s..

To analyze the last term, we will use a simplified proof of a Laplace approximation. First note, given the regularity conditions established in the proof of theorem 7, a.s. *υ_N_* → *υ*\*, and *D*^2^*l_N_* → *D*^2^*El_N_* locally uniformly. Thus, for each *y*, since 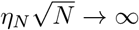, and and *η_N_* → 0 (so that if 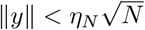 then 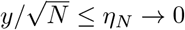), a.s.

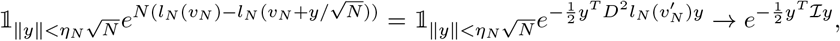

where 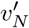 is on a ray connecting *υ_N_* to 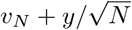. As well, eventually,

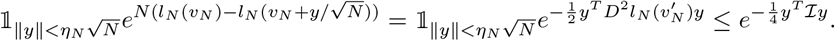

The right hand side is integrable and takes the form of a Gaussian pdf. Thus, integrating the Gaussian pdf, the last term of equation 15 goes to 0 a.s. by the dominated convergence theorem.

To analyze the third term of equation 15, recall from the proof of 7 that *l_N_* is convex, so, the value of −*Nl_n_* is less on the annulus *B*(*υ_N_, η_N_* + || *υ_N_* – *υ*\*||) \ *B*(*υ_N_, η_N_* – || *υ_N_* – *υ*\*||) than on *B*(*υ_N_, η_N_* – || *υ_N_* – *υ*\*||). Thus, to demonstrate that this term vanishes, it suffices to show that ||*υ_N_* – *υ*\*||/*η_N_* → 0 a.s.. Recall from the proof of 7 that we showed that a.s. *υ_N_* → *υ*\* and *D*^2^*l_N_* converges to 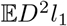 uniformly in a neighborhood of *υ*\*. Thus, eventually, recalling the definition of *ϵ*′ as less than the minimal eigenvalue of 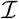, and defining *t* ↦ *υ_t_* as a linear path from *υ_N_* to *υ*\*,

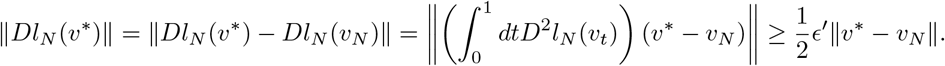

On the other hand, defining *e_k,b_* as above, 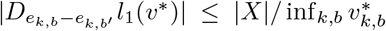 and so, *D_e_k, b_−e_k,b′__l*(*υ*\*) is subexponential. Recalling 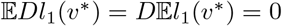, using Bernstein’s inequality (theorem 2.8.1 in Vershynin [65]),

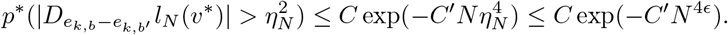

Since 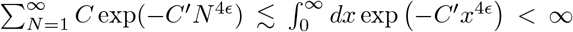, by the Borel-Cantelli lemma, a.s. eventually, 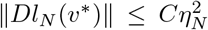 for some *C* > 0. Finally, since *η_N_* → 0, we have ||*υ_N_ – v*\*||/*η_N_* → 0 a.s..

To analyze the first term of equation 15 first note that for small enough *η_N_*, recalling that 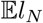 is convex with maximum at *υ*\*, and by the definition of *ϵ*′, we can Taylor expand around *υ*\* and find

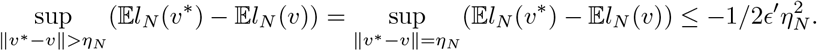

We will also show below that a.s. eventually, for all *υ* away from the boundary (i.e. outside a fixed neighborhood of the boundary), 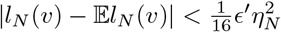. For now, assume that this is the case. So, a.s. eventually, 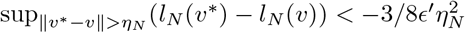, by the triangle inequality. Having bounded the numerator, we now turn to the denominator. Note that by equi-continuity, since *Jη_N_/h_N_* is eventually less than log2, 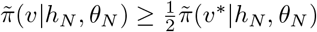 for all *υ* ∈ *B*(*η_N_*). As well, again, by a triangle inequality, a.s. eventually, for all 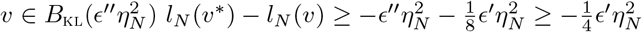. Recall that the volume of 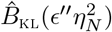 is equal to 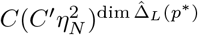 for some *C, C*′ > 0. Then the first term of equation 15 is bounded above by

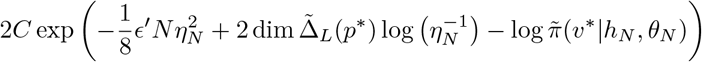

for some *C* > 0. This expression goes to 0 as 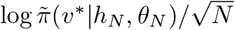 is bounded below and thus 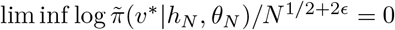.

We now show that a.s. eventually, for all *υ* away from the boundary, 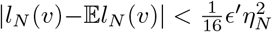. First write

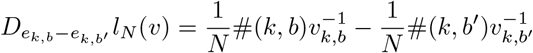

which is almost surely eventually bounded by the strong law of large numbers for all *υ* away from the boundary of 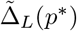. The derivatives of 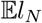 with respect to *υ* are similarly bounded away from the boundary; say the derivatives of both functions are eventually bounded by *J*′. Also note that the random variables |*l*_1_(*υ*)(*X*)| ≤ *C*″|*X*| are uniformly sub-exponential for all *υ* away from the boundary. The covering number of 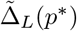 by balls of radius 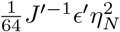 is 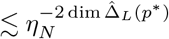. Say (*υ_i_*)_*i*_ are centers of the balls of such a covering. By uniform sub-exponentiality and Bernstein’s inequality (theorem 2.8.1 in ?]), for small enough *η_N_*, 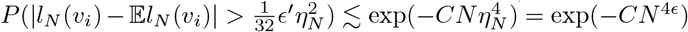 for some *C* > 0. Now, for some *C, C*′ > 0,

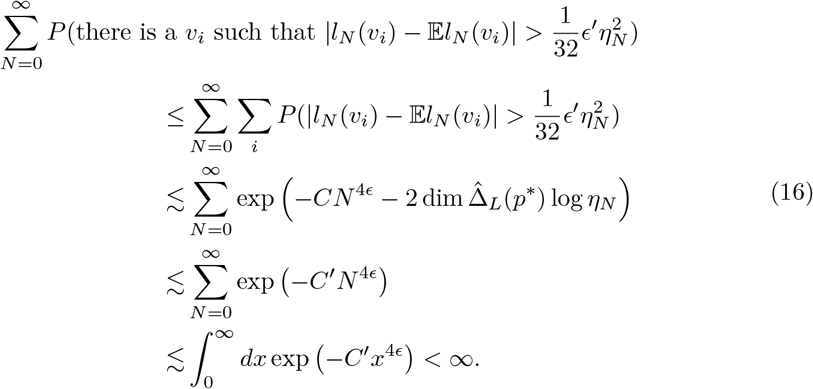

By the Borel-Cantelli lemma, 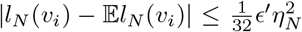 for all *i* a.s. eventually. Thus, eventually, by the triangle inequality and the a.s. eventual boundedness of the derivatives of *l_N_* and 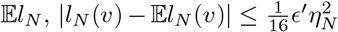 for all *υ* away from the boundary a.s. eventually.

We now focus on the behavior of not just any sequence of *h_N_, θ_N_*, but rather specifically on *h_N_, θ_N_* which maximize the marginal likelihood.^3^ The next two results both use a proof by contradiction strategy that relies on the following logic.

### Remark 2.

Fix *h, θ*. We showed in theorem 7 that log *r_N_*(*h, θ*) = *O*(log *N*) a.s. and we can conclude from theorem 12 that 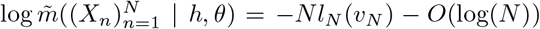. Thus, 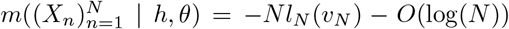. On the other hand, for any *h*′, *θ*′, log*r_N_*(*h*′, *θ*′) ≤ 0 and 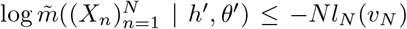. Thus for the maximizers of *m, h_N_, θ_N_*, it is a contradiction if logr_*N*_(*h_N_, θ_N_*) ≲ −*N^β^* or 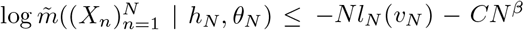 for any *β* > 0: say 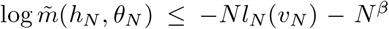. Then, for some *C* > 0, 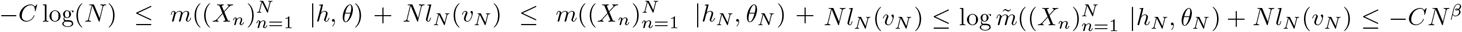, a contradiction. On the other hand, say log*r_N_*(*h_N_, θ_N_*) ≲ −*N^β^*. Then 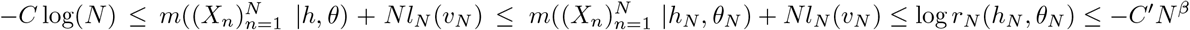, also a contradiction.

### Proposition 13.

*Say (h_N_)_N_ and (θ_N_)_N_ are sequences maximizing* 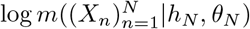 *for each N. Then a.s. there is no subsequence (h_N_j__)_j_ and (θ_N_j__)_j_ such that for some ϵ* > 0, 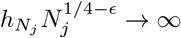 *and for some β* > 0, 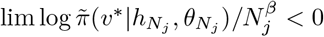.

*Proof*. Assume the opposite. Define (*υ_N_*)_*N*_ and pick (*η_N_*)_*N*_, *ϵ*′ as in theorem 12 such that a.s. eventually, for all *υ* away from the boundary, 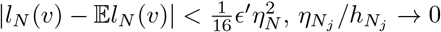, and 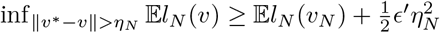. Then, eventually,

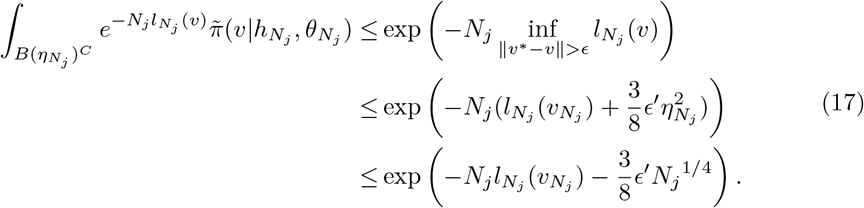

where *B*(*η_N_j__*)^*C*^ denotes the complement of *B*(*η_N_j__*). On the other hand, by equi-continuity of the prior density, since *η_N_j__/h_N_j__* becomes small, for some *C* > 0

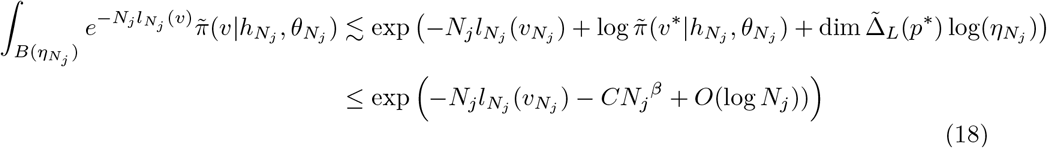

for some *C* > 0. By remark 2, this completes the proof.

We have so far explored what happens to the marginal likelihood when *hN* does not converge quickly to 0, showing that it satisfies a Laplace-like approximation in this case. Next we show that *h_N_* will in fact converge to zero quickly only if the estimated autoregressive model *f*(*θ_N_*) converges to the optimal parameter value *υ*\*.

For a sequence (*θ_N_*)_*N*_ define, for *k* ∈ acc_*L*_(*p*\*), 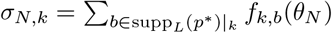 and λ_*N,k*_ = 1 – *σ_N,k_*.

### Proposition 14.

*Say* (*h_N_*)_*N*_ *and* (*θ_N_*)_*N*_ *are sequences maximizing* 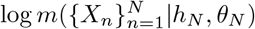. *Then a.s*., 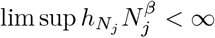 *for some β* > 0 *along a subsequence* (*N_j_*)_*j*_ *only if* 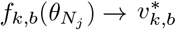 *for all k, b* ∈ supp_*L*_(*p*\*).

*Proof*. Take a subsequence such that: *h_N_j__* → 0; 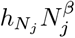 and *h_N_j__ N_j_* both converge, the latter possibly to ∞; *f_k,b_*(*θ_N_j__*) converges for all *k,b*; and *f_k,b_*(*θ_N_j__*)/*h_N_j__* converges, possibly to ∞, for all *k, b*. Note since [0, ∞] is compact, every subsequence with 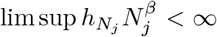 has a further subsequence with these properties. Thus it will be sufficient to show that 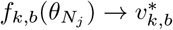 for all *k* ∈ acc_*L*_(*p*\*), 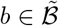. Now define λ_*k*_ = lim λ*_N_j,k__* and *σ_k_* similarly for all *k* ∈ acc_*L*_(*p*\*).

The proof will proceed in two parts. First we will show that if λ_*k*_ ≠ 0 for some *k* ∈ acc(*p*\*), then 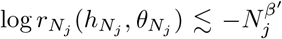 for some *β*′ > 0. This is a contradiction by remark 2 so that λ_*k*_ = 0 and *σ_k_* = 1 for all *k*. Then we will show that if 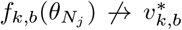 for any *k, b* ∈ supp_*L*_(*p*\*), eventually 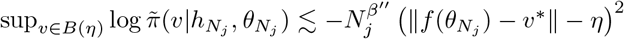. for some *β*″ > 0 for small *η*. Assume this is the case for now. By similar logic to that in equation 17 of proposition 13, for small fixed *η*, it can be seen that for some *β*′″, *C, C*′ > 0,

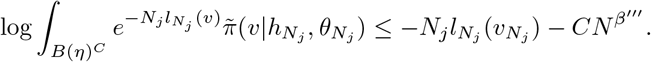

As well,

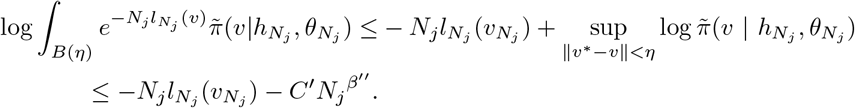

using the fact that 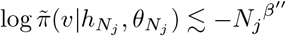. This is also a contradiction by remark 2 and the statement of the theorem follows.

Part one: Assume that for some *k*′, λ_*k*′_ > 0. Performing the Stirling approximation on the terms of log *r_N_j__* depends on the behavior of *σ_N_j_,k_*/*h_N_j__*. Based on the properties of the subsequence we chose, this quantity converges. If it converges to a number greater than or equal to 1 we can perform the usual Stirling approximation with *O*(1) error. On the other hand, if *σ_N_j_,k_*/*h_N_j__* has limit less than 1, using the properties of the Gamma function we write

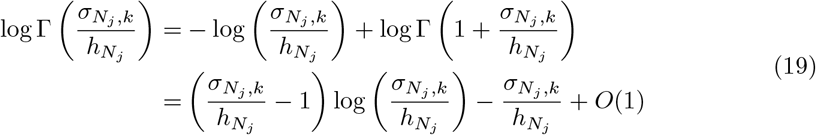

where additional *O*(1) terms were added explicitly in the second line so that the approximation is similar in form to the usual Stirling approximation with the exception of a 1 in the first term instead of 1/2. Define *γ_k_* = 1/2 if the limit of 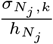 is greater than or equal to 1 and 1 otherwise. Finally recall that *h_N_j__* → 0 and write

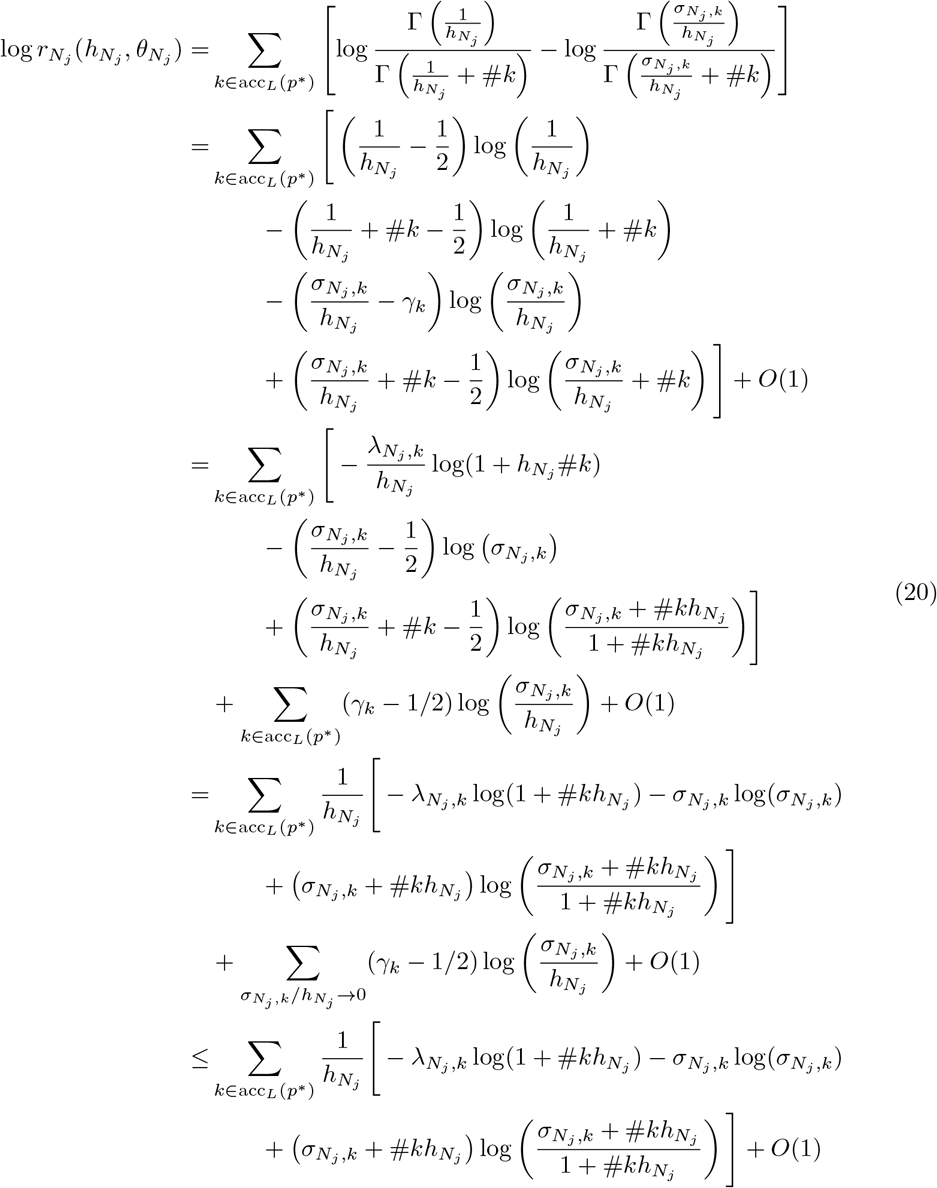

The function

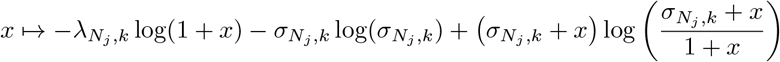

has intercept 0, and derivative 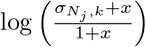, and is thus convex since the derivative is increasing (Fig S2). As *x* → ∞, the function is −λ*_N_j,k__* log*x* + *O*(1) while the function has tangent *x* ↦ *x* log *σ_N_j_,k_* at *x* = 0. In our case, we evaluate at *x* = *h_N_j__ N_j_*, which, based on the chosen subsequence, is either bounded or goes to infinity. First assume *h_N_j__ N_j_* is bounded, say by *M*, and recall that we assumed λ_*k′*_ > 0 for some *k*′, so *σ_k′_* < 1. Then, because the function is decreasing and eventually has negative derivative at 0, we can eventually bound it on [0, *M*] by a line with negative slope and intercept 0 (Fig S2), so eventually, for some *C, C*′ > 0,

**Figure S2:**
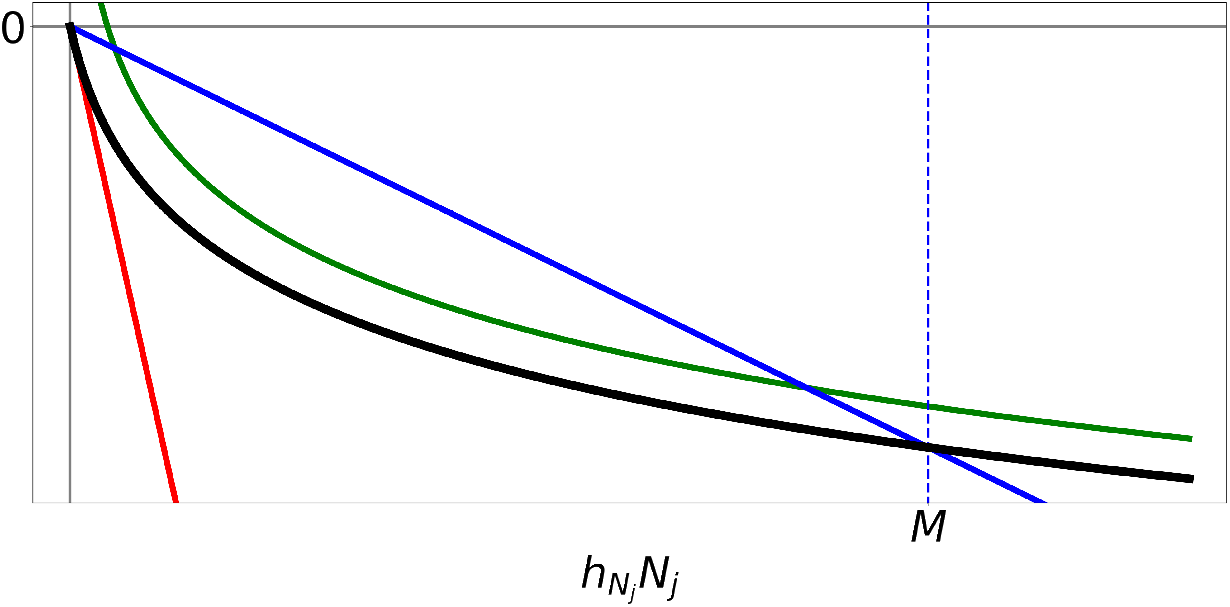
Graph of the function evaluated at *h_N_j__ N_j_* in black when *σ_N_j_, k_* < 1. The red line shows the tangent at 0 with slope log(*σ_N_j_,k_*) < 0. The blue line shows that in this case, where *σ_N_j_,k_* < 1, the function may be dominated by some line for all values less than *M*. The green line shows that as *h_N_j__ N_j_* → ∞, the function is − λ*_N_j_,k_* log(*h_N_j_ N_j__*) + *O*(1).

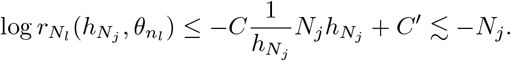

Otherwise *h_N_j__ N_j_* → ∞ so, by the above remark about the limits of the function as *x* → ∞,

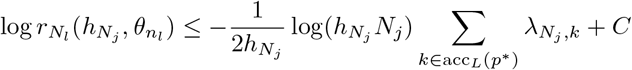

for some *C* > 0 eventually. Recalling that 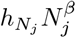 is eventually bounded above, and by assumption log(*h_N_j__ N_j_*) → ∞,

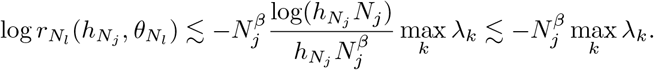

This completes part one of the proof.

Part two: Assume 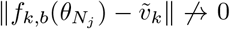. We will perform the same technique to allow a Stirling approximation of the prior: define *Γ_k,b_* = 1/2 if the limit of *f_k,b_*(*θ_N_j__*)/*h_N_j__* is greater than or equal to 1 and 1 otherwise. Then, for all 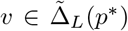 away from the boundary, recalling that we showed in part 1 *σ_N_j_,k_* → 1 for all *k*, if 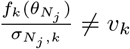 for some *k*,

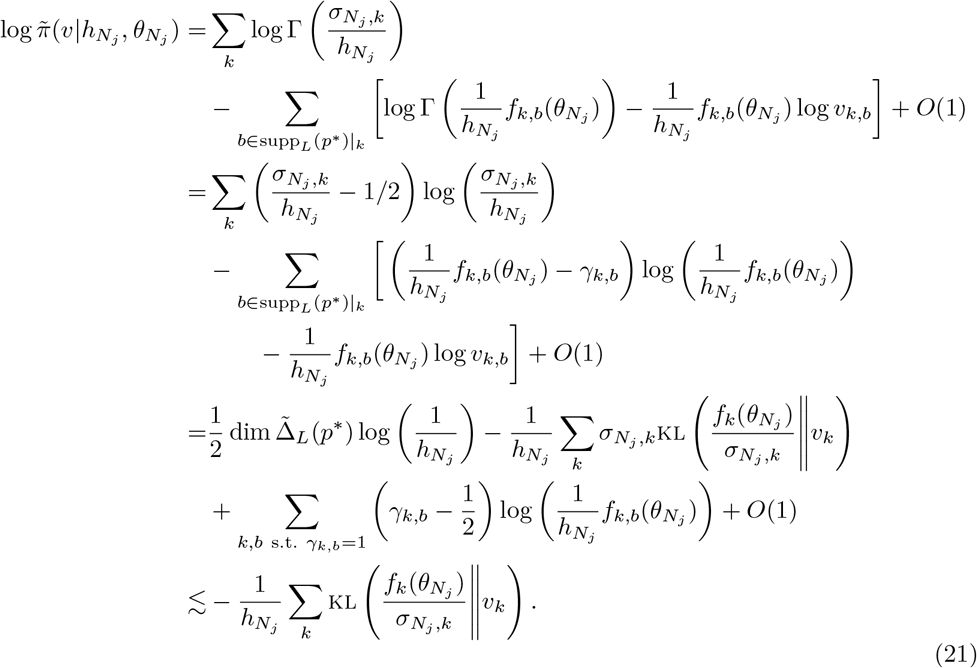

Now note *h_N_j__* < *N^−β^* and for any norm |·|, by Pinsker’s inequality,

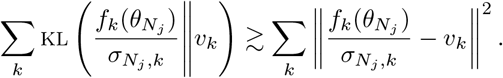

One may check that (Σ_*k*_||·||^2^)^1/2^ is also a norm and *σ_N_j_,k_* → 1 for all *k*, so

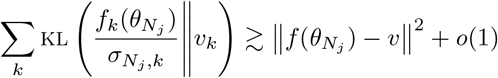

for any norm ||·||. Now note if *η* < ||*f*(*θ_N_j__*) – *υ*\*||,

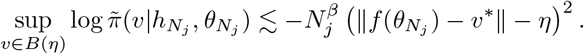

This concludes part two.

We now have the tools to determine the behavior of *h_N_* and *f*(*θ_N_*) in the well and misspecified cases.

### F.1 The well-specified case

We now examine the asymptotic behavior of empirical Bayes inference for the BEAR model in the well-specified case, or, more precisely, when the model is well-specified “at resolution *L*”, in the sense that there are 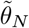 such that for all *k,b* ∈ supp_*L*_(*p*\*), 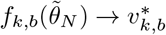 (we say the model is misspecified at resolution *L* otherwise). We first show that the misspecification diagnostic is guaranteed to converge to zero (*h_N_* → 0), correctly indicating that the model is well-specified, and that the embedded AR model converges to the true transition probabilities (*f*(*θ_N_*) → *υ*\*). We also give a bound on the rate for the convergence of *h_N_*, a power of the dataset size. We then establish additional weak conditions under which *θ_N_* also converges to the true value *θ*\*.

#### Proposition 15.

*Say the model is well-specified and* (*h_N_*)_*N*_ *and* (*θ_N_*)_*N*_ *are sequences maximizing* 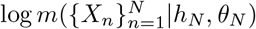. *Then h_N_N^1/4−ϵ^* → 0 *for every ϵ* > 0 *and* 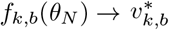 *for all k, b* ∈ supp_*L*_(*p*\*) *with both sequences converging in probability*.

*Proof*. If *U* is a neighborhood of *υ*\* and *β* > 0, proposition 14 shows that

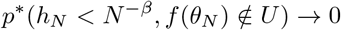

(otherwise *p*\*(*h_N_* < *N^−β^, f*(*θ_N_*) ∉ *U* for infinitely many *N*) > 0). We show below that *p*\*(*h_N_* ≥ *N^−1/4+ϵ^*) → 0 for any *ϵ* > 0 and it will thus follow that we also get *f*(*θ*) → *υ*\* in probability.

Proposition 13 shows that

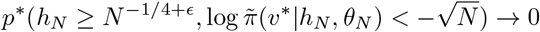

as *h_N_* ≤ *N*^−1/4+*ϵ*^ if and only if *h_N_N^1/4−ϵ/2^* ≥ *N^ϵ/2^*. Thus it is sufficient to show that

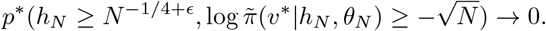

On this set, we may apply theorem 12, but we will need to control 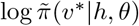.

For any *h, θ*, defining *γ_k,b_* = 1 if 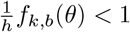 and 1/2 otherwise, and 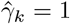 if 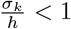 (where recall 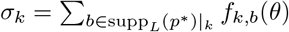) and 1/2 otherwise, by the same derivation as equation 21,

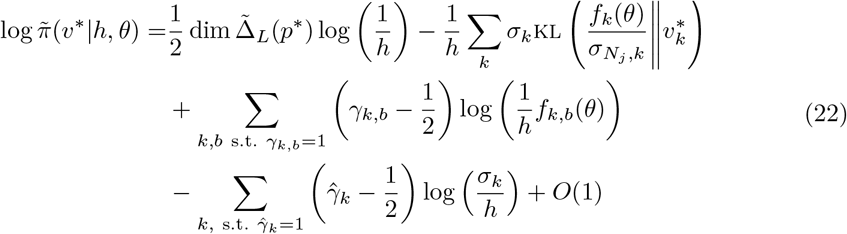

where *O*(1) is uniform over *h* or *θ*. Since 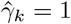 only if *γ_k,b_* = 1 for all *b* ∈ supp_*L*_(*p*\*)|_*k*_, by the concavity of the log function, the sum of these last two terms is negative. Thus,

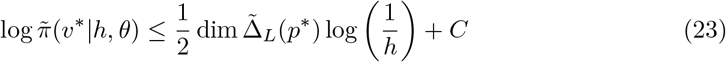

for all *h, θ* for some *C* > 0.

Now we derive a lower bound for 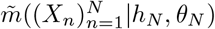. Pick 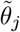 such that for all *k, b* ∈ supp_*L*_(*p*\*), 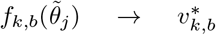. Thus, 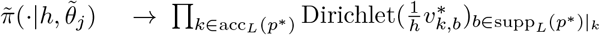 for any *h* > 0 in distribution. And as *h* → 0, we also have 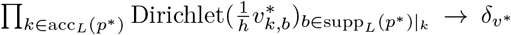. So, pick a sequence 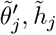 such that 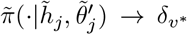 in distribution.^4^ Then 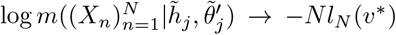. Thus, 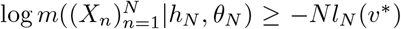. Also recall that from the proof of theorem 7 that, defining *Z_N_* = *Nl_N_*(*υ_n_*) – *Nl_N_*(*υ*\*), *Z_N_* converges in distribution (to a chi-squared distribution). Since log *r_N_* ≤ 0 we can write

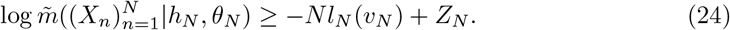

Now, when both *h_N_* ≥ *N^−1/4+ϵ^*, 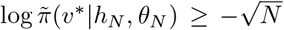, applying theorem 12, we’ve shown that with probability going to 1, for some fixed *C* > 0,

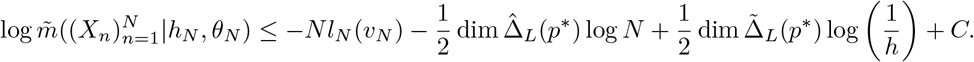

Thus, as *h_N_* ≥ *N*^−1/4+*ϵ*^,

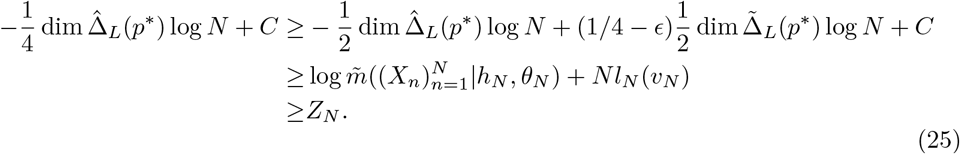

Since *Z_N_* converges in distribution, this occurs with vanishing probability.

We have thus far discussed the asymptotic behavior of *h_N_* and *f*(*θ_N_*). To draw conclusions about *θ_N_* itself, we need to place some assumptions on the autoregressive function *f*. Here we provide an example of such assumptions, drawn from the theory of M-estimators, which say in essence that *f* must have an isolated peak at *θ*\*. These assumptions are enough to guarantee that the empirical Bayes estimate of the AR model parameter *θ* converges to the true value *θ*\*.

#### Corollary 16.

*Say θ*\* ∈ Θ *and d is a metric on* Θ *such that* 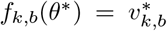 *for all k,b* ∈ supp_*L*_(*p*\*) *and for all δ* > 0,

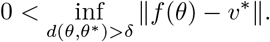

*Then θ_N_* → *θ*\* *in probability*.

*Proof*. Since by proposition 15 we have ||*f*(*θ_N_*) – *υ*\*|| = *OP*(1), we may apply theorem 5.7 of van der Vaart [64] to get the result.

Taking a step back, a perhaps surprising aspect of these results is the weak conditions on *f*. Were we, instead of trying to diagnose misspecification in the AR model, simply trying to analyze uncertainty in the AR model’s parameter estimate, we might proceed by putting a prior on *θ* and performing Bayesian inference for the AR model. In this case, to guarantee asymptotic normality and well-calibrated frequentist coverage, we would in general need strong conditions on *f*, such as bounded third derivatives [43]. Intuitively, the task of diagnosing misspecification might seem to be harder than describing parameter uncertainty, but our conditions on *f* in this section and the next are in fact much weaker, involving no restrictions on the derivatives of *f* whatsoever.

### F.2 The misspecified case

We now consider the case where the AR model is misspecified at resolution *L*. In this case, we can rewrite the marginal likelihood of the BEAR model (using propositions 13 and 14 to apply theorem 12) as

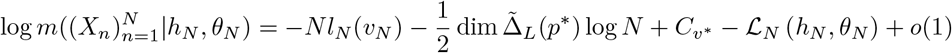

where we define 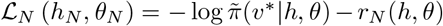.^5^ This expression for the marginal likelihood takes the form of a modified Laplace approximation where, instead of the original prior *π* evaluated at the true parameter value, we have the prior over the support of the data, 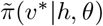, as well as the additional term *r_N_*, which is *O*(log *N*) rather than *O*(1) and depends on the concentration of the prior outside the support of the data. Instead of the standard empirical Bayes behavior described by Petrone et al. [48], wherein the prior probability of the true parameters is maximized, we instead heuristically expect that the objective function 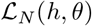 is minimized. The following result makes this intuition formal, showing that *h_N_* and *θ_N_* indeed behavior similarly to the minimizers of 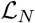.

#### Corollary 17.

*If the model is misspecified at resolution L, a.s*. 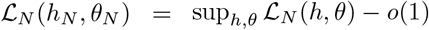.

*Proof*. Say 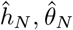 are sequences such that 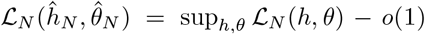. For fixed *h, θ*, we have 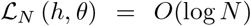. Thus, for any *β* > 0 we clearly have 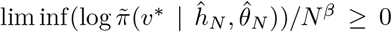 and since we are in the misspecified case, following the logic of proposition 14, equation 20 may be used to see that we also have 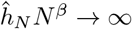. Thus theorem 12 may be applied to 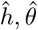 and a comparison of the Laplace approximations of 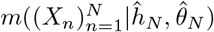 and 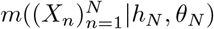 gives the result.

We next examine in greater detail the behavior of the misspecification diagnostic *h_N_*, along with the AR parameter estimate *θ_N_*. There are two cases to consider. First, if the support of the AR model matches the support of the data-generating distribution (that is, supp(*f*(*θ*)) = supp_*L*_(*p*\*) for all *θ*), then *r_N_* = 0 and 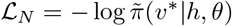; we thus recover the standard empirical Bayes behavior of Petrone et al. [48], with *h_N_* and *θ_N_* asymptotically maximizing the prior probability of the true parameter value. In this case we find that *h_N_* converges to a finite positive value. The second case to consider is when 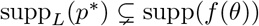. Here, we have *r_N_* ≠ 0, and in particular 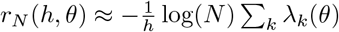. In this case we find that *h_N_* → ∞. Thus, in either case, 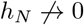, and so *h_N_* will correctly diagnose misspecification in the AR model.

#### Corollary 18.

*If the model is misspecified at resolution L but supp*(*f*(*θ*)) = supp_*L*_(*p*\*) *for all θ, h_N_ is eventually bounded above and below*.

*Proof*. Recall from proposition 14 that if *h* → 0, 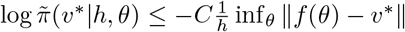 for some *C* > 0. This expression diverges to −∞ as *h* → 0. We also showed in proposition 15 that 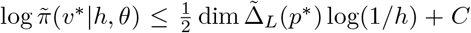 for some *C* > 0. This expression also diverges as *h* → ∞. Combining these two observations along with corollary 17 we get the result.

To say something about *θ_N_*, due to corollary 17, we may use the theory of extremum estimators we can apply theorem 5.7 of van der Vaart [64], replacing limits in probability with a.s. limits to get

#### Corollary 19.

*Say the model is misspecified at resolution L but supp*(*f*(*θ*)) = supp_*L*_(*p*\*) *for all θ. Say also that θ*\* ∈ Θ, *h*\* > 0 *and d is a metric on* Θ *such that for every δ* > 0,

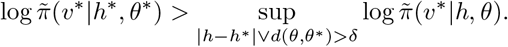

*Then θ_N_* → *θ*\* *and h_N_* → *h*\* *a.s*..

Now we consider the case where the support do not match, i.e. inf_*θ*_ max_*k*_ λ_*k*_(*θ*) > 0, where 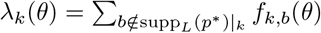.

#### Proposition 20.

*If the model is misspecified at resolution L*, 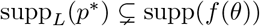 *for all θ, and* inf_*θ*_ max_*k*_ λ_*k*_(*θ*) > 0, *then h_N_* → ∞.

*Proof*. We first show *h_N_* is a.s. bounded below. Since *h_N_N^β^* → ∞ for all *β* > 0, if *h_N_j__* → 0 for some subsequence, we showed in proposition 14 that a.s. 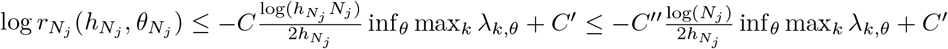 for some *C, C′, C″′* > 0. In particular, log*r_N_j__* ≲ −*O*(log(*N*)) but 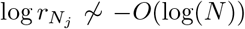 if *h_N_j__* → 0. Thus, since log *r_N_*(*h, θ*) ≥ −*C* log(*N*) for fixed *h, θ*, for some *C* > 0 dependent on *h, θ* and 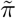 also diverges as *h* → 0, the assumption that *h_N_* maximizes the marginal likelihood is contradicted. Thus, 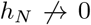. In particular, we showed in proposition 15 (equation 23) that 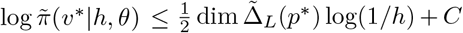 for some *C* > 0 so we get that 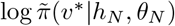 is bounded above a.s..

Assume *h_N_* is bounded above; we will show that this leads to a contradiction. Define *γ_N,k_* = 1/2 if *σ_k_*(*θ_N_*)/*h_N_* ≥ 1 and *γ_N,k_* = 1 otherwise. Define 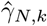 similarly for 1/*h_N_* alone. We next perform the same trick as in proposition 14, expanding 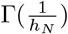 in the form of a Stirling approximation, to analyze *r_N_* further. Noting that log(*h_N_N*) → ∞, we have a.s.,

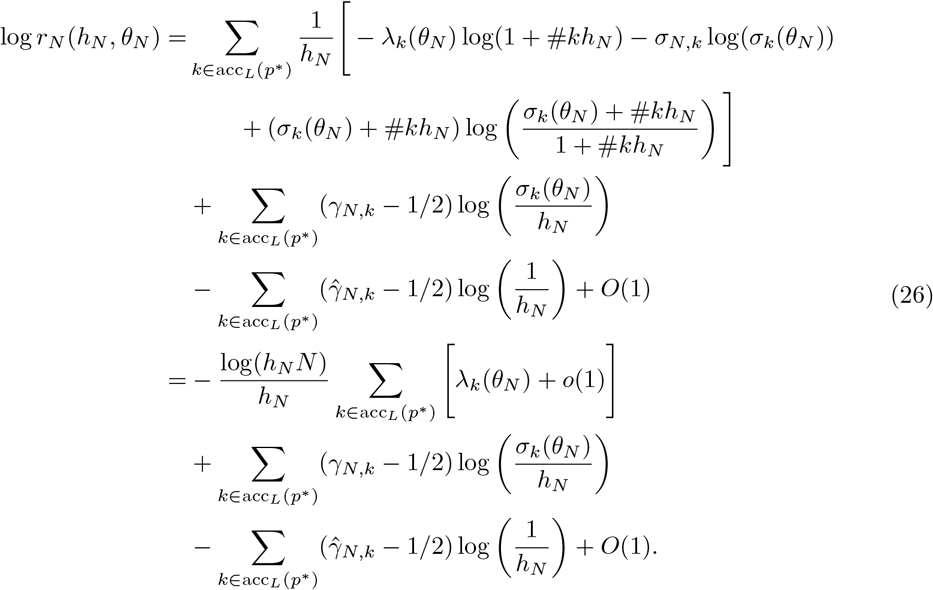

Note 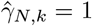 only if *γ_N,k_* = 1 so that the the sum of these last two terms is negative. So, since *h_N_* is bounded above, log *r_N_*(*h_N_, θ_N_*) ≤ −*C*log(*N*) inf_*θ*_ max_*k*_ λ_*k*_(*θ*) for some *C* > 0. Thus, since we also have that 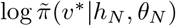 is bounded above a.s., we get that 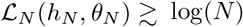 a.s.. On the other hand, with fixed *θ*, if 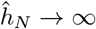 (so we still have log(*h_N_N*) → ∞), then

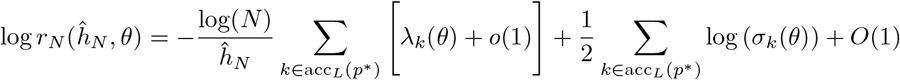

which is −*o*(log *N*), where we wrote 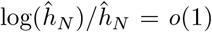. Now pick 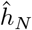 increasing slowly so that 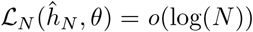. This is eventually less than 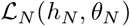, a contradiction. Thus, *h_N_* → ∞.

We can also study the behavior of *θ_N_* in this mismatched supports case, using again the theory extremum estimators. We briefly outline the strategy, omitting details. Further analysis of equations 22 and 26 gives an objective, as *h* → ∞,^6^

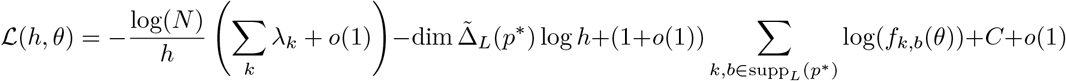

for some fixed *C* > 0. Careful analysis of the *o*(1) terms shows that *h* approaches 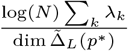. Plugging this value of *h* in, the objective becomes

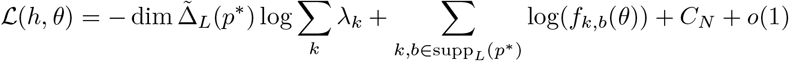

for some constant *C_N_* dependent only on *N* and *p*\*. One can then see that *θ_N_* is an M-estimator of 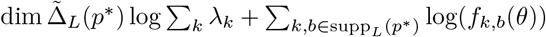 and apply a similar analysis as in corollary 19.

So far we have seen that 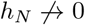 when the AR model is misspecified at resolution *L*, but exactly what value will *h_N_* take and what can it tell us about the amount of misspecification?

Here we analyze the objective 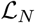 heuristically to address these questions. From the expansions in proposition 14, we can write, for reasonable values of *h, θ*, assuming not too much misspecification,

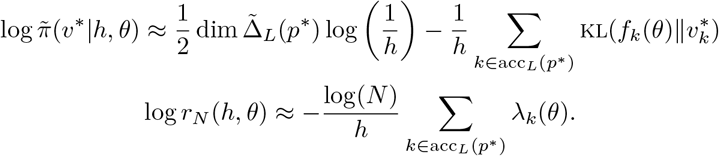

We see, then, that *θ_N_* and *h_N_* depend on an unconventional but valid divergence between the AR model and *p*^*(*L*)^: the sum of the KL divergence between the AR model transition probabilities (from kmers that occur with non-zero probability) and the true transition probabilities, plus a penalty proportional to log(*N*) when the support of the AR model does not match the support of *p*\*. We can thus interpret *h_N_* not only as a diagnostic of misspecification, but also as a measurement of the *amount* of misspecification, and make comparisons between different AR models on the basis of their *h_N_* values.

## G Hypothesis testing

In this section we use the results of the above sections to develop goodness-of-fit and two sample tests.

### G.1 Goodness-of-fit test

Say *p*\* is a distribution on *S* with 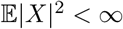 and say *X*_1_, *X*_2_,⋯ ~ *p*\* iid. Say 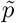 is another distribution on *S* with 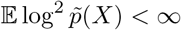 where the expectation is with respect to *p*\*. We are interested in testing whether or not 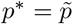, so we will consider the Bayes factor

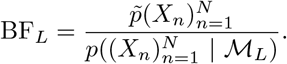

This test asks whether or not 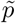 approximates *p*\* at least as well as the optimal model in 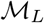. We can use it in particular to test whether 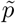 matches the data-generating distribution *p*\* at resolution *L*, that is, whether 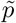 matches *p*^*(*L*)^.

#### Proposition 21.

*Given L, consider a* 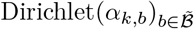 *prior on the simplex in* 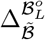 *corresponding to the L-mer k. For all L, assume α_k,b_* > 0 *for* (*k, b*) ∈ *supp_L_*(*p*\*) (otherwise 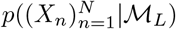 *is eventually* 0 *a.s*.). *Then if* 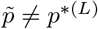,

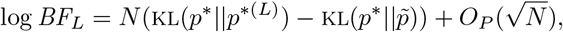

*which goes to* ∞ *in probability if* 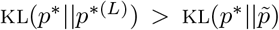 *and to* −∞ *in probability if* 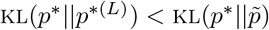. *If* 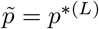

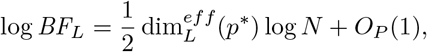

*which goes to* ∞ *in probability*.

*Proof*. Note that as shown in the proof of theorem 7, 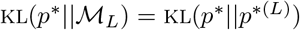, and

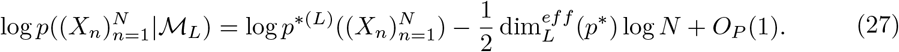

As well, 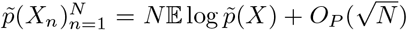 and a similar expression can be written for *p*^*(*L*)^. These two facts prove the result.

#### Remark 3.

One may also consider a Bayes factor that integrates over many *L*:

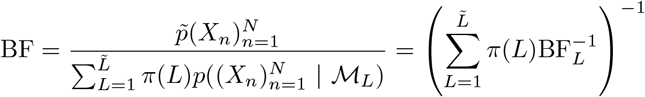

for a prior *π* with 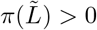. By proposition 21, this Bayes factor goes to 0 if 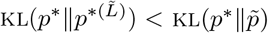 and goes to ∞ if 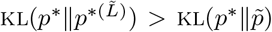 or 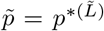 (this later condition is implied by 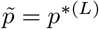 for some 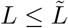 and 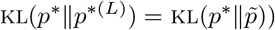. Thus this Bayes factor has the same asymptotics as 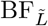.

### G.2 Two-sample test

To set up the two-sample testing problem, consider two distributions *p*_1_ and *p*_2_ on *S* such that 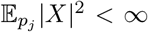 for *j* ∈ {1, 2}. We will assume that the two groups of datapoints are sampled together according to a mixture model with observed labels. That is, let *j*_1_, *j*_2_,… be observed Bernoulli iid random variables indicating the group, with *j_n_* = 1 with probability *β* and *j_n_* = 2 with probability 1 – *β* for a 0 < *β* < 1. Then, let *X_n_* ~ *p_j_n__* independently. The pooled dataset thus follows the generative process *X*_1_, *X*_2_, ⋯ ~ *p*\* = *βp*_1_ + (1 – *β*)*p*_2_ iid. We are interested in whether or not *p*_1_ ≠ *p*_2_. To make this question theoretically tractable, we will fix the lag *L*, and attempt only to discern whether 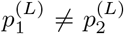 where 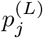 is the best approximation to *p_j_* in 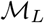 (as defined in section E). In other words, we attempt to distinguish between *p*_1_ and *p*_2_ only up to a “resolution”, in analogy to Holmes et al. [27]. We thus consider the Bayes factor

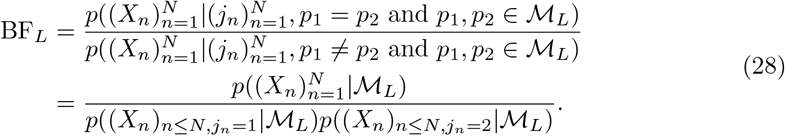

In the subsequent remark, we also extend the theory to Bayes factors that integrate over all *L* up to some fixed maximum.

Consider independent 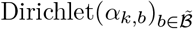 priors on the simplexes in 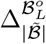 corresponding to the *L*-mers *k*. Assume *α_k,b_* > 0 for (*k, b*) ∈ supp_*L*_(*p*\*) = supp_*L*_(*p*_1_) ∪ supp_*L*_(*p*_2_).

#### Proposition 22.

*If* 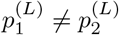,

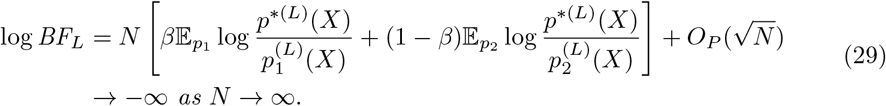

*Otherwise* 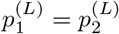 *and*

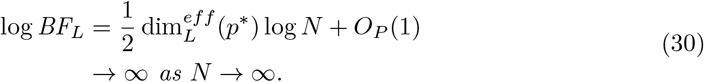

*Proof*. First note that as shown in the proof of theorem 7, noting |{*n*|*j_n_* = *j*}| /*N* = *O_P_*(1),

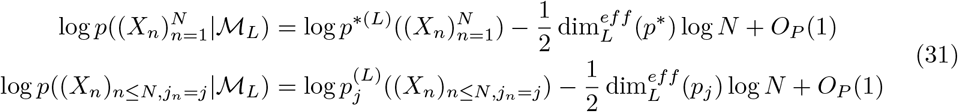

for *j* ∈ {1, 2}. As well, 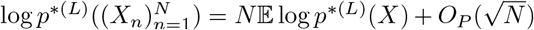 by our assumption on the moments 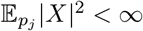 and similar expressions exist for *p*_1_ and *p*_2_. Finally note that

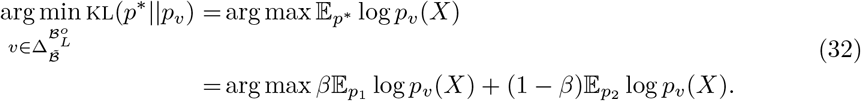

Thus, if 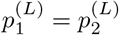 then 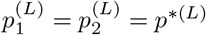.

First assume 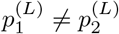. So, we have

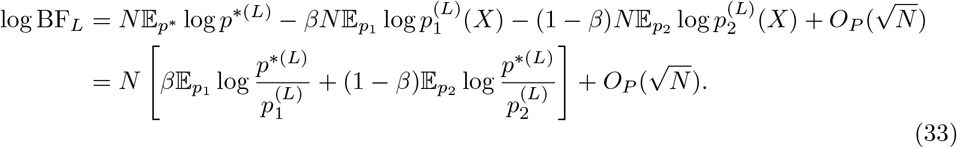

Note 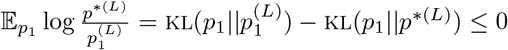 by the definition of 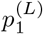. Since 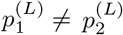, at least one of 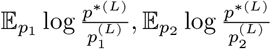 must be negative and so log BF_*L*_ → −∞.

Now say 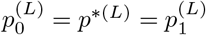. In this case,

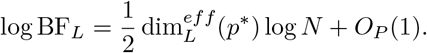

Clearly log BF_*L*_ → ∞.

#### Remark 4.

One may also consider a Bayes factor that integrates over many lags:

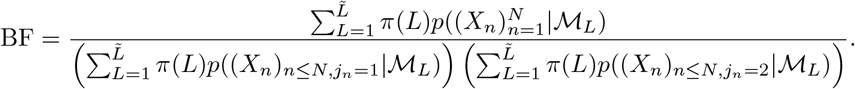

By theorem 7, for all three sums, eventually either (a) assuming the condition for consistency in corollary 11 the term corresponding to the smallest *L* such that 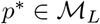 will dominate, if 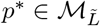, or (b) the term corresponding to 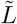 will dominate, if 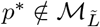. Thus, by analysis similar to that of proposition 22, in any case, we have equation 29 with *L* replaced by 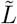, so that the Bayes factor goes to 0 if 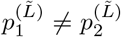. If, on the other hand, we have 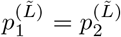, then there are two cases: 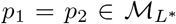 for some 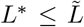 (and *L*\* is picked to be the smallest such lag), or 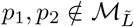. In the first case, 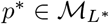 so the asymptotics of BF are identical to that of *BF_L*_* and we can refer to proposition 22 to see that the Bayes factor goes to ∞. In the second case, we may still have 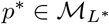 for some minimal 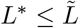; if *p*\* is not a Markov model with 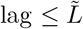, call 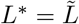. In this case, by the analysis of proposition 22,

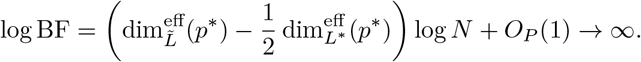

Thus the asymptotics of this integrated Bayes factor are identical to that of 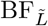.

## H Consistency in the infinite *L* case

So far we have only studied consistency in the finite lag *L* case, that is, our results only show that we can approximate *p*\* up to some finite resolution *L* (corresponding to the largest available lag). In this section, we develop frequentist and Bayesian consistency results for the fully nonparametric model, that is, we allow for priors with support over all lags *L* up to infinity, and show that we can approximate *p*\* itself even if 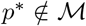. The Bayesian consistency result is our main result, and the most practically useful, but the frequentist result is a natural first step toward the Bayesian result, and an opportunity to develop novel constructions (such as the projection algorithm in section H.2) useful in proving the Bayesian result.

### H.1 Frequentist consistency

We first show that maximum likelihood estimation is consistent, using the method of sieves described in Geman and Hwang [20]. The idea is to increase the size of the model class with the amount of data *N* slowly enough to avoid over-fitting. We define the model class considered for *N* data points first with the lag *L*, but also by restricting transition probabilities to be bounded below by a *ν*: In particular, when there are *N* datapoints, the model class we consider, or the *N*-th “sieve”, is 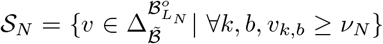 where 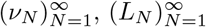 are sequences with *L_N_* → ∞, *ν_N_* → 0.

#### Theorem 23.

*Say X*_1_, *X*_2_, ⋯ ~ *p* iid where p* is a subexponential distribution on S. Say p_νN_ is a maximum likelihood distribution with* 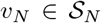 *given* 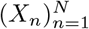. *p_υ_N__* → *p*\* *and* KL(*p*\*||*p_υ_N__*) → 0 *a.s. if for some ϵ* > 0,

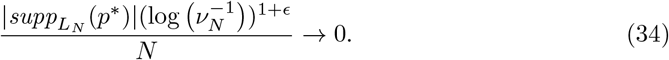

*Proof*. The proof follows that of theorem 3 of Geman and Hwang [20].

First note that 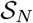 is compact and the likelihood function is continuous so a maximum likelihood *υ_N_* always exists. This satisfies condition C1 of theorem 2 of Geman and Hwang [20].

Next, to satisfy condition C2 (b) of theorem 2 of Geman and Hwang [20] we show that there are 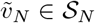 such that 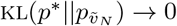. First, for each *L*, pick a distribution *p^L^* on *S* such that for all |*X*| ≤ *L*, *p^L^*(*X*) > 0 and KL(*p*\*||*p^L^*) → 0 as *L* → ∞ (for example, pick *p^L^*(|*X*| > *L*) = *p*\*(|*X*| > *L*), *p^L^*(·||*X*| > *L*) = *p*\*(·||*X*| > *L*) and *p^L^*(·||*X*| ≤ *L*) positive with KL(*p*\*(·||*X*| ≤ *L*)||*p^L^*(·||*X*| ≤ *L*)) < 1/*L*). 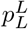 as defined in proposition 3 is a lag *L* Markov model with positive transition probabilities. Thus, for large *N*, its transition probabilities are in 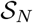. Now notice,

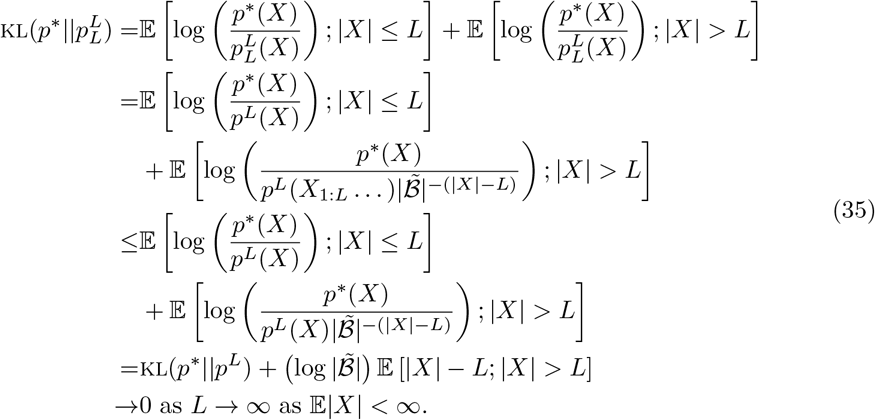

Now we can pick 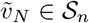 such that 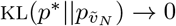.

That KL(*p*\*||*p_N_*) → 0 implies *p_N_* → *p* for distributions *p_N_* on *S* follows from Pinsker’s inequality. This satisfies condition C2 (a) of theorem 2 of Geman and Hwang [20]. However, note that the proof of theorem 2 of Geman and Hwang [20] also shows that if *υ_N_* is an MLE in *S_N_* and the conditions of the theorem hold, then KL(*p*\*||*p_υ_N__*) → 0 a.s..

Finally, we define a partition of each *S_N_* that satisfies conditions i-iii of theorem 2 of Geman and Hwang [20] to get the result. Pick a sequence *ρ_N_* → 0 with 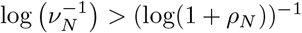 eventually. Call 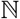 the set of positive integers and for a 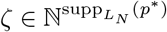, define

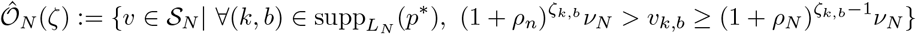

so that 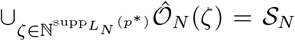 (Fig. S3). Call 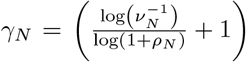 and note (1 + *ρ_N_*)^*γ_N_−1*^ *ν_N_* = 1. Thus the number of choices of *ζ* that give non-empty sets, call this 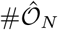, is bounded above by 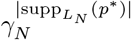. Now notice eventually so that 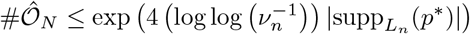.

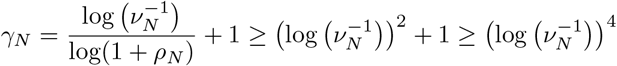

**Figure S3:**
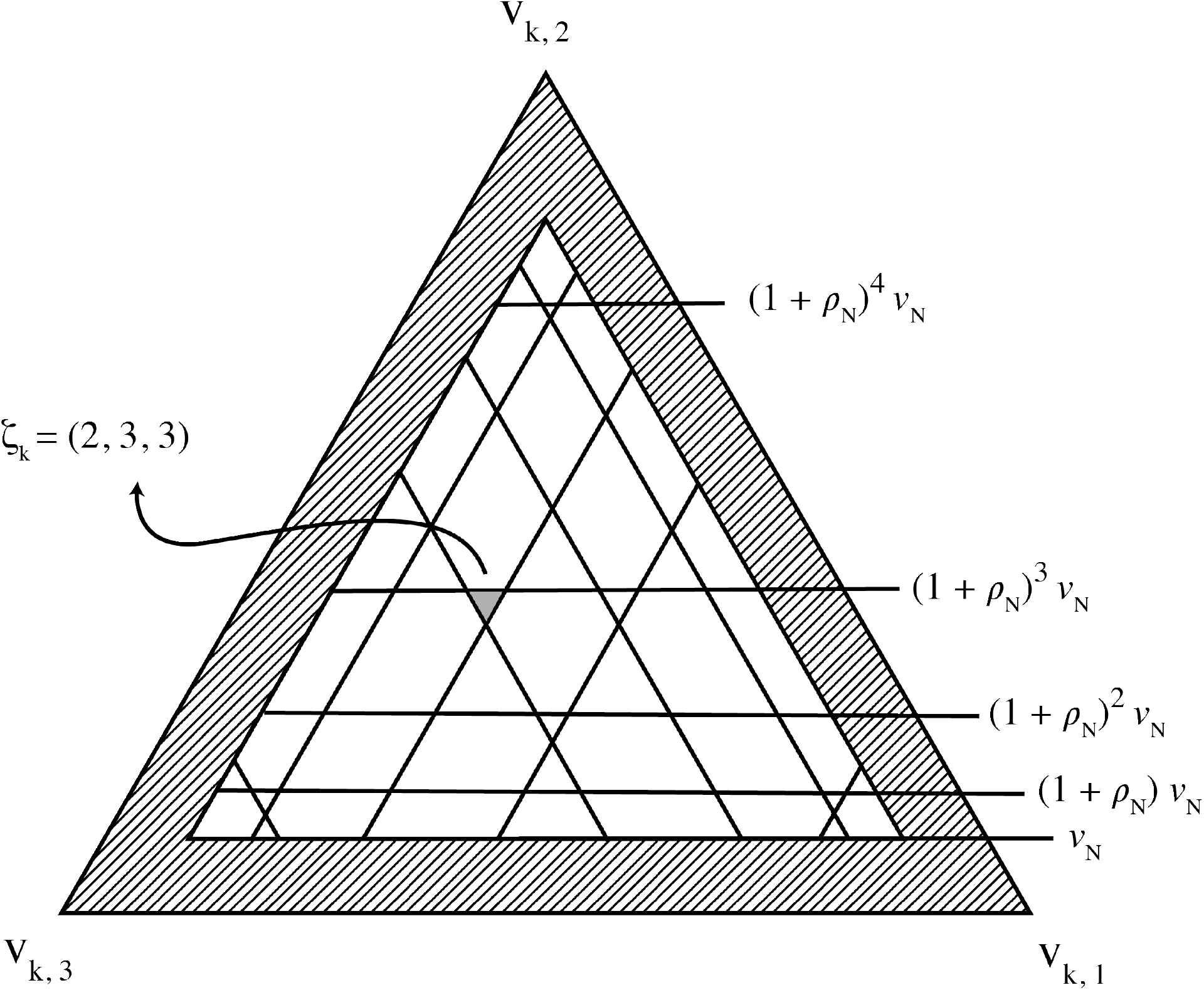
Sieves 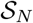 are broken up into subsets 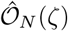, each a Cartesian product of subsets of 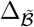, and these subsets in turn are indexed by *ζ_k_* for each *k*. Here we illustrate one such subset of 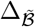, when 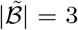 and 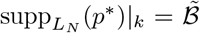. The region included in 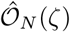 when *ζ_k_* = (2, 3, 3) is shown in solid gray, while all other possible subsets for different values of *ζ_k_* are shown in white. The region adjacent to the border of the simplex (hatched lines) corresponds to those transition vectors that have components less than *ν_N_* and are therefore not part of the sieve 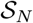.

Say *η* > 0 and, picking a 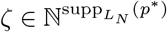, define

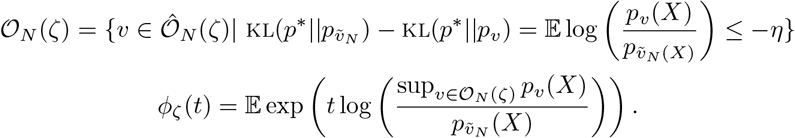

Note 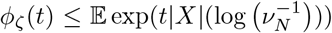 which is finite for small enough *t* by assumption. *ϕ_ζ_* and the bound 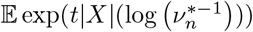 are partition functions for exponential families so, since they are finite for small *t*, they are *C*^∞^ with derivatives obtained by exchanging differentiation and integration for small *t* by theorem 4.5 of van der Vaart [64]. In particular, for 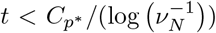 for some *C_p*_* that depends on *p*\*, defining another constant that depends on *p*\*, 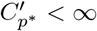,

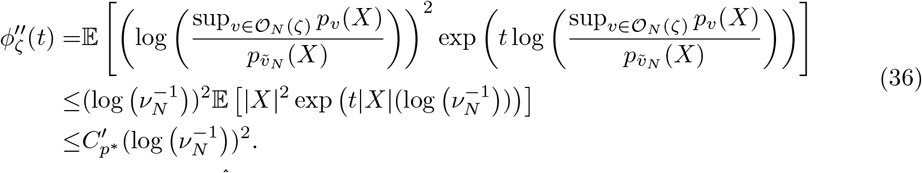

As well, for any 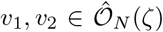, for all (*k,b*) ∈ supp_*L_N_*_(*p*\*), | log (*υ_1,k,b_/υ_2,k,b_*) | < log(1 + *ρ_N_*) < *ρ_N_*. Thus, for all 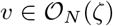, since if *p*\*(*X*) > 0 then all *L_N_*-mer-base transitions in *X* are in supp_*L_N_*_(*p*\*), 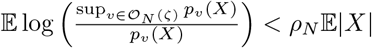. So, defining 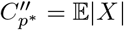,

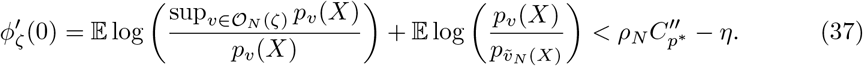

Putting things together we get, for small *t*,

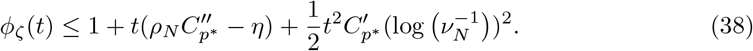

Picking 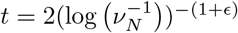 for some *ϵ* > 0 gives, for large enough *N*, for any *ζ*, 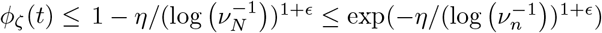. Finally note that

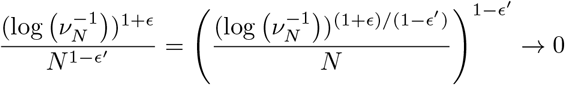

by equation 34 if *ϵ, ϵ*′ are small enough. Now write, for large *N*′ and positive constants *ϵ*″, *C, C*′, *C*″,

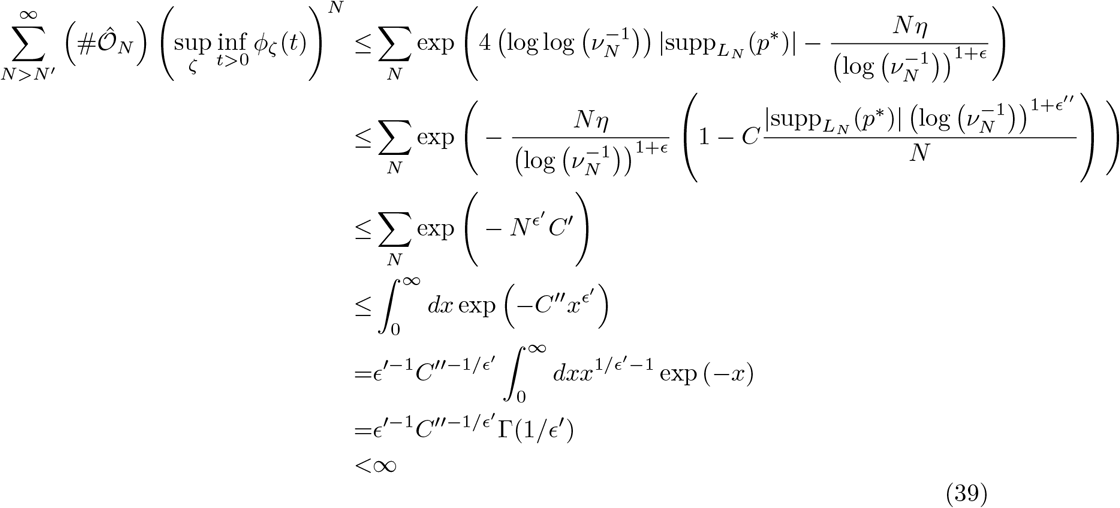

using the assumptions of the theorem and replacing *ϵ* by *ϵ*″ to absorb 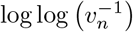 (note one can make *ϵ, ϵ′, ϵ″* as close to 1 as desired). This shows that all conditions of theorem 2 of Geman and Hwang [20] are satisfied.

#### Remark 5.

To pick viable (*L_N_*)_*N*_, (*ν_N_*)_*N*_, note 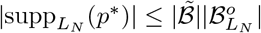, so, since

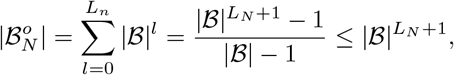

we have 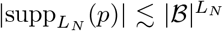. Thus, as an example, for *c*_1_, *c*_2_ > 0 such that 1 > *c*_1_ + *c*_2_, 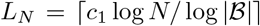 and 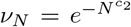 satisfy condition 34. We can see that without any *a priori* knowledge of |supp*_L_N__*(*p*\*)| we are forced to pick a very slow growing sequence (*L_N_*)_*N*_, and thus it is likely that are model class is too conservative for *p*\* whose support have cardinality far from the upper bound. By adapting *L_N_* to the content of the data in addition to its cardinality, the Bayesian approach described in section H.3 does not suffer from this conceptual issue.

### H.2 The projection algorithm

Fix *L* and *ν* for this section and define 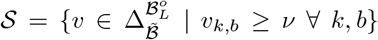. Given data *X*_1_,…, *X_N_*, any maximum likelihood estimate (MLE) in 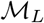, *υ*, has, for every *L*-mer *k* that is seen in the data, 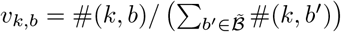 where #(*k, b*) is the number of times *k* is seen in the data immediately preceding *b*. If 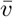 is a MLE in 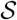, it will be shown that for each *L*-mer *k* that is seen in the data, 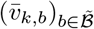 is equal to a “projection” of 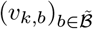 onto the smaller simplex 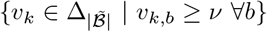. This projection is defined in algorithm 2, and the rest of this section will be devoted to its properties, including continuity, bounds, and proof of the above statement in proposition 28. Some of these bounds will be used to prove the consistency of nonparametric Bayesian inference in section H.3. For ease of exposition, we will first present a conceptually simpler version of the projection algorithm, algorithm 1.

**Algorithm 1.**
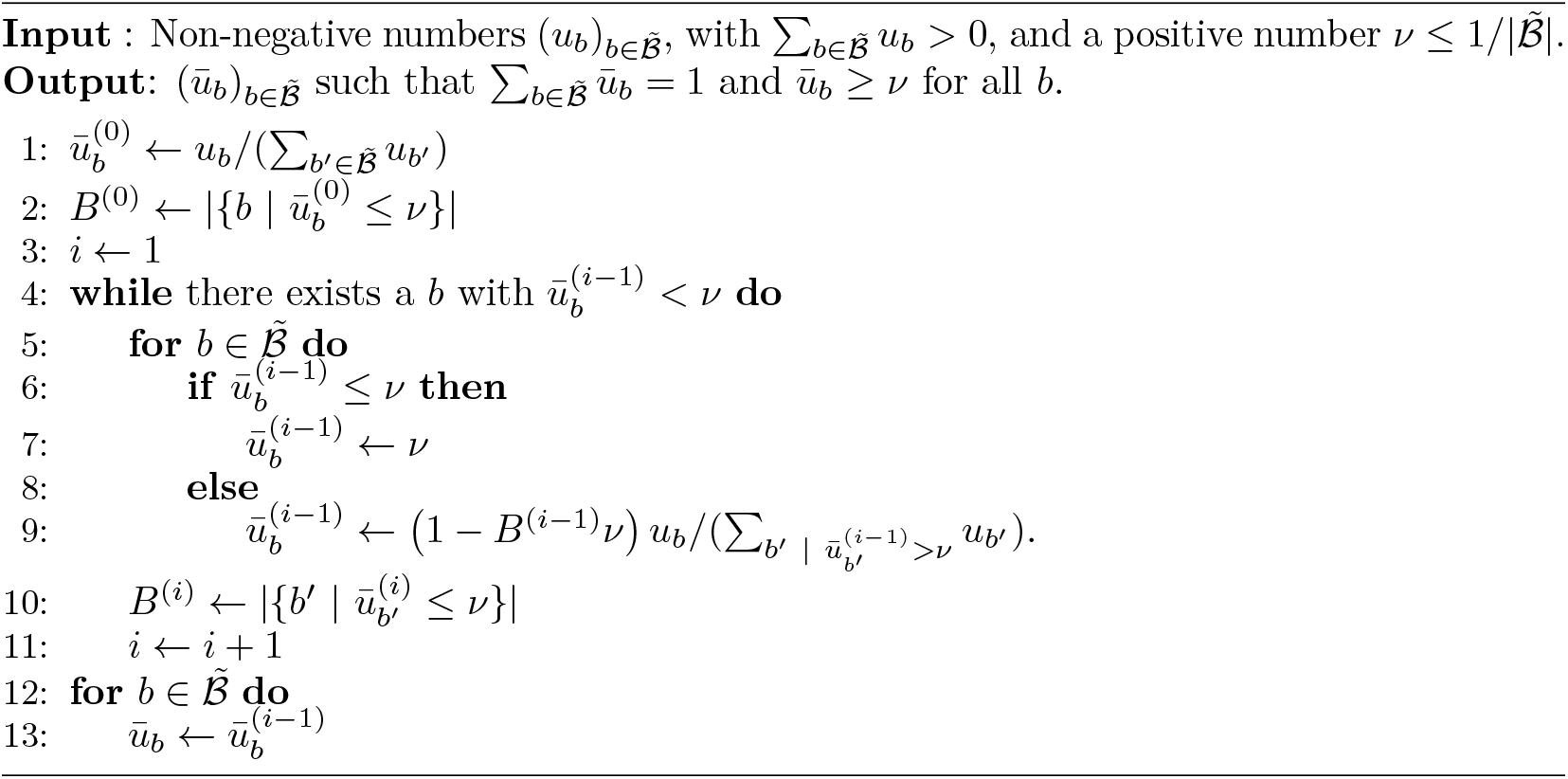
Projection algorithm I

#### Proposition 24.

*Say algorithm 1 is applied to non-negative numbers* 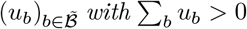. Define 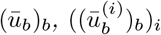 *and* (*B*^(*i*)^)_*i*_ *as in the algorithm. Say the algorithm terminates at step I*.

1. *For all i*, 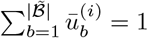.
2. *If* (*u_b_*)_*b*_ *are scaled by a positive constant, the output* 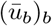 *remains the same*.
3. *Say* 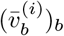 *is the i-th iteration of algorithm 1 with input* 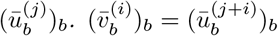.
4. 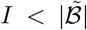. *The algorithm remains unchanged if the while loop were replaced by “**for*** 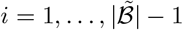 ***do***”.
5. 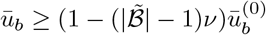.

*Proof*. Results 1 and 2 are clear. For 3, note that if both 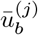 and 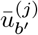 are greater than *ν*, then 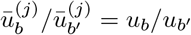. Thus, if 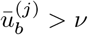,

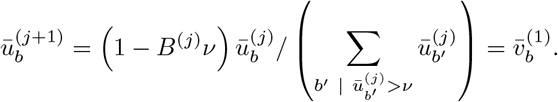

Similar logic may be used to show 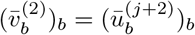 and so on.

To see 4, notice that for every *i* ≤ *I*, at least one *b* has 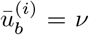 while 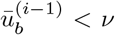. Thus, 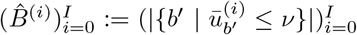 is a strictly increasing sequence. 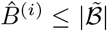 as 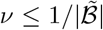. If 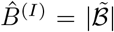 then 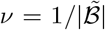 and, by property 1, 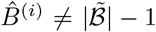 for every *i*. In any case, the sequence 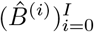 may take on at most 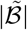 values (including 0) and thus 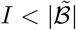. The second statement of 4 follows from the fact that for all *b*, 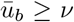 and thus would remain unaltered by the procedure in the while statement.

Finally, for 5, first say 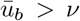 and note that 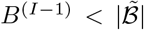 (otherwise the algorithm is terminated or property 1 is violated).

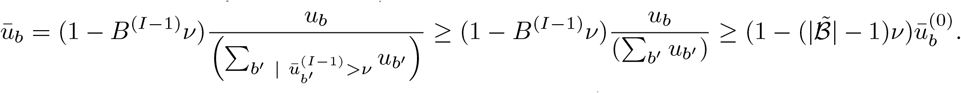

Now say 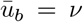. Call *i*′ the first step such that 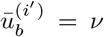. If *i*′ = 0 or *i*′ = 1 then 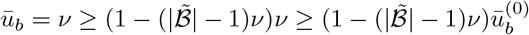. Finally, if *i*′ > 1 then

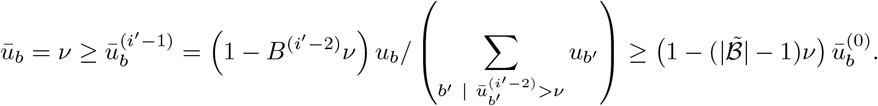

Thus in all cases 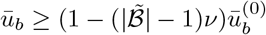.

We now turn to the main projection algorithm.

**Algorithm 2.**
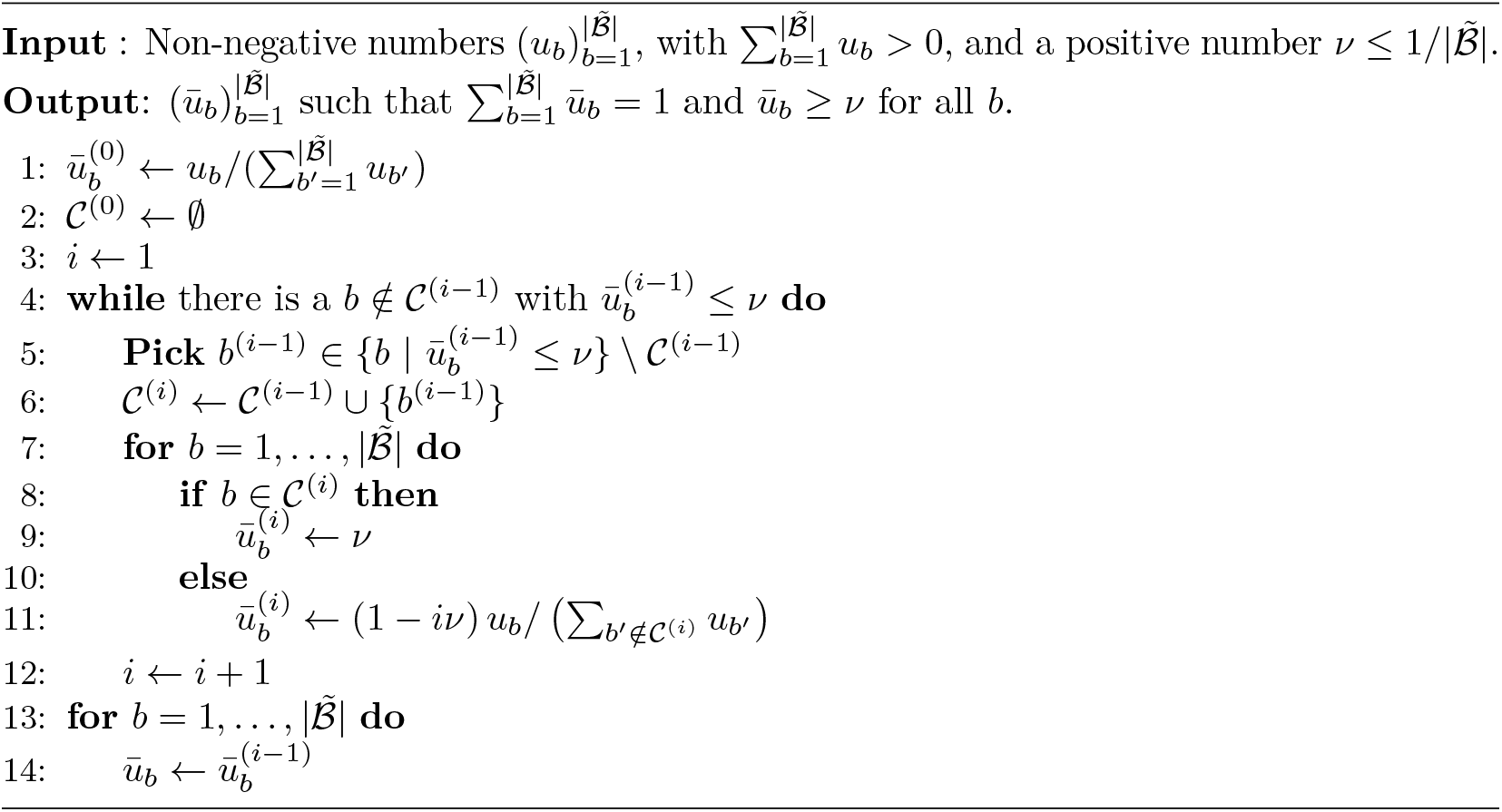
Projection algorithm II

**Figure S4:**
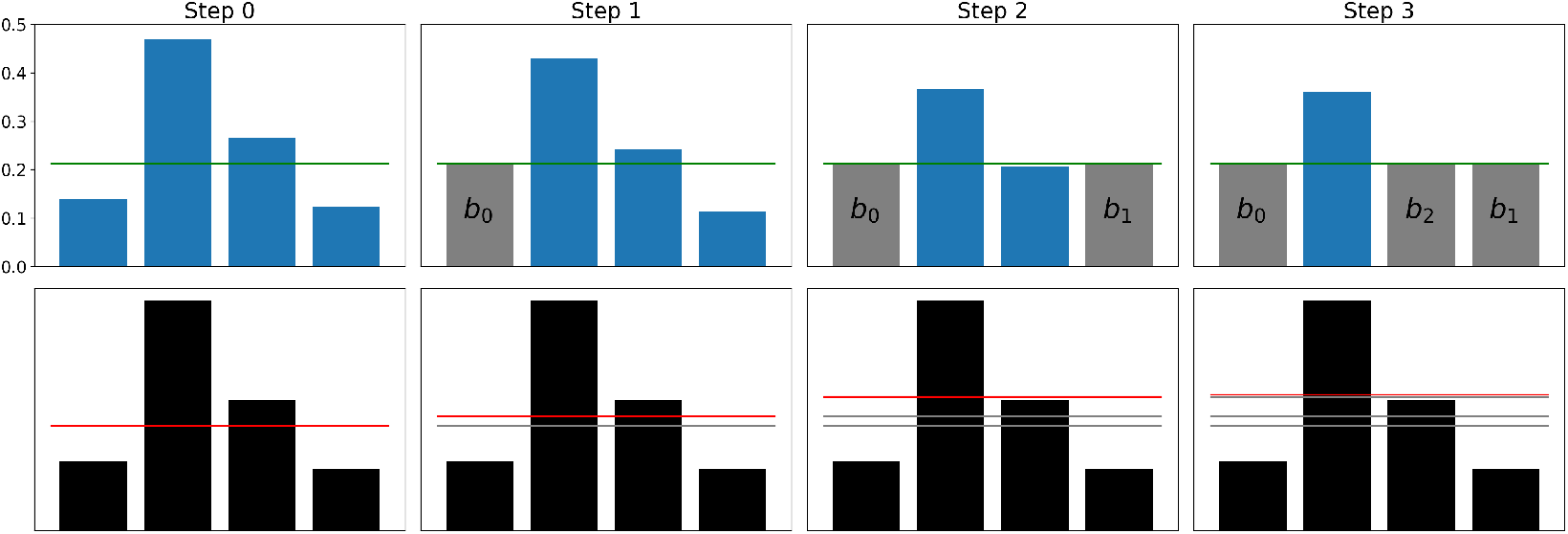
Example application of algorithm 2. 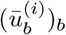 at the end of each step of the algorithm is shown on the top row with *ν* in green and those elements in 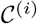 in grey. 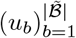 is shown as black bars in the plots in the bottom row with 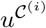 shown as a red line. 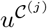 for previous steps *j* < *i* are also shown on the bottom row as grey lines. The scale of the inputs 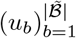 is of no consequence for the algorithm.

An example run of algorithm 2 is visualized in figure S4 (top row). Clearly this algorithm returns 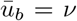 if 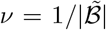 and all the following results are trivial. Thus below we will assume 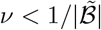.

#### Remark 6.

We will first consider an alternative representation of the algorithm.

Given a 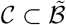, call

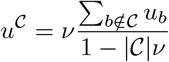

and if 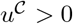, define

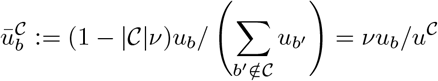

for 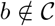 and 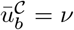 for 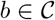; so one gets 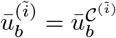 at each iteration 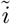. If 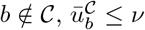 if and only if 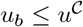.

Say 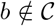 and call 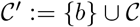.

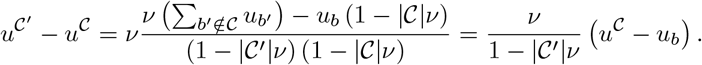

Thus 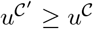 if and only if 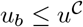 with equality if and only if 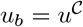.

We can see that at iteration *i* the next *b*^(*i*−1)^ is chosen from 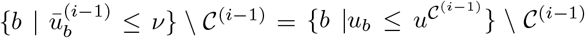, i.e. from those *b* with *u_b_* below the threshold 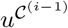. Thus, 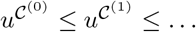. This is reflected in figure S4 (bottom row).

By induction (or from inspection of figure S4), one may show that all the elements *b* of 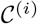 must have *u_b_* below the threshold 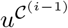 and the algorithm is complete only when all *b* with *u_b_* below the threshold 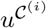 are inside 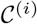. In other words, for *i* < *I* (where *I* is the final iteration) we have 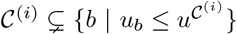, and 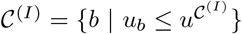.

The important points from the above remark are summarized as:

#### Lemma 25.

1. *Given a* 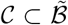, *say* 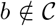 *and call* 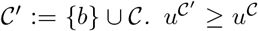 *if and only if* 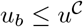 *with equality if and only if* 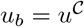.
2. 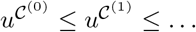.
3. *If the algorithm ends on step I*, 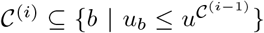 *for all i* ≤ *I*, 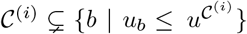 *for i* < *I*, *and* 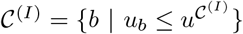.

#### Proposition 26.

*Say algorithm 2 is applied to non-negative numbers* 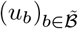 with Σ_*b*_ *u_b_* > 0. *Define* 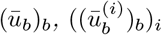 *and* 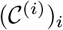 *as in the algorithm. Say the algorithm terminates at step I*.

1. *The output of the algorithm is the same regardless of the choice of* (*b*_0_, *b*_1_,…).
2. *The output of the algorithm is the same as that of algorithm 1*.
3. *we can replace lines 4 and 5 of algorithm 2 with*
4. ***while** there is a* 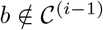 *with* 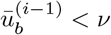 ***do***
5. 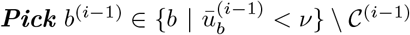 *and receive the same output. With this adjustment*, 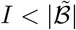.
6. *Say* 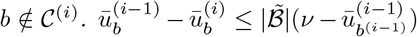 *so that* 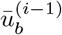 *is close to* 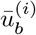 *if* 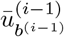 *is close to ν*.

*Proof*.

1. Say the choices (*b*^(0)^,…, *b*^(*I*)^) were made when running the algorithm. Consider a different sequence of choices (*b*^′(0)^,…, *b*′^(*I*′)^) to produce 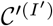. Note that by lemma 25, 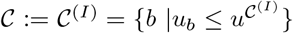 and 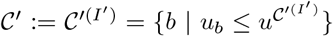. Without loss of generality assume 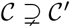 so 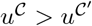. We will show that this leads to a contradiction. Pick the smallest *i* ≤ *I* such that 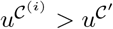. Then 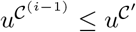, so by lemma 25, 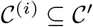. Pick an enumeration 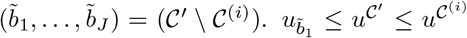 so 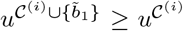. By induction, one may show that 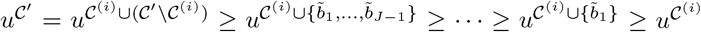. This contradicts the choice of *i* above. Thus, 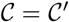 and 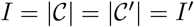. Moreover, since the final output 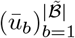 of the algorithm can be defined purely in terms of the final set 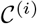, the output must be identical among runs of the algorithm.
2. Consider choosing (*b*^(0)^,…, *b*^(*i*)^) as such: first pick 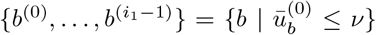, which we know can be done since by lemma 25, 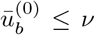 if and only if 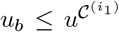 and 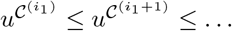. This is equivalent to one step of the while loop of algorithm 1. Then choose 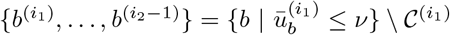, which we can do by similar logic. This is equivalent to the second step of the while loop of algorithm 1. Continuing the construction in the same way, by conclusion (1) above, we get that the outputs of algorithms 1 and 2 are identical.
3. Note, by lemma 25, picking a *b*^(*i*−1)^ with 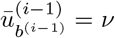 gives 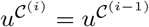 and 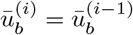 for all *b*. Say (*b*_0_,…, *b_i_*), *i* < *I* are selected in the algorithm such that 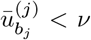 for each *j* ≤ *i* and all 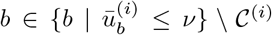 have 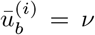, then 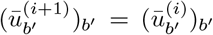 and all 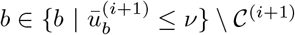 have 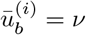. Continuing by induction demonstrates property
4. That 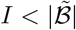 follows by the same logic as conclusion (4) in proposition 24 on algorithm 1.
5. Say 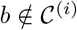,

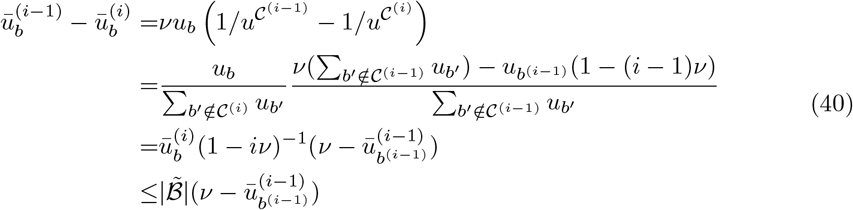

with the last inequality since 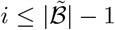 and 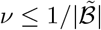.

Next we show that the projection defined by algorithm 26 is continuous.

#### Lemma 27.

*Say* 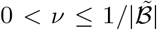 *and* 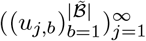 *is a sequence of sets of non-negative numbers, each with at least one positive element, with u_j,b_* → *u_b_ for each b as j* → ∞, *where* 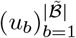 *is set of non-negative numbers with at least one positive element. Apply algorithm 1 or 2 to each set* 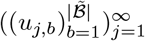 *to get* 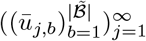 *and to* 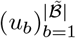 *to get* 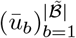. *Then* 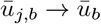 *for all b*.

*Proof*. Define 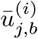 as in the steps of algorithm 2, with *b*^(0)^, *b*^(1)^,… to be defined below. Say 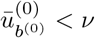. Eventually, 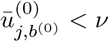 and thus it becomes possible to pick *b*^(0)^ in the first step of the algorithm for all large enough *j*. Then, we get 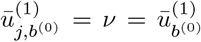. For *b* ≠ *b*^(0)^, 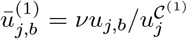 as defined as part of lemma 25. 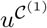 is a continuous function of 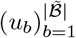 so that 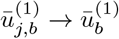 for all *b*. Using the same logic, for large enough *j*, we may pick an *b*^(1)^ with 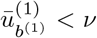 and see 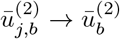 for all *b*. We may continue as such until the algorithm terminates for 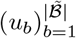 by property (3) in proposition 26. Thus, for some *i*, we have that 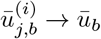 for all *b*.

Note each 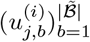 may require another 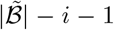 steps for the algorithm to complete. For large enough *j*, we have the implication 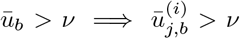 for all *b* so that if for a *b*, 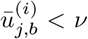, then 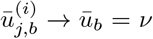. Applying property (4) in proposition 26 to each of the remaining steps of the algorithm applied to 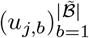 for high enough *j*, considering 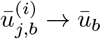 for all *b*, we can see that 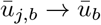 for all *b*.

Finally, we can show that the projection algorithms 1 and 2 indeed return the MLE on the sieve 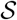, given observed kmer transition counts.

#### Proposition 28.

*Given data X*_1_,…, *X_N_, a lag L, and a positive number* 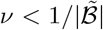, *say* 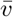 *is an MLE in* 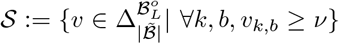. *For every L-mer k that has been seen in the data*, 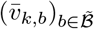 *is equal to the output of algorithm 1 or 2 applied to* 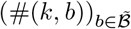 *where* #(*k, b*) *is the number of times k is seen in the data immediately preceding b*.

*Proof*. The likelihood of the data under a 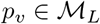 is

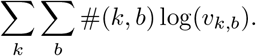

Thus, the MLE in 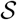 can be found by finding, for each *k* with #*k* > 0,

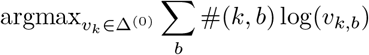

where 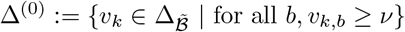.

Say *k* has been seen in the data, so the MLE on 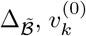, is unique and satisfies 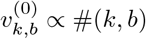. Call 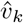 an MLE on Δ^(0)^. Say 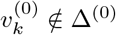 so that for some *b*, 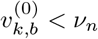. By the uniqueness of the MLE, the likelihood of the data under 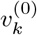 must be strictly greater than under 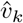. Connecting 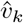 and 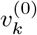 by a line, considering the concavity of the log likelihood function, the likelihood must be decreasing from 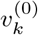 to 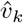. As the likelihood function is analytic and not constant on the line, it must be strictly decreasing. Thus the line cannot intersect Δ^(0)^ except at 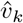. For every *b*, 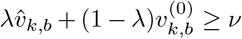 for all λ ∈ [0, 1] if 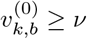; for all λ ∈ [*c*, 1] for a *c* < 1 if 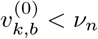 and 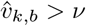; and only for λ = 1 if 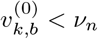 and 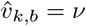. Therefore, for some *b*^(0)^ such that 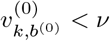 we have 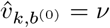.

Call 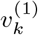 the MLE on 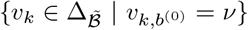. Using Lagrange multipliers again, one may see that

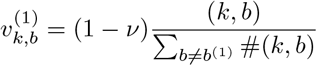

for *b* ≠ *b*^(0)^. Note that 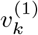 is the result of one step of applying algorithm 2 to 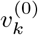 using *b*^(0)^. Call 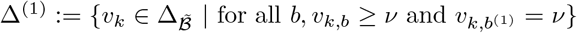 so 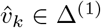. One may perform the same analysis as above to see that if for some *b*, 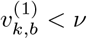, then there is a *b*^(1)^ such that 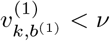 and 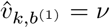.

We may then construct 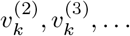 by applying algorithm 2, picking *b*^(*i*)^. Defining Δ^(*i*)^ in analogy to Δ^(0)^ and Δ^(1)^, the algorithm stops at step *i* when 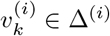 and 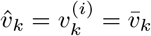. That 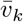 is unique follows from property (2) in remark 26.

### H.3 Bayesian consistency

In this section we take a Bayesian approach to inferring a subexponential *p*\* from data *X*_1_, *X*_2_, ⋯ ~ *p*\* iid. We put a prior on *L*, with support over all *L* > 0, to construct a nonparametric Bayesian model and then study the consistency and concentration rate of its posterior. Recall that the Bernstein von-Mises theorem states that given some regularity conditions, for a Bayesian parametric model, the posterior concentrates in a neighborhood centered at the data-generating distribution, with radius proportional to 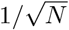. For nonparametric models in general, and (as we shall see) the BEAR model in particular, the concentration rate of the posterior can be strictly slower than 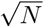 [21, 52].

In order to guarantee consistency and derive a concentration rate, we will, instead of placing a prior directly on *L*, place a prior on sieves constructed similarly to those in section H.1. In particular, define for all *L*, *ν*′ > 0 and *ν* > 0 the sieve

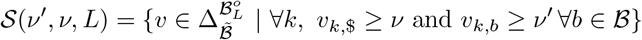

where *ν* is a lower bound on the stop transition probability and *ν*′ is a lower bound on all other transitions. In particular, we will define a prior over the sieves that depends on how well a distribution from each sieve can match *p*\*. Define the sieve approximation mismatch

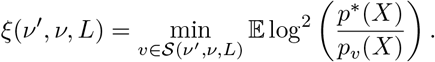

In the next section, we will show that we can guarantee *ξ* is sufficiently small by using the fact that *p*\* is subexponential. Here, we define the prior.

We may now define our prior:

#### Condition 29.

*Assume, for monotonic sequences (ν_m_)_m_, (L_m_)_m_, and a distribution on the natural numbers π*,

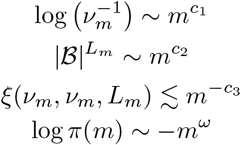

*with c*_1_, *c*_2_, *c*_3_ > 0 *and* 1 > *c*_1_ + *c*_2_. *c*_3_ *must obey the following condition: calling* 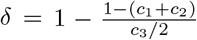, *δ* > 0 *and* (1 – *δ*)^−1^(*c*_1_ + *c*_2_) ≥ *ω* > *c*_1_ + *c*_2_. Consider positive numbers 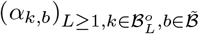 *such that* sup *α_k,b_* < ∞ *and* inf *α_k,b_* > 0. *Consider a prior* Π *on the disjoint union* 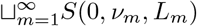 *that factorizes as such:*

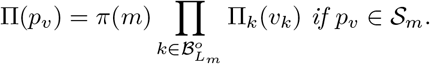

*where for a* 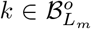, Π_*k*_ *is a restricted and renormalized* 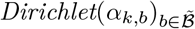 *prior on the simplex in* 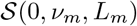 *corresponding to transition coefficients out of k*.

Note as well the difference between the sieve we approximate *p*\* with (*S*(*ν_m_, ν_m_, L_m_*)) and the one our prior is defined over (*S*(0, *ν_m_, L_m_*)). It is best to consider the constraints on *c*_1_, *c*_2_, *ω* with the fact that *c*_3_ is limited in the values it may take on by how well *p*\* can be approximated by finite lag Markov models. Our main result will be the consistency of the posterior under this prior and the calculation of its concentration rate.

#### Remark 7.

Using the techniques in section H.2, we can see that the maximum *a posteriori* estimate on each sieve 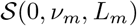 has, for every *k* that has been seen in the data,

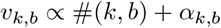

if 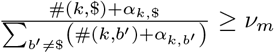; otherwise, *υ*_*k*,$_ = *ν_m_* but we still have *υ_k,b_* ∝ #(*k, b*) + *α_k,b_* for 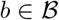. One may then compare the densities of the maximum *a posteriori* estimators in each sieve across *L* to get the maximum *a posteriori* estimator of the entire posterior.

We now discuss two interpretations of this prior. On the one hand, 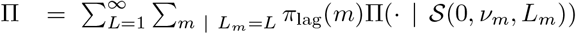 and thus, since 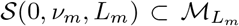, and the fact that multiple *m* correspond to the same *L_m_*, the prior can be interpreted as similar to putting a prior on the lag, with the standard Dirichlet priors on each 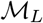, but with the prior having a “staircase” shape for very small stopping probabilities. On the other hand, we have carefully chosen the values of *ν_m_* and *L_m_* in order to balance the size of 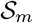 against the amount of information about *p*\* received from *m* datapoints. How this works will become clear in the proof of theorem 35.

In section H.3.1 we will show that there exists a *c*_3_ such that *ξ*(*ν_m_, ν_m_, L_m_*) ≲ *m*^−*c*_3_^, i.e., *p*\* may be efficiently approximated by the sieves. Then we will derive our main result with the concentration rate in section H.3.2. Finally we describe how to use this result in practice on real data in section H.3.3. Throughout we will consider a data generating distribution *p*\* and all expectations will be with respect to the data generating distribution unless otherwise stated.

#### H.3.1 Approximating subexponential sequence distributions

In this section we will be interested in finding an asymptotic upper bound for *ξ*(*ν_m_, ν_m_, L_m_*) of the form *m*^−*c*_3_^, thus showing that a prior as in Condition 29 exists (proposition 32). The result relies on the assumption that *p*\* is subexponential; our main consistency result (theorem 35) would only require 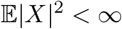 if Condition 29 were somehow otherwise satisfied. In its essence, this section is about constructing approximations to subexponential sequence distributions, with control not only over the expected log ratio of *p*\* and the approximating distribution *p* – the KL divergence, 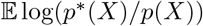 – but also over the variance of this log ratio – i.e. control of 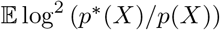. We will make use of lemma 3 but need another construction and technical lemma.

Note that if *p*\* is a distribution on *S* and *X* ∈ *S*,

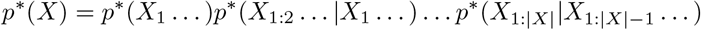

where, recall, for a sequence *Y*, possibly not terminated by $, *p*\*(*Y*…) = *p*\*({*X* ∈ *S* | *X_i_* = *Y_i_* ∀*i* ≤ |*Y*|}). Thus a probability distribution on *S* may be described by its infinite-lag transition probabilities *p*\*((*Y, b*)… |*Y*…) for sequences *Y* not terminated by $ and 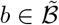, ignoring those *Y* with *p*\*(*Y*…) = 0. Infinite-lag transition probabilities were considered in the construction of 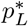 in proposition 3. Below we will be interested in constructing another distribution from *p* by projecting, for some *L*, the transition probabilities at each *Y* with |*Y*| < *L* onto 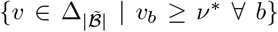. This first lemma will be used to guarantee the existence of this distribution.

**Figure S5:**
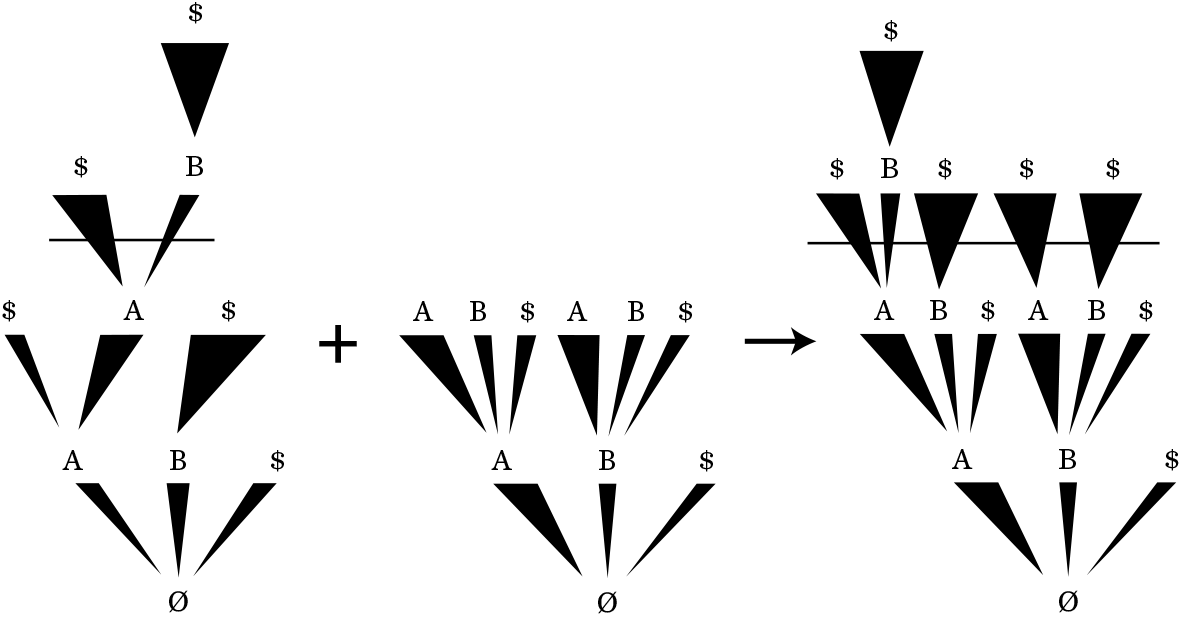
Example application of this construction to the distribution *p*\* on the left, with the *υ* represented in the center. Transition probabilities for kmers smaller than *L* = 2 are those defined by *υ* while those after are those of the original distribution. Thickness of lines denote probability of particular transition.

##### Lemma 30.

*Say p* is a probability distribution on S. Given a lag L and positive numbers* 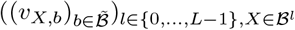 *with* 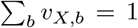 *for all X, there is a p^*L^ such that for all sequences Y not terminated by* $,

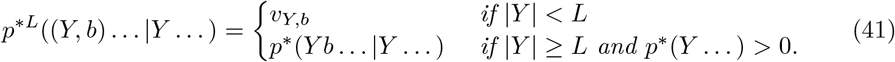

*Proof*. For *X* ∈ *S*, |*X*| ≤ *L* define

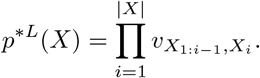

For 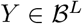 with *p*(*Y*…) = 0, define

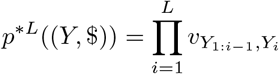

and *p^*L^*(*X*) = 0 for *X* ∈ *S* with *X*_1:*L*_ = *Y* and *X*_*L*+1_ ≠ $. Finally, if *p*\*(*Y*…) > 0 define, for all *X* ∈ *S* with *X*_1_ ⋯ *X_L_* = *Y*,

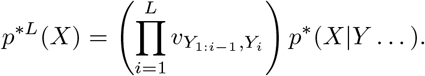

It is not difficult to check that *p^*L^* is well defined and satisfies the requirements in the statement (Fig. S5).

Finally, we write a technical lemma:

##### Lemma 31.

*There exists a positive constant C such that for any p* and p that are distributions over S*,

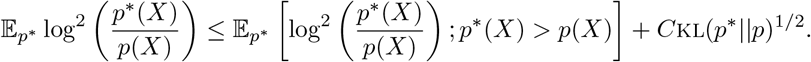

*Proof*, *x* ↦ (log *x*)^2^ is differentiable with derivative 2*x*^−1^ log *x*. The derivative is bounded above on [1, ∞), say by *C*. Thus, for all *x* ≥ 1, (log*x*)^2^ ≤ (log 1)^2^ + *C*(*x* – 1) = *C*(*x* – 1). Now,

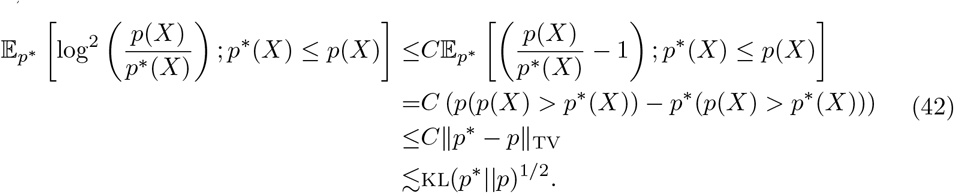

##### Proposition 32.

*If* 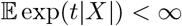 *for some t* > 0 *then* 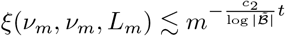.

*Proof*. To approximate *p*\* with a distribution in 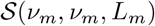 we will use the construction in lemma 3, however we must make sure that the transition probabilities are not less than *ν_m_*. To do so, for sequences *X* without $, with |*X*| < *L_m_*, define 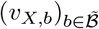 to be the output of the application of algorithm 1 or 2 to 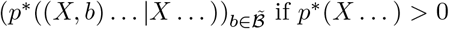. For *X* with *p*\*(*X*…) = 0, make any choice of (*υ_X,b_*)_*b*_ with *υ_X,b_* ≥ *ν_m_* for all *b*. Thus, for all *X, b, υ_X,b_* ≥ *ν_m_*. Now, by lemma 30, there is a distribution *p^*L_m_^* with the same infinite-lag transition probabilities as *p*\* for |*X*| ≥ *L_m_* and infinite-lag transition probabilities 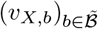 for |*X*| < *L_m_*. Finally perform the construction in lemma 3 to *p^*L_m_^* to produce a 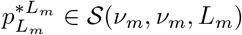.

By lemma 31

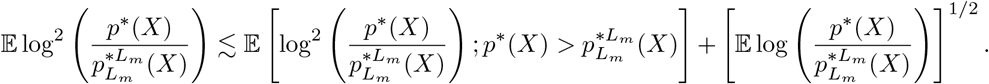

To achieve our result, we will show the first of these terms is 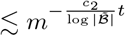 and one may use a similar proof to make the same deduction about the second term.

First we will split the term into two that represent the “distance” from *p*\* to *p^*L_m_^* and that from *p^*L_m_^* to 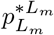:

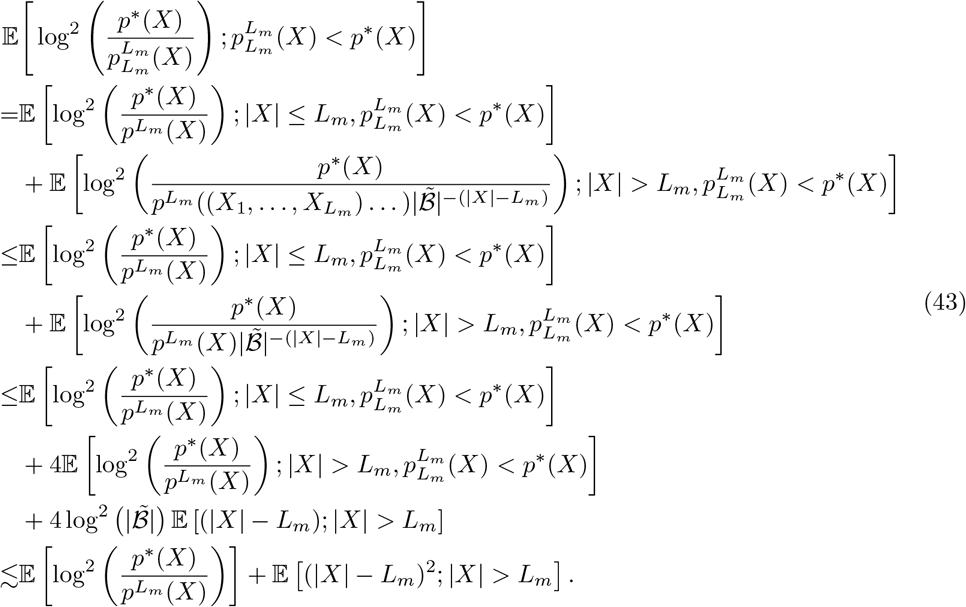

Now we will show each of these two terms 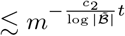 in turn.

We will first consider 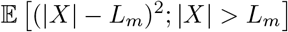.

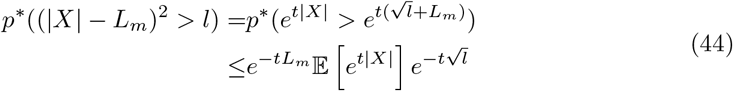

by Markov’s inequality, so

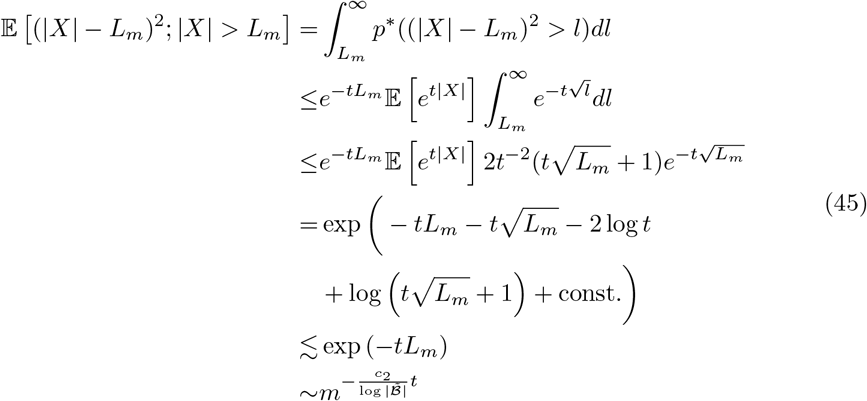

as desired.

For the other term in equation 43, again by lemma 31,

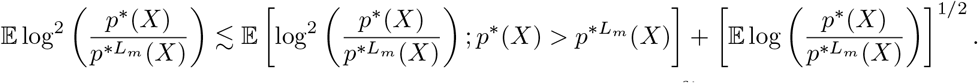

In this case, we will show that the first of these terms is 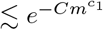 for some positive constant *C*, and by a similar proof one may show the same for the second. This will complete the proof of part 2.

If *p*\*(*X*) > *p^L_m_^*(*X*) ≥ 0, by the definition of *p^*L_m_^*,

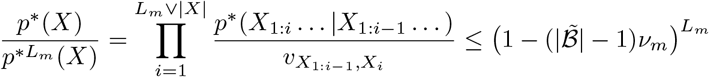

with the inequality by property (5) in proposition 24. Thus,

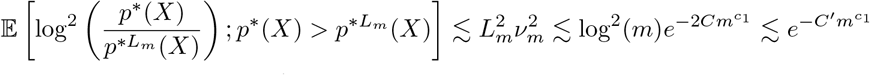

for two positive constants *C, C*′.

#### H.3.2 Consistency and rate

The proof of theorem 35 relies on a consequence of theorem 2.1 of Ghosal et al. [21], which is stated in a simplified form herein as theorem 33. Intuitively, the key challenge in establishing nonparametric consistency is that the size of the space of probability measures 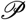 (infinite dimensional) may overwhelm the evidence provided by the data, leading to a posterior that is too spread out. To establish consistency, theorem 2.1 of Ghosal et al. [21] requires that the prior over probability measures is sufficiently large on a neighborhood of *p*\* (denoted 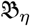), and sufficiently small on the complement of an effectively parametric (finite dimensional) subset of 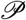 (denoted 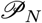).

##### Theorem 33.

*Say* 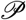 *is a set of probability measures*, 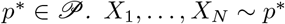. *X*_1_,…, *X_N_* ~ *p*\* *iid, d is the Hellinger distance*, Π *is a distribution on* 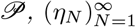 *is a sequence of positive numbers such that η_N_* → 0 *and* 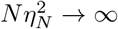, *and* 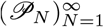 *are a sequence of subsets of* 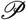. *Define, for positive η*,

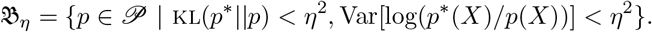

*Then if*

i. 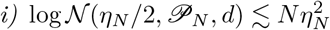
ii. 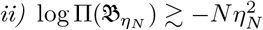
iii. *For an ϵ* > 0, 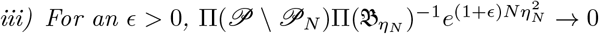

*Then for large enough M*,

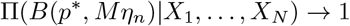

*in probability, where B*(*p*, δ*) *is a Hellinger ball of radius δ centered at p*\*

*Proof*. For some *C*,

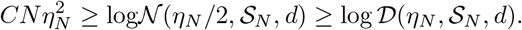

Defining 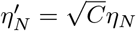, condition 2.2 in theorem 2.1 of Ghosal et al. [21] is satisfied for the sequence 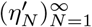. Note condition 2.4 is also satisfied by the above condition ii.

Note by lemma 8.1 in Ghosal et al. [21]

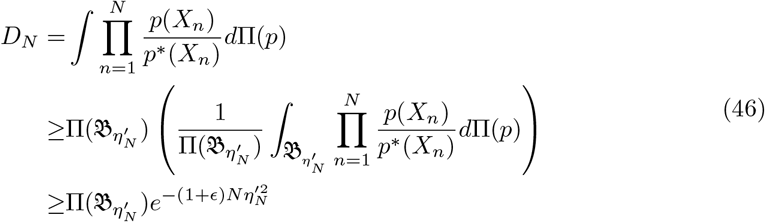

with probability 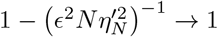. Call the set where this occurs *A*. As in the proof of theorem 2.1 of Ghosal et al. [21], for large enough *M, C*′, we may then use condition i to write

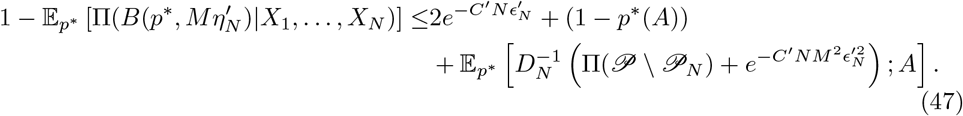

By conditions ii and iii, this last term → 0 for large enough *M*. Finally, write 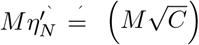 to get the result in terms of *η_N_*.

To work with sieves without restrictions on transition probabilities to 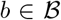 we need the following technical lemma.

##### Lemma 34.

*Assume for positive numbers* 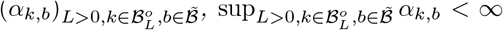 *and* 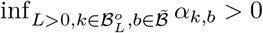. *Consider independent* 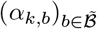 *priors on each simplex of* 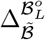 *indexed by* 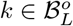. *Call the joint distribution* Π. *Then, for some C, ϵ* > 0, *for all ν* > *ν*′ *small, L*,

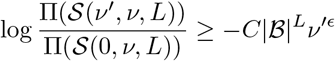

*Proof*. Define 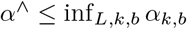. Let *Z_k_* ~ Dirichlet(*α_k,b_*)_*b*_ for some *k*. As a property of the Dirichlet distribution,

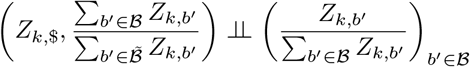

Call this later variable *Y_k_*, and note 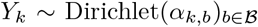. Now for any 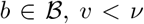, *υ* < *ν*, since 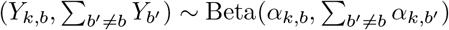,

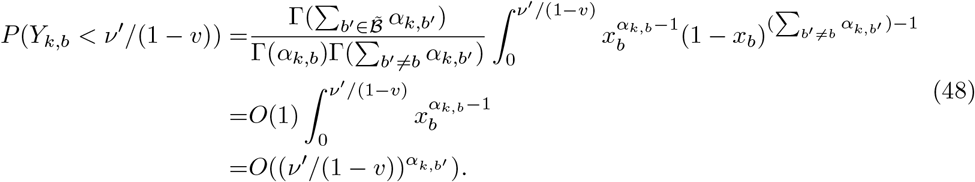

Thus, using a union bound, for some *C*, regardless of the choice of *k*,

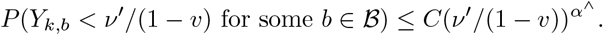

Thus, for some *C*′ > 0, calling *F*_*k*,$_ the density of *Z_k,b_*, noting *P*(*Z*_*k*,$_ > *ν*) = *O*(1) for small *ν*,

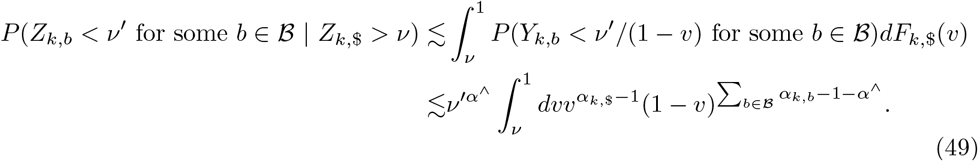

The integral is equal to the probability of a 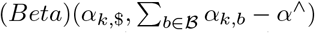 distribution being greater than *ν* and is thus *O*(1). For small enough *ν, ν*′, for some *C*′ > 0,

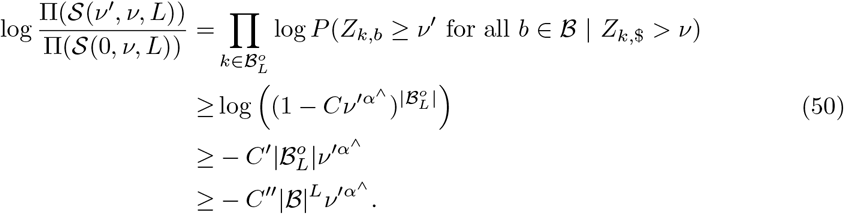

We can now prove the main result, establishing posterior consistency and the posterior convergence rate. We show that the prior in condition 29 satisfies the conditions of 33. In particular, we use sieves 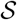 to define the effectively parametric subset 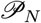 of the infinite dimensional space of probability measures 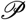, and then condition 29 controls the prior probability over the 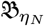 and 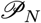.

##### Theorem 35.

*Assume p*\* *is sub-exponential and thus we can choose a prior as in condition 29. For any large enough M*,

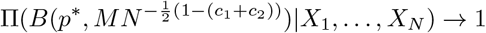

*in probability where B*(*p*\*, *δ*) *is a Hellinger ball of radius δ centered at p*\*.

*Proof*. The proof will proceed by checking the conditions of theorem 33. First define a monotonic sequence 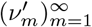 with 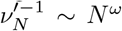, *ξ_N_* = *ξ*(*ν_N_, ν_N_, L_N_*), 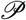 the set of distributions on *S*, and

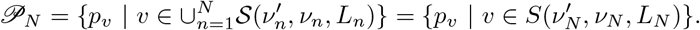

Throughout we will use 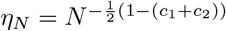 and so checking the conditions of theorem 33 will demonstrate a posterior concentration rate of 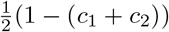.

First we will check condition i. Define, for 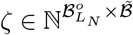, *ρ_N_* > 0,

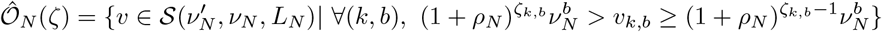

(where 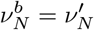 if *b* ≠ $ and equal to *ν_N_* otherwise) so that 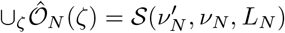 (Fig. S3).

Note that for *υ*_1_, 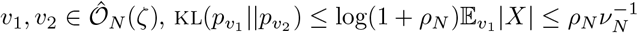 the last inequality as *p*(|*X*| > *L*||*X*| ≥ *L*) ≥ *ν_N_* where the last inequality comes from *p*(|*X*| = *L*||*X*| ≥ *L*) ≥ *ν_N_* and a geometric sum (this is where a distinction between *ν_N_* and 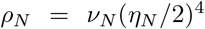 is necessary). Defining *d* as the Hellinger metric,

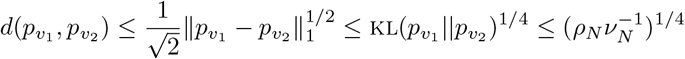

so picking *ρ_N_* = *ν_N_*(*η_N_*/2)^4^, for *υ*_1_, 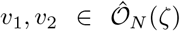, *d*(*p_υ_1__, p_υ_2__*) ≤ *η_N_*/2. Call 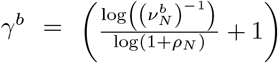 and note 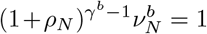. Thus the number of choices of 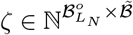 that give non-empty 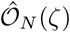, is bounded above by 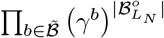. Note also that since *ρ_N_* → 0, 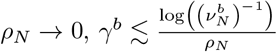. Now we can establish condition i of theorem 33:

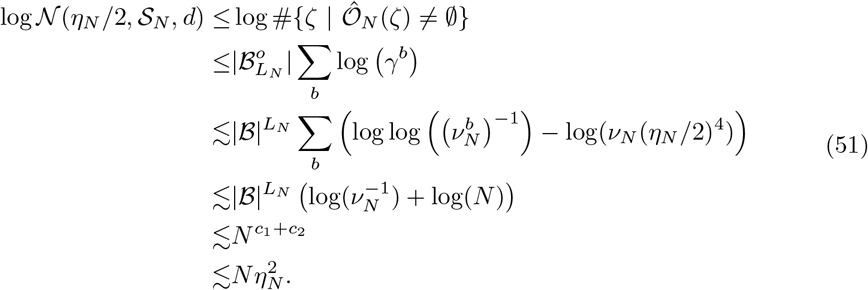

Now we will demonstrate condition ii. Define, as in theorem 33,

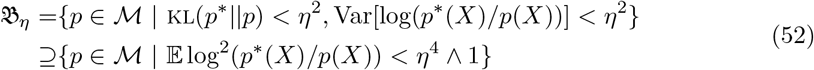

since 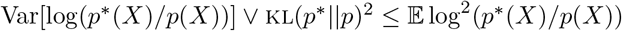.

Fix *N*. First we will delineate a volume in 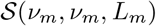 for any *m* > 0 that is within 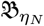. Using the definition of *ξ*, we can label a 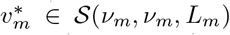 such that 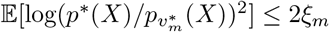. Note that if there exists a 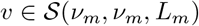 such that for some *ρ_m_* > 0 and all *k, b*, 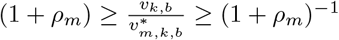 then

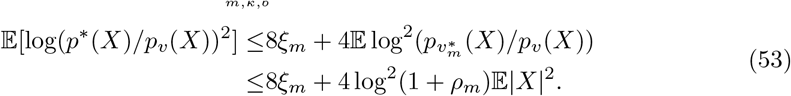

Now pick, for large enough *m*,

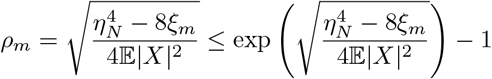

so that if 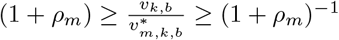 for all *k, b*, then 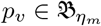.

Fixing *k*, the probability under a Dirichlet(*α_k,b_*)_*b*_ distribution of 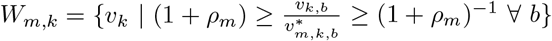 (depicted in Fig. S6(A)) is, considering the case where 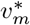 is on one of the corners of the simplex {*υ_k_* | *υ_k,b_* ≥ *ν_m_*}, at least

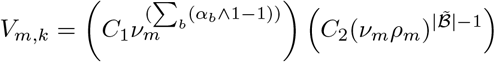

where the first term is a lower bound on the density and the second on the volume of *W_m,k_* and *C*_1_, *C*_2_ are constants depending on 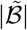. *C*_1_ > 0 as inf_*k,b*_ *α_k,b_* > 0. As well, one may check that the volume is minimized should 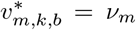 for all but one *b*; in this case, the volume forms a particular diamond-like shape with side-lengths scaled as *ν_m_ρ_m_* and dimensionality 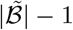 (Fig. S6(B))), (if 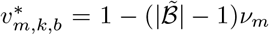, then the condition 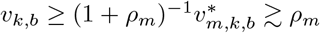 does not affect the *W_m,k_* for large *m* as *ν_m_* → 0) (Fig. S6).

Now we will lower bound the probability of 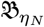 by the probability of the above defined volume for a particular *m, m_N_*. Call 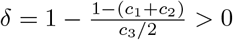 and define

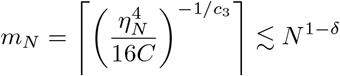

so that 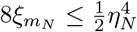 for all *m* ≥ *m_N_*, and *m_N_* → ∞. Now,

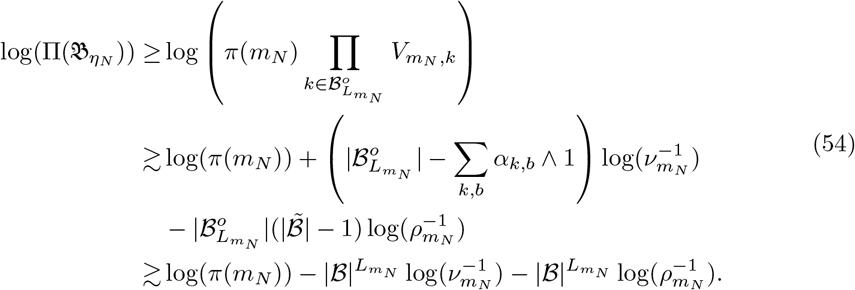

For the first term, due to condition 29, (*c*_1_ + *c*_2_) > (1 – *δ*)*ω* > (1 – *δ*)(*c*_1_ + *c*_2_), so,

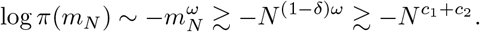

The second term has

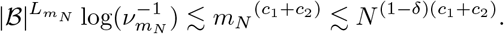

Finally, for the third, note that since 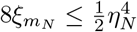,

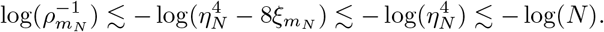

Thus,

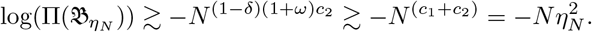

Finally, for condition iii, note

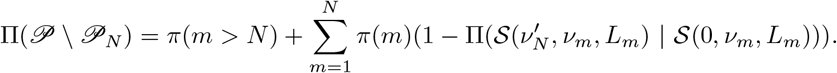

From lemma 34, we have, for *C, C*′, *ϵ* > 0, the second term is dominated by

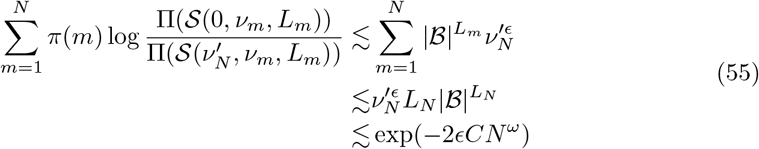

for some *C* > 0. On the other hand, since one may check that *π*(*m* + 1)/*π*(*m*) < 1/2 for all *L*, we have *π*(*m* > *N*) ≤ *π*(*N*). Thus,

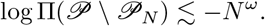

Now we may write, for any *ϵ* > 0, since *ω* > *c*_1_ + *c*_2_

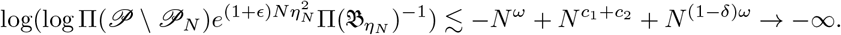

**Figure S6:**
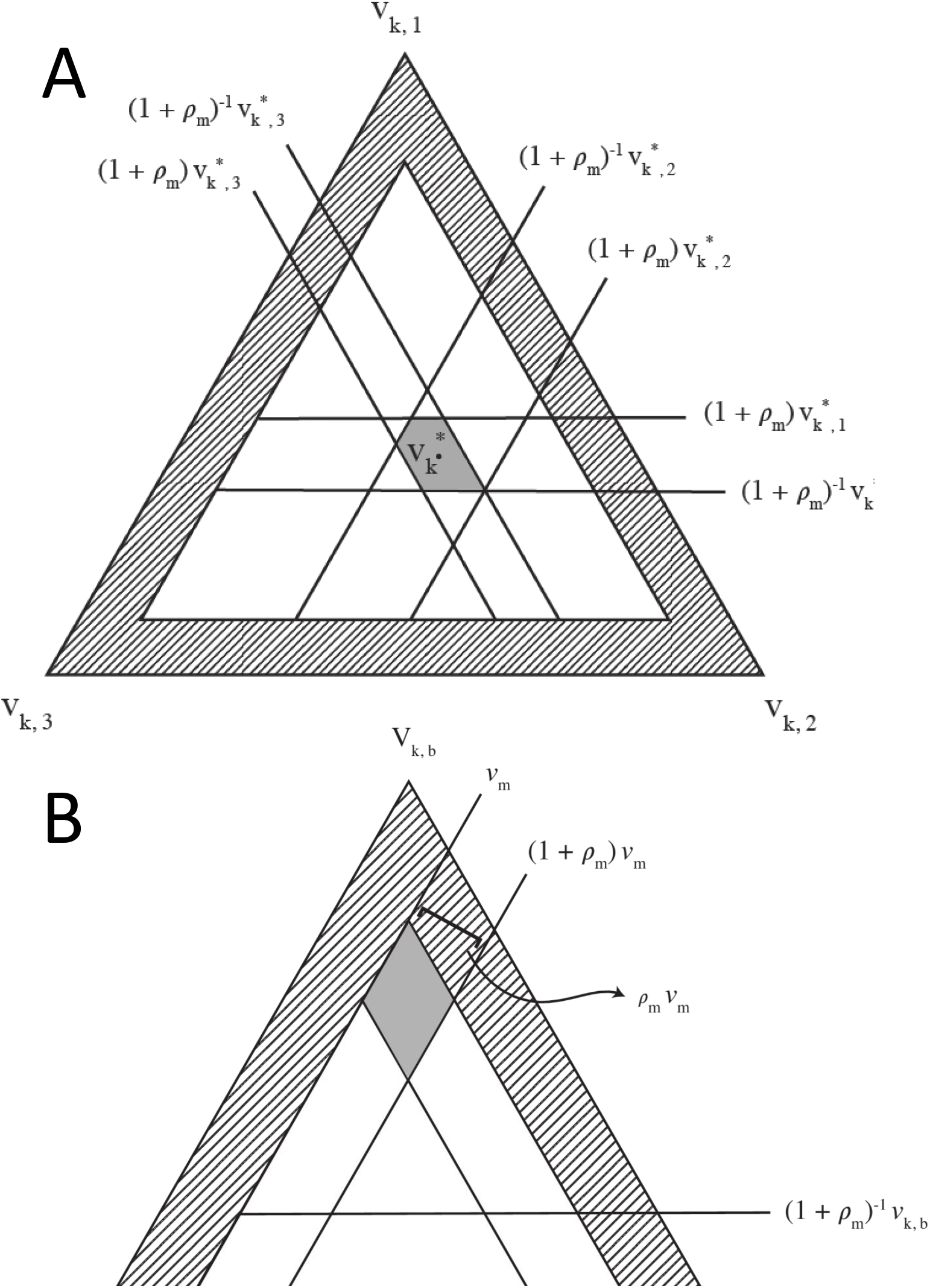
(A) Example of a set *W_m,k_* (solid gray) where 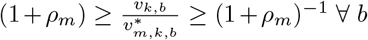 on 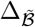 for a particular *k* and *m* when 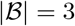. (B) Depiction of minimum volume possible. The dashed region represents those transition probabilities that have components less than *ν_m_*.

#### H.3.3 Use in practice

Theorem 35 reveals that the choice of prior controls a kind of bias-variance tradeoff in the model’s posterior. In particular, from condition 29 we have

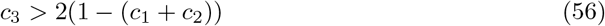

Decreasing the prior hyperparameters *c*_1_ and *c*_2_ decreases the width of the posterior distribution (which plays the role of variance). However, reducing *c*_1_ and *c*_2_ forces down *c*_3_ (by the definition of *ξ*), and this reduces the weight that the prior places on larger sieves that can match the data distribution better (i.e. sieves with lower *ξ*(*ν, ν, L*) values), consequently increasing the model’s bias. When *c*_1_ and *c*_2_ become low enough, the bias becomes overwhelming, equation 56 is violated, and consistency is no longer guaranteed.

In practice it is often sensible heuristically to set *ν_m_* = 0. In the case, for instance, of short-read sequencing data, there’s relatively little correlation between the letters of the read and where it terminates. The probability of stopping is thus often similar across different kmers, even when comparing among kmers of different length. As the posterior concentrates at a roughly constant stopping probability, even a low one, *ν_m_* quickly becomes irrelevant as it decays to zero exponentially. When *ν_m_* = 0, the prior simplifies: it can be written as a distribution over lags *π*(*L*) times independent Dirichlet priors on each 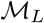 for *L* ∈ {1, 2,…}. The prior over lags takes the form

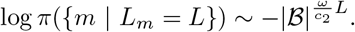

Since *ω* > *c*_2_, we may write 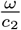 as 1 + *c* for a small *c* > 0.

## I Toy models

In this section we describe in depth our simulation experiments.

### I.1 Finite lag models

This subsection describes experiments conducted to study in practice the finite lag consistency results described in Sections E and F, and includes details on the results presented in Section 2 and Figure 2.

#### I.1.1 Setup

To simulate data, we used an AR model with parameters *θ* = (*A, B*) defined by the function,

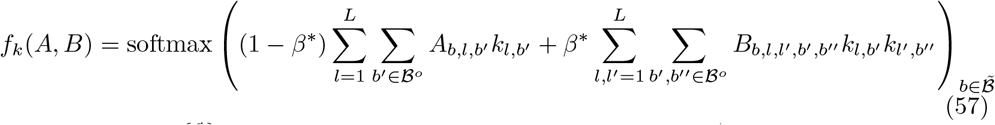

where 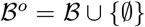 and *k_l,b_* is 1 if *k_l_* = *b* and 0 otherwise. The AR model thus takes the form of a multi-output logistic regression, with *β*\* controlling the contribution of the pairwise interaction terms. In each independent simulation, rows of the matrix *A* were sampled following,

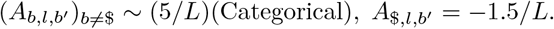

for each *l, b*′, where (Categorical) denotes a one-hot encoded sample from a Categorical distribution with uniform probabilities. The matrix *B* was generated similarly,

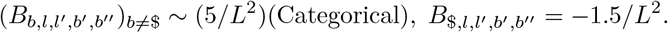

for each *l, l′, b′, b″*. Simulations were repeated five times for each *β*\* value. We set *L* = 5.

We trained the *θ* parameter in the AR models using maximum likelihood and the *h, θ* hyperparameters in the BEAR model using empirical Bayes. In both cases, we trained without mini-batching, using 1000 steps of the Adam optimizer with a training rate of 0.05 [33].

To approximate the KL divergence and total variation distance between the models and the data, 2,000 independent sequences were sampled from the data-generating distribution *p*\* and used to calculate averages of log (*p*\*(*X*)/*p*(*X*)) and 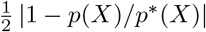 respectively, where *p* is either the maximum likelihood estimator (for the AR models) or the posterior predictive (for the BEAR models, estimated using the maximum *a posteriori* value). (Note that the total variation distance is equal to half the *L*^1^ distance since the set of sequences is countable.)

The parameter *A* is not identifiable, so to compare between the value of *A* inferred by the models and the true data-generating value, we transformed *A* to a canonical representation. Define 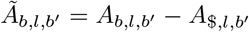 and define the canonical representation

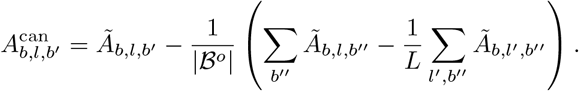

##### Proposition 36.

*Two linear AR matrices A, A′ define the same linear AR model of lag L if and only if A^can^ = A^′can^*.

*Proof*. Define the vector space

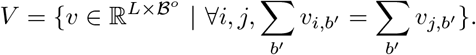

One hot encodings of sequences of length *L* are contained in *V*. As well, it can be seen that *V* is spanned by the vectors 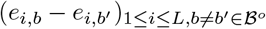 (where *e_i,b_* is the indicator of position *i, b*) and the vector consisting of ones in each entry. This basis of *V* is made up of linear combinations of one hot encodings of sequences of length *L* and thus the span of one hot encodings of sequences of length *L* is *V*. The orthogonal complement of *V* is spanned by (*e_i_* – *e*_1_)_1<*i*≤*L*_ where *e_i_* is 1 at position *j, b* if *j* = *i* and 0 otherwise. The transformation

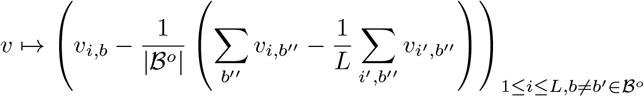

preserves *V* and annihilates the orthogonal complement of *V* and is thus the orthogonal projection onto *V, P_V_*.

Thanks to the softmax in Equation 57, two linear AR matrices *A* and *A*′ define the same linear AR model if there is a constant *C* such that for all sequences *k* of length *L* and 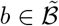,

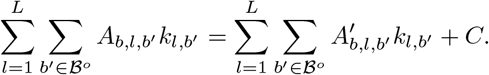

This is equivalent to the condition

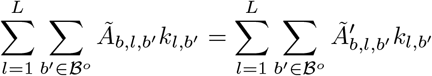

for all *k, b* and thus to the condition

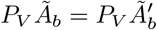

for all *b*.

#### I.1.2 Results

We first fixed *L* at the same value as the simulation data, to study the effect of the structured prior in the BEAR model. Figure 2A shows the convergence in KL of each model as the dataset size increases, and Figure S8 the convergence in total variation distance. Figure 2B shows the convergence of the hyperparameter *h* in the BEAR model. In Figure S7, we compare the parameter *A* inferred with the AR model to the true data-generating value using the Frobenius norm of the canonical representation of each; likewise for the parameter *A* inferred with the BEAR model. In this well-specified case, we see that the BEAR model parameter estimate converges just as quickly as the AR model.

**Figure S7:**
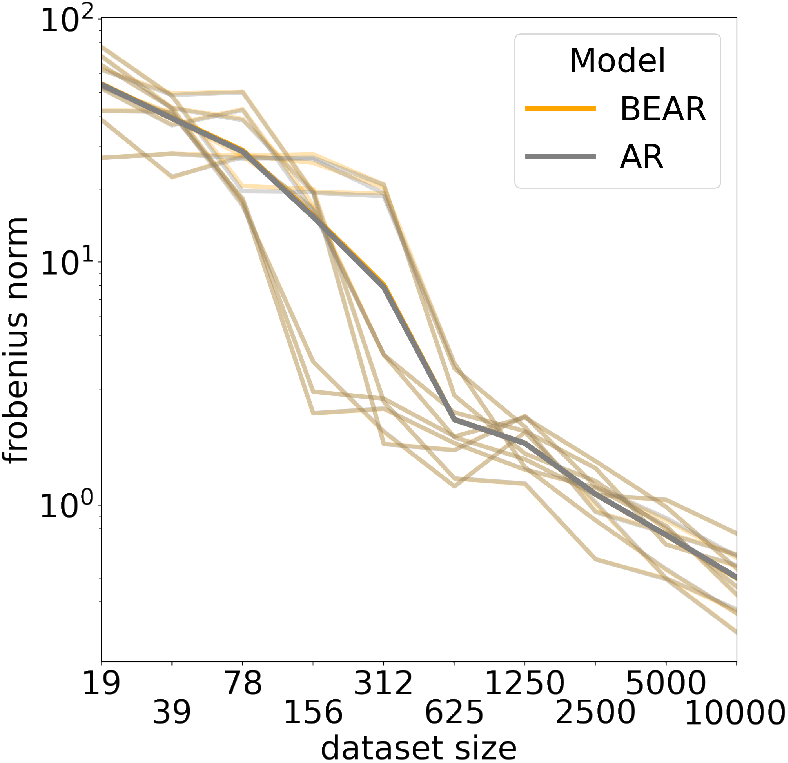
Frobenius norm between the canonical representation (Section I.1) of the AR model parameters *θ* inferred by fitting an AR model with maximum likelihood and those inferred by fitting the BEAR model with empirical Bayes, in the well-specified (*β*\* = 0) case. Thick lines show the average across five independent simulations (small lines). Note that the differences between the two models are indistinguishable relative to the variation across datasets and the variation as dataset size increases.

**Figure S8:**
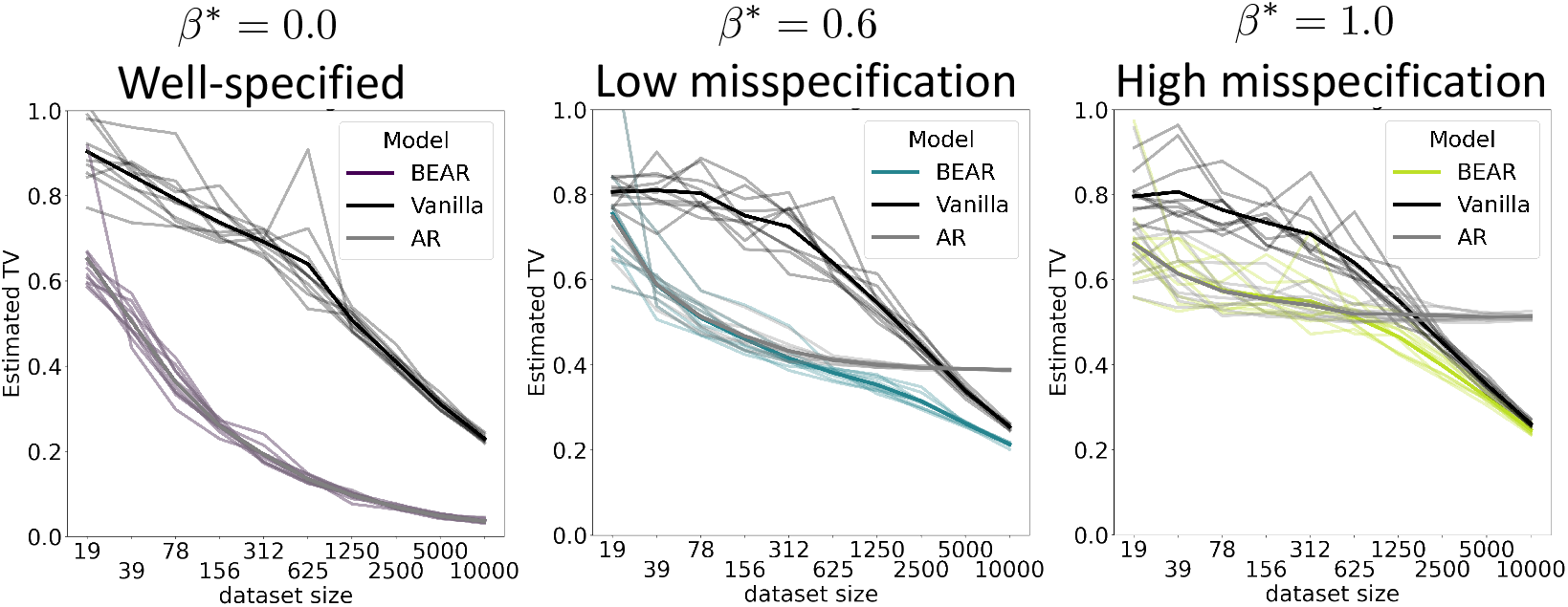
As in Figure 2A, except using the total variation distance in place of the KL norm.

**Figure S9:**
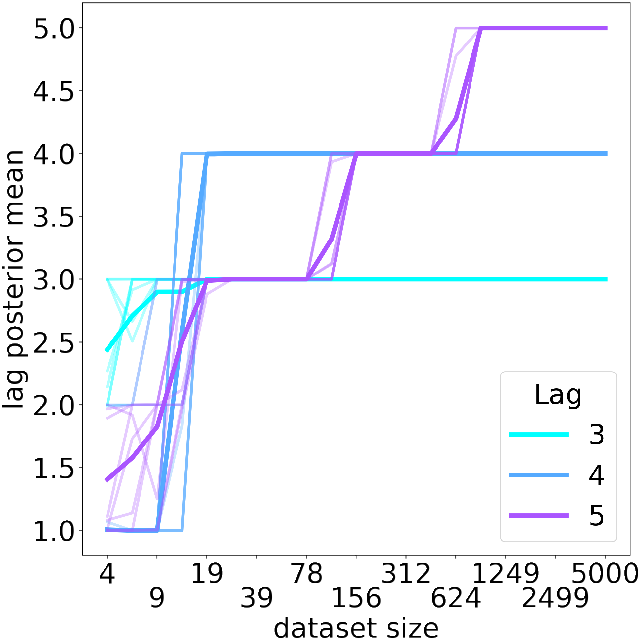
Mean of the BEAR model posterior over lags, as a function of dataset size. Thick lines show the average across five independent simulations (thin lines).

Next we considered inference of *L*. We simulated data from models with different *L* values (*L* ∈ {3, 4, 5}) and *β*\* = 0. We computed the expected value of *L* under the posterior with a uniform prior on lags from 1 to 8. Figure S9 shows that the inferred lag converges to the true data-generating value.

### I.2 Infinite lag models

This subsection describes experiments conducted to study the infinite lag (nonparametric) consistency results of Section H in practice.

#### I.2.1 Setup

To generate from a distribution that was not a finite lag AR model, we chose the first letter in each sequence *X* uniformly from the alphabet 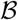, then sampled the rest of the sequence following,

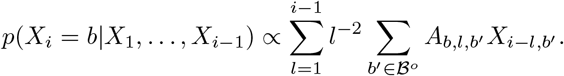

In each independent simulation, the parameter *A* was sampled as *A_b,l,b′_* ~ Bernoulli(0.2) for each *l, b* and *b*′ ≠ $, and as *A_b,l,b′_* ~ (0.2)(Bernoulli(0.2)) for each *l, b* and *b*′ = $.

Following Section H.3.3, we set *ν_m_* = 0 and used the prior on lags *π*(*L*) ∝ exp(−4^(1+*c*)*L*^). We used a Jeffreys prior (*α_k,b_* = 1/2 for all *k, b*) and took the maximum *a posteriori* value of *L* and *υ*. We also considered the maximum likelihood estimator of *L* (i.e. with the prior dropped). To approximate the KL divergence and the total variation distance, we used 30,000 samples; the training procedure was otherwise the same as in Section I.1.

#### I.2.2 Results

We examined the convergence of the posterior predictive distribution of the BEAR model for different values of the prior hyperparameter *c*. In all cases we see convergence to *p*\* in both total variation and KL (Figure S10AB). Decreasing *c* produces a longer-tailed prior, making the maximum *a posteriori* value of *L* diverge more quickly with dataset size (Figure S10C). In this example, decreasing *c* yields faster convergence to *p*\*. Using the maximum likelihood value of *L* (equivalent to an improper uniform prior) yields even faster convergence to *p*\*. As discussed in Section H.3.3, lower *c* corresponds to larger *c*_2_, and so is expected to yield lower posterior variance but larger bias; in this simulation, the reduction in bias clearly contributes more to accurate density estimation. This may be because the data-generating distribution is close enough to a finite-lag Markov model that the asymptotics of the BEAR model behave similarly to the finite-lag case.

**Figure S10:**
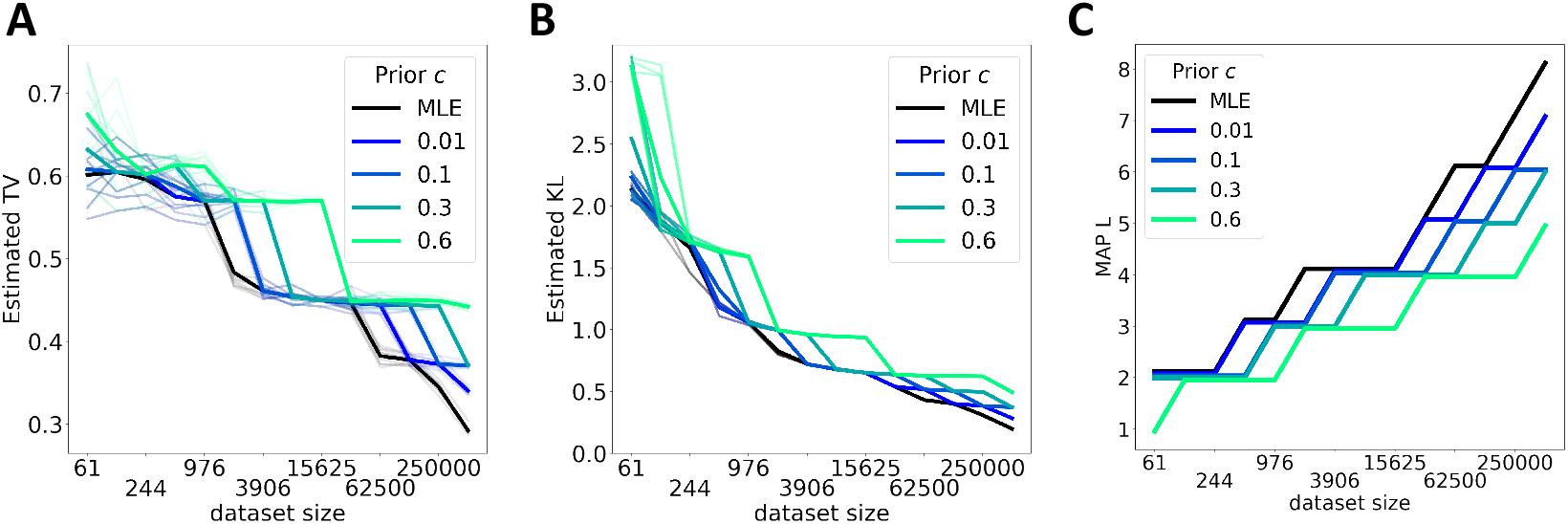
Convergence in total variation (A) and KL (B) between data-generating distribution and model. Thick lines indicate averages across five individual simulations (thin lines). (C) Maximum *a posteriori* estimator of the lag *L* in an individual example simulation.

### I.3 Hypothesis testing

This subsection describes experiments conducted to study the hypothesis testing consistency results of Section G in practice.

#### 1.3.1 Setup

We used the same setup as in Section I.1.1, including the same training and divergence estimation procedures, and sampled datasets from a linear AR model with different values of *β*\*.

In the goodness-of-fit test, we set 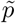 (the model we aimed to test) to a linear AR model with the true, data-generating value of the parameter *A* but *β*\* = 0. We embedded the same linear model, with the same value of *A* and *β*\*, in the BEAR model to compute a Bayes factor. Here we set *h* = 10^-3^, and fixed *L* at the data-generating value, *L* = 5.

In the two-sample test, instead of comparing to 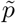 directly, we compared to samples drawn from 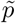. Here we used a Jeffreys prior rather than embed a more complex AR model. We explored both fixing *L* = 5 and using a truncated uniform prior *π* (*L*) = 1/8 for *L* from 1 to 8 (to evaluate both forms of the consistency results in Section G).

#### 1.3.2 Results

We first examined the consistency of the goodness-of-fit test, using the Bayes factor 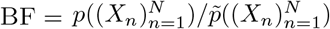 which compares the probability of the data under the BEAR model to the probability under the model of interest 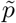. Figure S11A shows the Bayes factor diverge to +∞ when the data does not match the model (*β*\* > 0), but diverge to –∞ when the data does match the model (*β*\* = 0). We also explored the Bayes factor as function of *h*, holding the amount of data fixed at *N* = 2500 (Figure S11B). In the limit *h* → 0, the BEAR model reduces to its embedded AR model 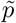, and so the Bayes factor converges to 0. On the other hand, in the limit *h* → ∞, the BEAR model becomes diffuse and the Bayes factor diverges to negative infinity (accepting the null hypothesis). Intermediate values of *h* in effect “center” the test at the model 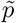 we aim to evaluate, increasing its power to detect differences between the data and the model [6].

We next examined the consistency of the two-sample test, using the Bayes factor 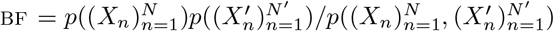, which compares the probability of the two samples being drawn from separate distributions to the probability of their being drawn from the same distribution. Both when using the Bayes factor computed with fixed lag *L* = 5, and when using the Bayes factor computed by marginalizing over a truncated uniform prior on *L*, we find consistency, with the Bayes factor diverging to +∞ when *β*\* > 0 and to –∞ when *β*\* =0 (Figure S12).

**Figure S11:**
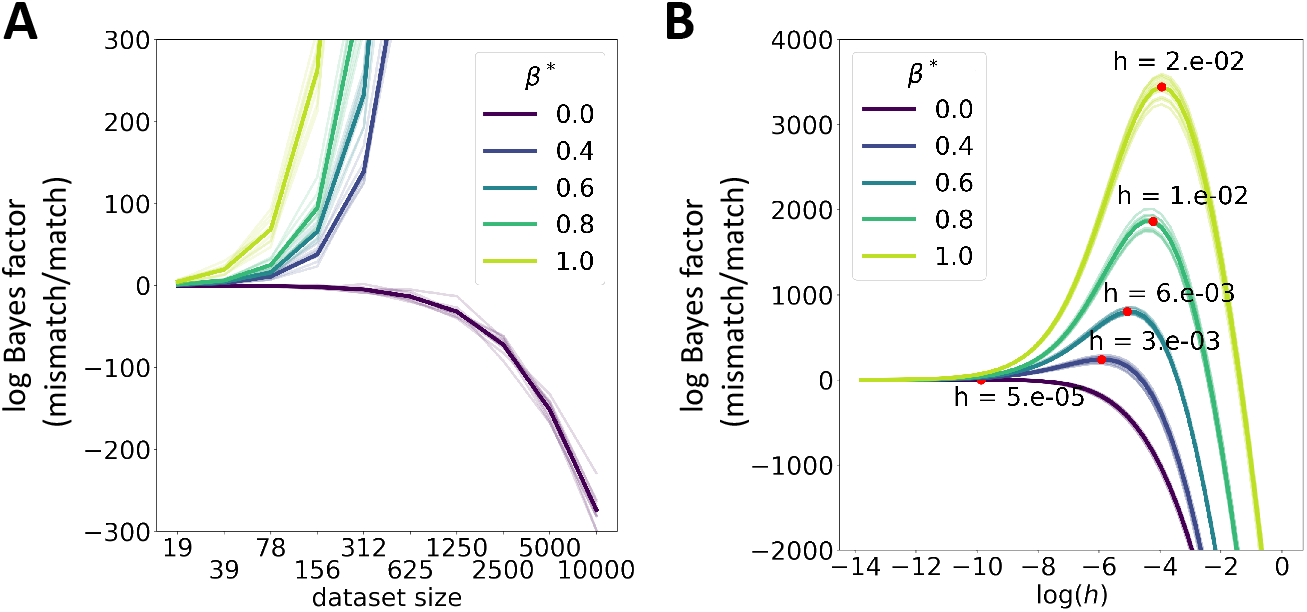
(A) Log Bayes factor for the BEAR goodness-of-fit test. (B) Log Bayes factor as a function of the hyperparameter *h*, with peaks identified by red points. In both subfigures, thick lines are averages across five simulations (thin lines).

**Figure S12:**
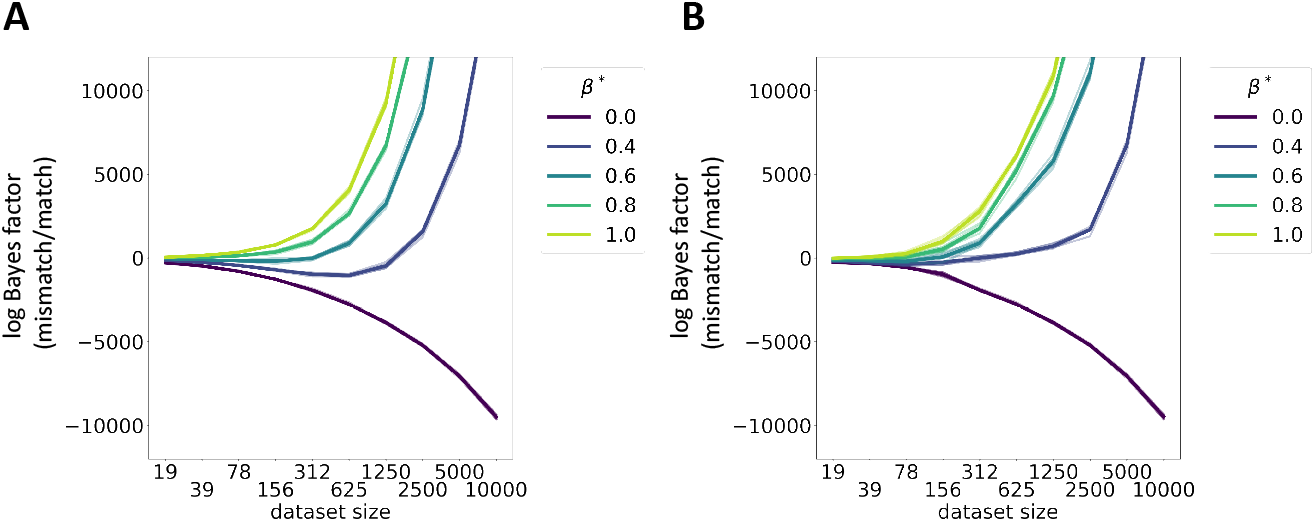
(A) Log Bayes factor for the BEAR two-sample test, using fixed *L*. (B) Log Bayes factor for the BEAR two-sample test, marginalizing over a truncated prior on *L*. In both subfigures, thick lines are averages across five simulations (thin lines). Dataset size is the size of each individual dataset that the two-sample test compares, not their pooled size.

## J Scalable inference

In this section we describe how BEAR models were trained at large scale on real data.

### J.1 Stochastic gradient estimates

Let 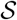 be a set of length *L* kmers *k* in 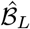 chosen uniformly at random (a minibatch). Then, we can form an unbiased stochastic gradient estimate of the marginal likelihood as

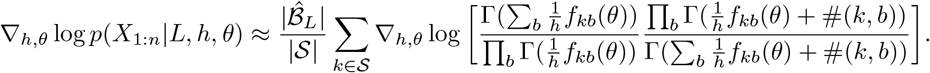

Note also that it is straightforward to parallelize the training algorithm by sending individual minibatches to individual processors at each step, then compiling the results.

**Table S1:** Dataset sizes. In nucleotides (nt). Dataset abbreviations as in Table 1.

| Dataset | Total nt | Max. sequence length (nt) |
| --- | --- | --- |
| YSD1 | 151,691,700 | 150 |
| *A. th.* 1 | 3,238,613,507 | 100 |
| *A. th.* 2 | 2,485,960,312 | 100 |
| *A. th.* 3 | 6,831,756,793 | 100 |
| PBMC | 34,935,800,234 | 91 |
| HL | 24,185,778,348 | 91 |
| GBM | 21,506,001,361 | 65 |
| HC | 2,283,930,547 | 202 |
| CD | 1,052,405,190 | 202 |
| UC | 956,179,237 | 202 |
| Bact. | 1,388,421,381 | 6,358,077 |

### J.2 Extracting summary statistics

KMC counts kmers in large sequence datasets, outputting a list of kmers *k* and counts #*k* that is typically too large to fit in memory. However, our inference procedure requires full count vectors #(*k*, ·). We take advantage of the lexicographical ordering of KMC’s output to merge kmer counts into count vectors in a (single pass) streaming algorithm. We also take advantage of the lexicographical ordering to construct count vectors #(*k*, ·) for all lags *L* given just KMC’s output for the largest lag *L*, thus reducing the number of times KMC needs to be run; this too is done using a single pass streaming algorithm. In order to quickly evaluate models by heldout marginal likelihood, it is convenient to store together the counts #(*k*, ·) associated with both the training and testing datasets. We accomplish this by merging the KMC output for different datasets as part of the same single pass streaming algorithm. This dataset merging is also useful in training the reference-based models proposed in Section L.1, and we merge reference genome counts with sequencing dataset counts in the same way.

### J.3 Code availability

Code for implementing BEAR models is available at https://github.com/debbiemarkslab/BEAR, along with documentation (including a tutorial for getting started and reproducing basic results); it is available under an MIT license. The models are implemented using TensorFlow and TensorFlow Probability, available under an Apache License [2, 15]. The code also uses NumPy [26], SciPy [66], and BioPython [10] (all BSD 3-Clause licenses). KMC is available under a GNU GPL 3 license.

## K Datasets

Here we briefly describe each data type and dataset used in evaluating BEAR models, along with some motivation for each. NCBI accession numbers and links for each dataset can be found in the supplementary table Datasets.xlsx. Dataset sizes are listed in Table S1. All data is publicly available for research use. Patient data was anonymized by the creators of each dataset, and further details on ethical oversight and patient consent can be found in the cited links and papers.

### K. 1 Whole genome sequencing

Whole genome sequencing is a standard technique for measuring genome sequences. It is often the starting point for running a genome assembly algorithm or variant caller, which aims to infer (non-probabilistically) the underlying genome from the read data. Directly modeling sequencing reads can be interesting, however, since (a) there are typically portions of the genome that are difficult to reliably assemble, such as centromeres and telomeres, (b) there may not be enough data to reliably detect variants via standard variant callers or assembly, and (c) although the experiment may be directed towards a particular organism’s genome other DNA may still be present.

- **YSD1** This is a bacteriophage found in the waterways of the United Kingdom which infects *Salmonella*. It was chosen as an example of a relatively small genome sequencing experiment (phage genomes are short). The sequencing experiment was reported in Dunstan et al. [17].
- **A. th.** Arabidopsis thaliana is a small flowering plant, used as a model organism in plant research. Structural variants are extremely complicated in plants, making traditional variant-calling methods challenging, and kmer-based analysis approaches are of considerable ongoing interest in the literature (see e.g. Voichek and Weigel [67]). The datasets are from the 1001 Genomes Consortium, https://1001genomes.org/ [1].

### K.2 Single cell RNA sequencing

Single cell RNA sequencing is an increasingly ubiquitous technique for characterizing the transcriptional state of cells. It is used to discover new cell types, track development and disease, as a readout in cellular engineering efforts, and more. Most analysis techniques coarse-grain the data by just counting transcripts or isoforms. Statistical modeling of reads at the nucleotide level may lead to new insight into the joint distribution of sequences and their expression levels, accounting for such phenomena as somatic variation and RNA editing. Single cell RNA sequencing is increasingly used as a method for understanding tumors and their microenvironment; cancer involves both genome mutations as well as transcriptional changes.

- **PBMC** Samples of peripheral blood mononuclear cells are easy to collect from humans, making this a standard type of single cell RNA sequencing dataset. These cells were taken from a healthy donor. The dataset is from 10x Genomics, using its v3 technology.
- **HL** These cells come from a human dissociated lymph node tumor, from a 19-year-old male Hodgkin’s lymphoma patient. The dataset is from 10x Genomics, using its v3 technology.
- **GBM** These cells were taken from a patient with glioblastoma, the most common primary brain cancer in adults, and include both tumor and peripheral cells. The dataset was reported in [13] and uses a distinct technology from 10x Genomics methods.

### K.3 Metagenomics

Metagenomics is an increasingly ubiquitous technique for characterizing microbiomes, including human and environmental microbiomes. Analysis often proceeds by local assembly, annotation of genes or taxa, etc. Statistical modeling of reads at the nucleotide level avoids this coarse graining and can enable detection and analysis of changes in the microbiome outside known genomic elements.

All three of the metagenomics datasets analyzed in the prediction experiments are from [41], a study of inflammatory bowel disease (IBD) as part of the Integrative Human Microbiome Project, and were taken from stool samples. IBD affects more than 3.5 million people worldwide.

- **HC** This dataset was collected from a control patient without IBD.
- **CD** This dataset was collected from a patient with Crohn’s disease, a form of IBD involving relapsing and remitting inflammation of the gastrointestinal tract.
- **UC** This dataset was collected from a patient with ulcerative colitis, a form of IBD involving relapsing and remitting inflammation of the colon.

We also examined metagenomics datasets from a study of kidney transplants [55]. Viral transmission from donor to recipient has been associated with complications and increases the risk of allograft failure. Schreiber et al. [55] performed metagenomic sequencing on patient urine samples before and after transplant to assess viral transmission. Further description of this dataset can be found in Section O.

### K.4 Full assembled genomes

Comparisons between distant species are challenging due to complex and large scale genomic changes over evolutionary time. However, generative probabilistic models of protein sequences separated by billions of years of evolution has yielded direct insight into their functional constraints, as well as improved understanding of the large scale evolution of life on earth [28, 51]. As a first step towards extending these ideas to whole genomes, we analyzed diverse bacterial genomes from across the tree of life.

- **Bact.** We examined reference bacterial genomes available in RefSeq [46]. Genomes were selected to be taxonomically diverse, representing different genera and families from across the kingdom of Bacteria; the NCBI accessions are listed in Datasets.xlsx.

## L Prediction experiments details

Here we provide details on the results reported in the **Predicting sequences** and **Measuring misspecification** subsections of the results (Section 6).

### L.1 Model architectures

- **Linear** The linear model is the same as that used in the toy experiments,

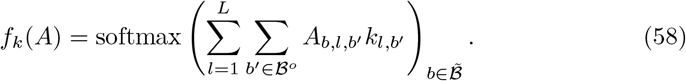
- **CNN** We use a four layer convolutional neural network with the architecture: input ↦ convolution ↦ elu ↦ elu ↦ softmax ↦ output, where the convolution is one-dimensional and the elu layers are exponential linear units. Layer normalization was used before each of the elu nonlinearities [4]. Exact details on the model architecture can be found in the supplementary code (Section J.3, function `make_ar_func_cnn` in `ar_funcs.py`).
- **Reference-based** Biologists often make use of a reference genome - a canonical example sequence that is intended to be representative of a species - in analyzing genome sequencing data; reference transcriptomes are used similarly in RNA sequencing analysis, etc.. Reads are aligned to the reference in order to infer the portion of the underlying genome or transcriptome that the read originated from. We built on this basic idea to design an AR model that uses a reference sequence to make predictions. In particular, let #_ref_(*k, b*) denote the number of times the length *L* +1 kmer (*k, b*) occurs in the reference sequence(s). One way to form a prediction is by normalizing these counts for each lag, i.e. *f_k,b_* = #_ref_(*k, b*)/∑_*b′*_, #_ref_(*k, b′*). We go a step further by (1) accounting for possible mutational or sequencing noise using a Jukes-Cantor mutation model, and (2) accounting for short reads by learning the stop symbol probability. Our complete model is

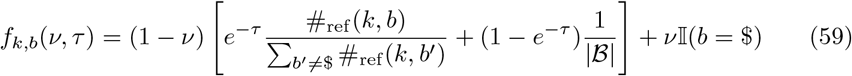

where *τ* ∈ [0, ∞) is the (scalar) Jukes-Cantor time parameter, *ν* ∈ [0,1], and 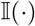 is the indicator function that takes value 1 when the expression is true and 0 otherwise. The reference sequences for each dataset are listed in the supplementary table Datasets.xlsx. In analyzing human single cell RNAseq data we pooled multiple reference transcriptomes. We included the reverse complement of each sequence as well as the original sequence when constructing the reference kmer transition counts.

**Table S2:** Training parameters. Train-test splits and Adam optimization parameters. Dataset abbreviations as in Table 1. Accum. steps stands for accumulation steps, the number of steps gradients were accumulated over. Paired end reads were treated as separate and split into train and test sets independently.

| Dataset | Train/test split | Epochs | Learning rate | Accum. steps |
| --- | --- | --- | --- | --- |
| YSD1 | 3:1 on reads | 500 | 0.01 | 10 |
| *A. th.* 1 | 3:1 on reads | 15 | 0.02 | 100 |
| *A. th.* 2 | 3:1 on reads | 15 | 0.02 | 100 |
| *A. th.* 3 | 3:1 on reads | 3 | 0.02 | 100 |
| PBMC | 3:1 on reads | 3 | 0.02 | 100 |
| HL | 3:1 on reads | 5 | 0.02 | 100 |
| GBM | 55:23 on cells | 4 | 0.02 | 100 |
| HC | 3:1 on reads | 10 | 0.02 | 100 |
| CD | 3:1 on reads | 10 | 0.02 | 100 |
| UC | 3:1 on reads | 10 | 0.02 | 100 |
| Bact. | 500:166 on genomes | 2000 | 0.01 | 1 |

### L.2 Training

The maximum marginal likelihood lag *L* was chosen for the vanilla BEAR model (with prior concentration parameter *α_k,b_* = 0.5 for all *k, b*). We found in general that the posterior was strongly peaked at a particular lag (Figure S15). All other models (both BEAR and AR) were run with this same lag (that is, we did not integrate over all lags in the BEAR model). Using a fixed lag *L* as a comparison point provides a controlled study of the effects of switching from an AR model of transition probabilities to the BEAR model’s AR-structured prior, and choosing *L* based on the vanilla BEAR model ensures that the comparison to the vanilla BEAR model is conservative.

The kmer count summary statistics were shuffled once before training (in chunks, due to the large size dataset size), and visited in the same order across epochs. Training was initialized only once; preliminary experiments suggested that training was robust to changes in the random seed. Gradient updates were computed in parallel across two GPUs, at double precision. The minibatch size was 250,000. Gradients were accumulated across minibatches to reduce variance (that is, the gradients from multiple minibatches were added together), and optimization was performed using Adam [33]. Models were trained to convergence. Detailed training hyperparameters are displayed in Table S2. The CNN models used 30 filters of width 8, except in the case of YSD1 where the filter width was reduced to 5 (for both BEAR and AR models); other neural network architecture hyperparameters are given in the supplementary code (function `make_ar_func_cnn` in `ar_funcs.py`). Experiments were run on an internal cluster (Tesla K80, Tesla M40 and Tesla V100 GPUs).

### L.3 Evaluation

Accuracy was evaluated based on the maximum likelihood prediction (in the case of AR models) and the maximum *a posteriori* prediction (in the case of BEAR models). Ties in prediction probabilities were resolved uniformly at random.

The perplexity was calculated based on the heldout test dataset as

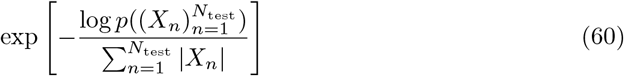

where 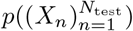 is the probability of the heldout data conditional on the maximum likelihood parameter value (in the case of AR models) or the marginal probability of the heldout data under the posterior predictive distribution (in the case of BEAR models).

**Table S3:**
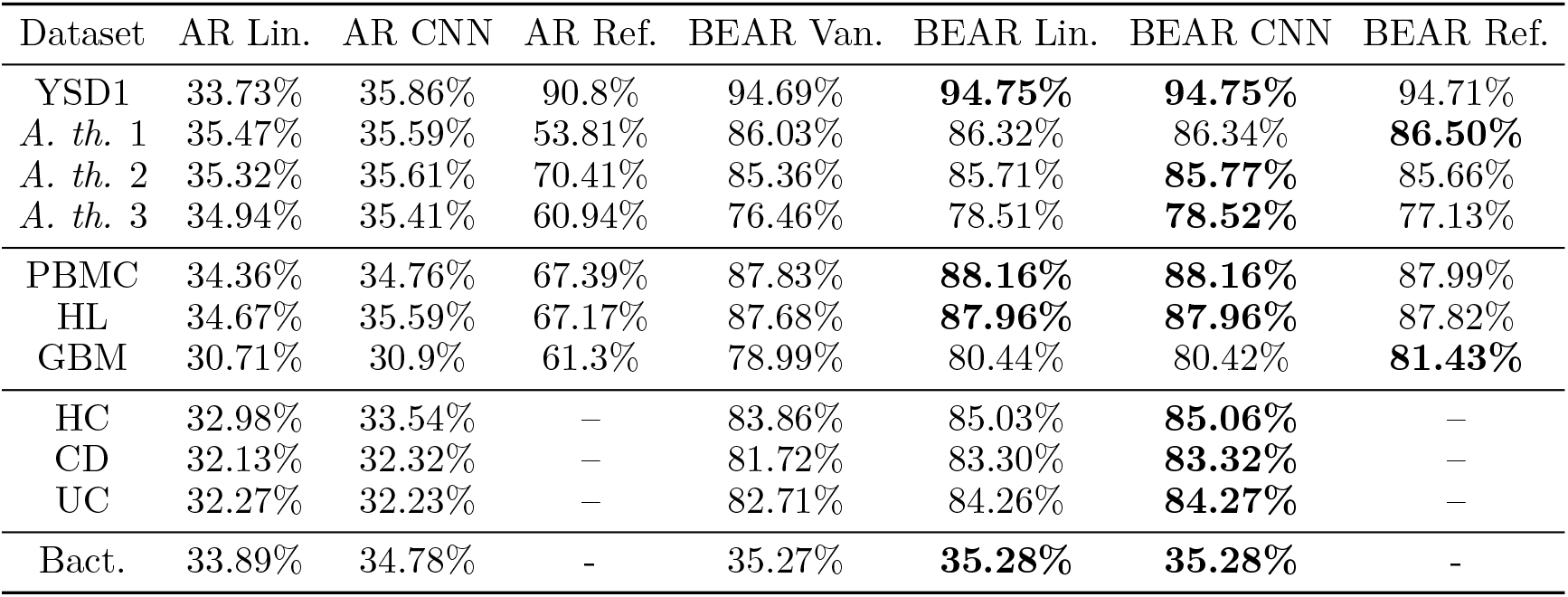
Predictive accuracy. *Whole genome sequencing data* YSD1: A Salmonella phage. *A. th.: Arabidopsis thaliana*, a plant (datasets represent different individuals). *Single cell RNA sequencing data* PBMC: peripheral blood mononuclear cells, taken from a healthy donor. HL: Hodgkin’s lymphoma tumor cells. GBM: glioblastoma tumor cells. *Metagenomic sequencing data* HC: healthy (non-CD and non-UC) controls. CD: Crohn’s disease. UC: ulcerative colitis. *Full assembled genomes* Bact.: Bacteria. *Models* Van: Vanilla (constant). Lin: Linear. CNN: convolutional neural network. Ref: reference genome/transcriptome model (only applicable to datasets with a reference).

| Dataset | AR Lin. | AR CNN | AR Ref. | BEAR Van. | BEAR Lin. | BEAR CNN | BEAR Ref. |
| --- | --- | --- | --- | --- | --- | --- | --- |
| YSD1 | 33.73% | 35.86% | 90.8% | 94.69% | **94.75%** | **94.75%** | 94.71% |
| *A. th.* 1 | 35.47% | 35.59% | 53.81% | 86.03% | 86.32% | 86.34% | **86.50%** |
| *A. th.* 2 | 35.32% | 35.61% | 70.41% | 85.36% | 85.71% | **85.77%** | 85.66% |
| *A. th.* 3 | 34.94% | 35.41% | 60.94% | 76.46% | 78.51% | **78.52%** | 77.13% |
| PBMC | 34.36% | 34.76% | 67.39% | 87.83% | **88.16%** | **88.16%** | 87.99% |
| HL | 34.67% | 35.59% | 67.17% | 87.68% | **87.96%** | **87.96%** | 87.82% |
| GBM | 30.71% | 30.9% | 61.3% | 78.99% | 80.44% | 80.42% | **81.43%** |
| HC | 32.98% | 33.54% | – | 83.86% | 85.03% | **85.06%** | – |
| CD | 32.13% | 32.32% | – | 81.72% | 83.30% | **83.32%** | – |
| UC | 32.27% | 32.23% | – | 82.71% | 84.26% | **84.27%** | – |
| Bact. | 33.89% | 34.78% | - | 35.27% | **35.28%** | **35.28%** | - |

### L.4 Further performance results

The maximum marginal likelihood lag *L* (under the vanilla BEAR model) for each dataset is reported in S4. Interestingly, the optimal lags are intermediate between the large kmer lengths (e.g. more than 30) often used for non-probabilistic assembly algorithms (e.g. [57]) and the small kmer lengths (e.g. less than 10) often used as features in clustering or classification algorithms (e.g. [3]). The marginal likelihood was in general strongly peaked at a particular value (Figure S15). Increasing the lag generally led to slightly better performance in terms of both perplexity and accuracy for the non-vanilla BEAR models and the AR models, but (unsurprisingly) worse performance for the vanilla BEAR model; the increases in AR model performance were far from enough to make up the difference with BEAR models (Table S5).

Plots of training loss versus wall clock time for an AR model and the corresponding BEAR model (with the same fixed lag *L*) are shown in Figure S13; the loss for each is normalized by the minimum and maximum values to be comparable (the BEAR model substantially outperforms the AR model). The BEAR model converges at least as fast as the AR model.

To evaluate performance as a function of dataset size, we subsampled reads uniformly at random without replacement from the YSD1 dataset, and retrained the models on the smaller datasets (Figure S14). The original dataset had ~ 1000× coverage of the bacteriophage genome, meaning that on average 1000 reads were observed overlapping each position in the genome. Note that the vanilla BEAR model performance falls off substantially relative to the BEAR model below ~ 3× coverage (in the case of the reference model) (Figure S14BD)

## M Generation details

Here we provide details on the results reported in the **Generating samples** subsection of the results (Section 6).

The CNN BEAR model was trained on the full (combined train/test data) *Arabidopsis thaliana* 1 dataset, with *L* = 17, using identical training parameters as in the performance experiments (Table S2). 50 bases were generated on the end of reads using the maximum *a posteriori* value of *υ*, and conditional on a stop symbol not occurring, i.e. following the distribution

**Table S4:**
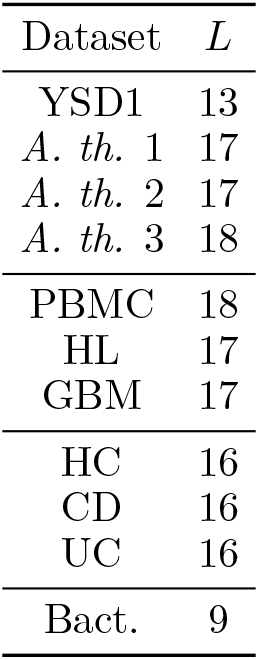
Maximum marginal likelihood lag *L*. Maximum marginal likelihood lag *L* for the vanilla BEAR model. Dataset abbreviations as in Table 1.

**Table S5:** Performance with increasing lag *L*. The symbol † indicates the maximum marginal likelihood lag *L* for the vanilla BEAR model. Dataset abbreviations as in Table 1.

| Perplexity |  |  |  |  |  |  |  |  |
| --- | --- | --- | --- | --- | --- | --- | --- | --- |
| Dataset | Lag | AR Lin. | AR CNN | AR Ref. | BEAR Van. | BEAR Lin. | BEAR CNN | BEAR Ref. |
| YSD1 | 13† | 3.953 | 3.873 | 1.266 | 1.165 | **1.144** | **1.144** | 1.145 |
| YSD1 | 20 | 3.937 | 3.855 | 1.352 | 1.177 | **1.138** | **1.138** | **1.138** |
| Bact. | 9† | 3.831 | 3.794 | - | **3.774** | **3.774** | **3.774** | - |
| Bact. | 12 | 3.807 | 3.772 | - | 3.776 | 3.741 | **3.738** | - |
| Accuracy |  |  |  |  |  |  |  |  |
| Dataset | Lag | AR Lin. | AR CNN | AR Ref. | BEAR Van. | BEAR Lin. | BEAR CNN | BEAR Ref. |
| YSD1 | 13† | 33.73% | 35.86% | 90.8% | 94.69% | **94.75%** | **94.75%** | 94.71% |
| YSD1 | 20 | 34.19% | 36.3% | 87.21% | 94.88% | 94.97% | **94.98%** | 94.91% |
| Bact. | 9† | 33.89% | 34.78% | - | 35.27% | **35.28%** | **35.28%** | - |
| Bact. | 12 | 34.42% | 35.13% | - | 35.54% | 35.86% | **35.93%** | - |

**Figure S13:**
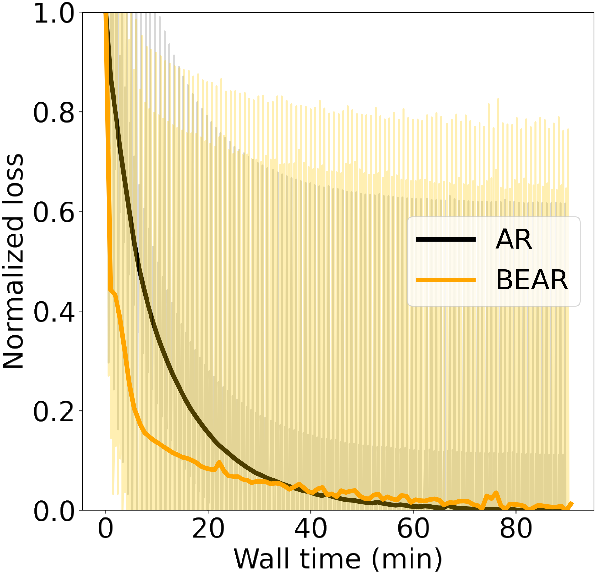
Relative loss (normalized to be between 0 and 1 based on minimum and maximum values) as a function of wall time for a CNN AR model versus the corresponding BEAR model on the YSD1 dataset (*L* = 20).

**Figure S14:**
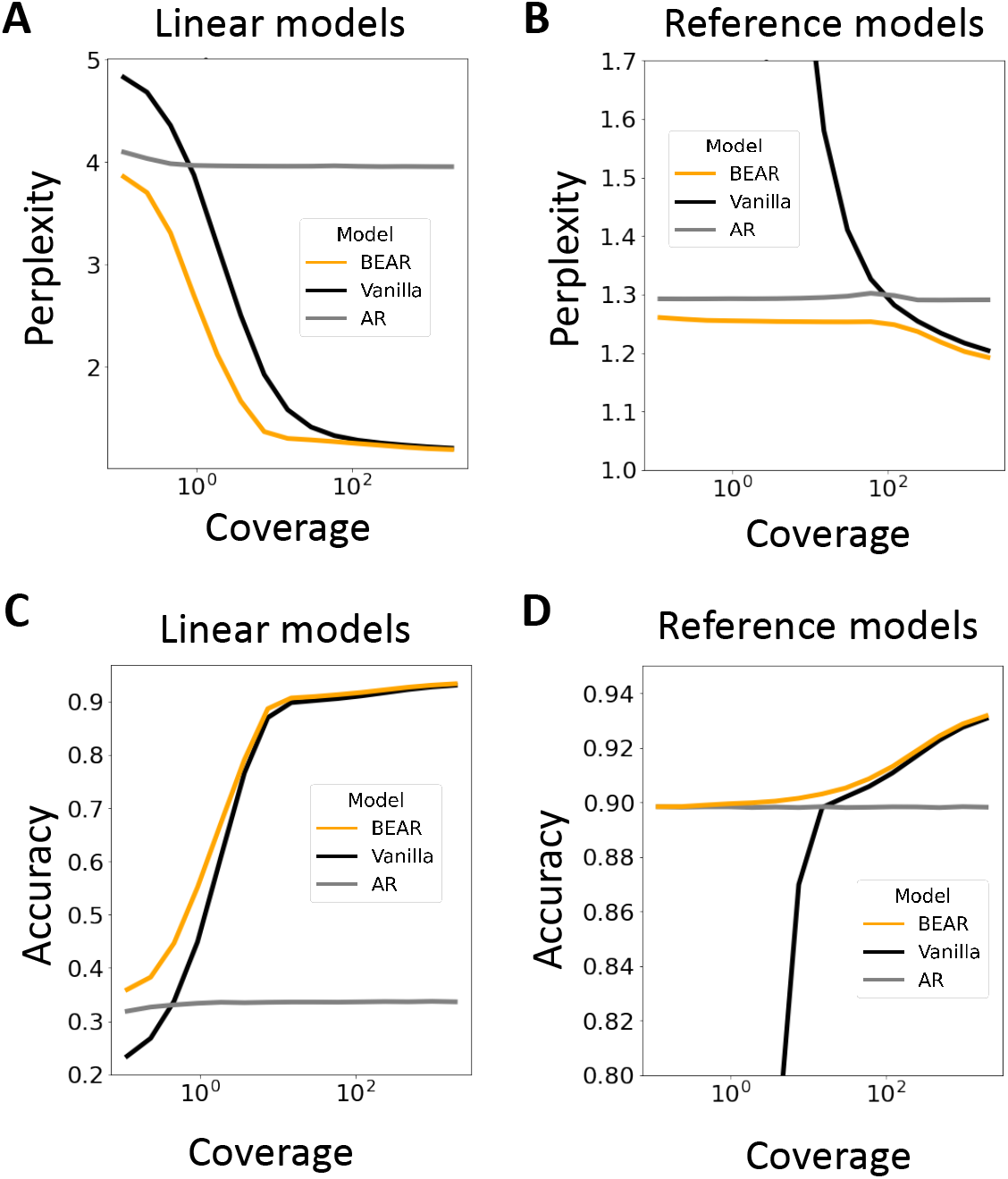
Perplexity (AB) and accuracy (CD) of AR and BEAR models as a function of total dataset size, measured in terms of coverage (coverage is the expected number of reads from each position in the genome; it is linearly proportional to the total number of reads). Subfigures A and C show results for the the linear AR model (and its BEAR embedding), and B and D for the reference-based AR model (and its BEAR embedding). The lag was held fixed in all cases.

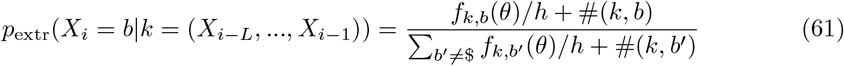

for *b* ≠ $ and *p*(*X_i_* = $|*k*) = 0, where recall #(*k, b*) is the number of times *b* is seen succeeding *k* in the data, and *θ* and *h* are the learned hyperparameters. The values of #(*k, b*) are retrieved from the dataset efficiently using the Jellyfish kmer indexing package [42]. 50 extrapolations each of length 50 were sampled without replacement using the stochastic beam search method proposed by Kool et al. [36].

We performed local assembly using SPAdes, starting from the last 17 bases of the read, and recorded the portion of each scaffold returned by SPAdes that extended in the direction of extrapolation. We used the `--careful` flag in SPAdes, following Voichek and Weigel [67].

The colors in Figure 3A correspond to unique paths through the 17-mer de Bruijn graph. Figure 3B plots the per nucleotide perplexity of the sampled extrapolations, i.e.

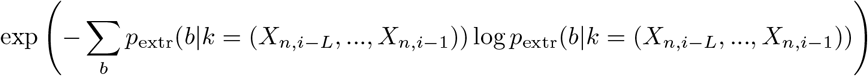

where *n* indexes the sampled extrapolation and *i* the position in the sample.

**Figure S15:**
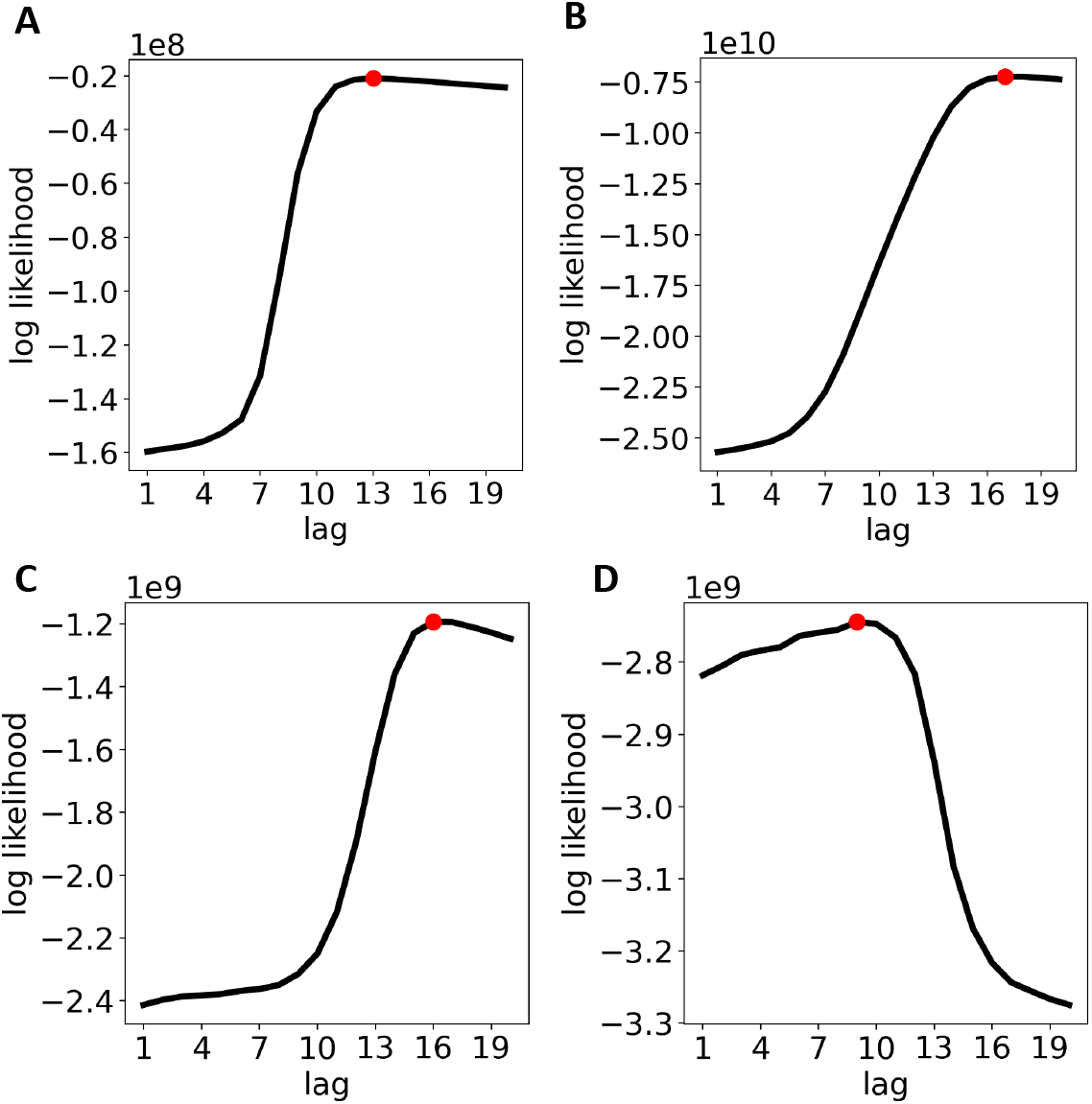
Marginal log likelihood under the vanilla BEAR model as a function of lag *L* for the bacteriophage YSD1 (A), glioblastoma GBM (B), control metagenomic HC (C) and bacteria Bact. (D) datasets. Note the large scale (upper left) of each plot.

## N Visualization details

Here we provide details on the results reported in the **Visualizing data** subsection of the results (Section 6).

### N.1 Latent representation model

As a local latent representation model, we used a categorical probabilistic principal component analysis (pPCA) model, with automatic relevance determination [38, 60]. We trained on kmers (*k_t_, b_t_*) of length *L* + 1 = 18 and used *D* = 20 latent dimensions. The complete model was,

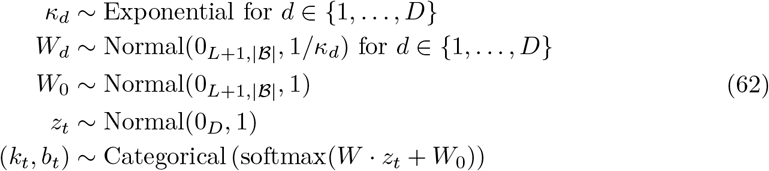

where *t* ∈ {1,…, *T*} runs over all length *L* + 1 kmers in the dataset, 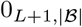 is an 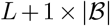 matrix of zeros, and 0_*D*_ is a length *D* vector of zeros. Here the local variable *z_t_* provides a representation associated with the kmer (*k_t_, b_t_*), the global parameter *W* controls the factors of variation, and *κ* determines the relevance of each factor through the variance of the prior on *W*. We trained this latent representation model, and embedded it into a BEAR model, in three stages.

#### Stage 1

First, we performed stochastic variational inference to learn the parameters of the model [34, 37, 50]. In particular, we used normally distributed mean field posterior approximations *q*(*W*),*q*(*z*|*k, b*), and a deterministic approximation to *κ*, and optimized the evidence lower bound (ELBO)

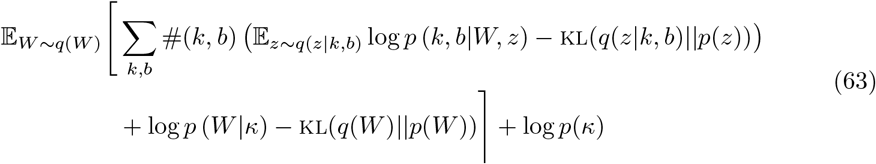

where #(*k, b*) denotes the number of kmers (*k, b*) seen in the data and the sum runs over all 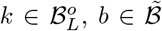. For the local latent variable *z*, we use a guide (recognition network) *q*(*z*|*k, b*) = Normal(*μ*(*k, b*), *σ*(*k, b*)) where *μ*(*k, b*) and *σ*(*k, b*) are each small CNNs. Gradients with respect to the variational approximation parameters were taken using automatic differentiation and the reparameterization trick (elliptical standardization), with one sample for the Monte Carlo approximation at each step.

#### Stage 2

Once the pPCA model was trained, we approximated its conditional distribution. In particular, we obtained a variational approximation to 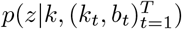, namely *q*(*z*|*k*), by optimizing the evidence lower bound

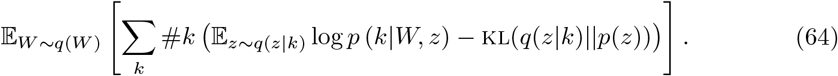

Note that *q*(*W*) was held fixed, at the value learned in stage 1. *q*(*z*|*k*) was parameterized analogously to *q*(*z*|*k, b*). Now we can approximate the conditional distribution of the pPCA model as

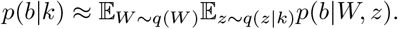

This defines an AR model.

#### Stage 3

Finally, we embedded the conditional pPCA AR model into a BEAR model and optimized *h* via empirical Bayes (note that here we are not using empirical Bayes to train the BEAR model’s embedded AR parameters *θ*, but instead embedding a pretrained AR model). Since the variational distribution *q*(*W*) was highly concentrated at a single point, we used a computationally convenient approximation to the marginal likelihood of the BEAR model, moving the expectation over the global parameters outside the log marginal likelihood:

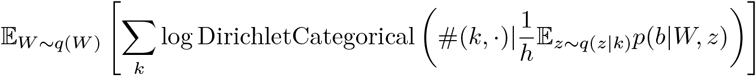

where DirichletCategorical (#(*k*, ·)|*α_k_*) denotes the probability of the count vector #(*k*, ·) under a Dirichlet-Categorical distribution with concentration vector *α_k_*.

#### Training protocol and hyperparameters

The entire variational inference and embedding procedure was implemented using the Edward2 [61] probabilistic programming language with a TensorFlow [2] back-end. We applied the method to the Hodgkin’s lymphoma single cell RNAseq described in section K, using the same train/test split as for the performance results in Section L. Optimization was performed with Adam with a batch size of 125, 000. Gradients were accumulated over 200 steps. The three stages of training described above were repeated iteratively four times until each converged. In each iteration, the first two stages were trained for 5 epochs, and we used a decaying learning rate across iterations {0.02, 0.02, 0.01, 0.005}; the third stage was trained for 100 batches with a constant learning rate of 0.1 across all iterations.

#### Inference results

At the end of training, the BEAR model had a perplexity of 4.276 on heldout data.

### N.2 Visualization and annotation

We next sought to understand in greater depth what the BEAR model had learned in the lymphoma dataset.

#### Reference model

We first aimed to understand how the model’s predictions differed from predictions based on the reference transcriptome. On the full dataset (combined train/test) we compared the log probability of each read under the pPCA BEAR model to the log probability of each read under a vanilla BEAR model trained on the reference transcriptome (Figure 3C; see Datasets.xlsx for details on the reference transcriptome). We found a substantial disparity between the two model’s predictions, with a number of reads having high probability under the BEAR model but low probability according to the reference model.

**Figure S16:**
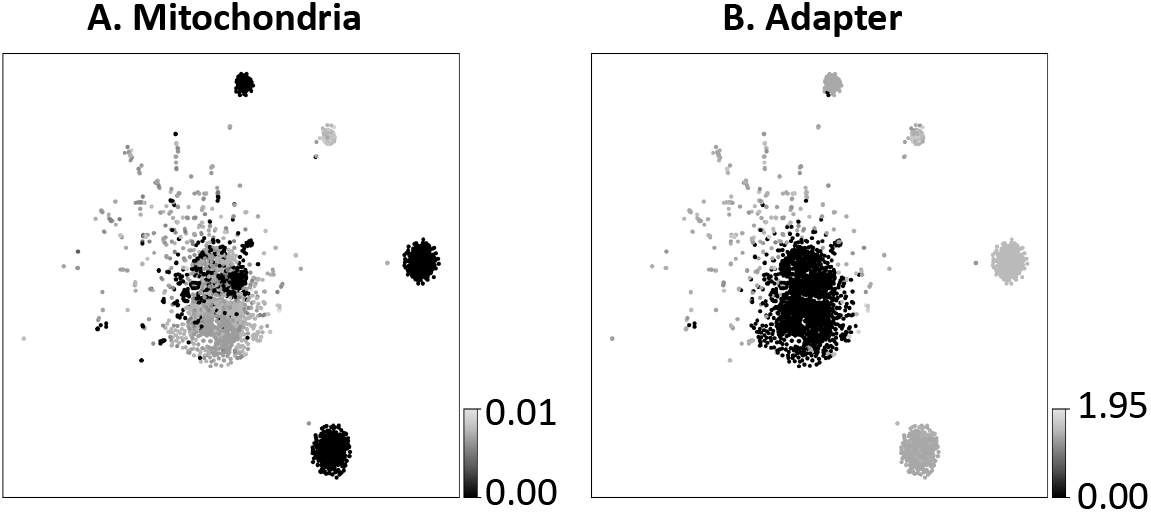
tSNE visualization of a cluster of single cell RNAseq reads colored by (A) latent embedding distance to the mitochondrial reference genome and (B) latent embedding distance from the sequencing adapter.

#### Alignments

Single cell RNAseq analysis often begins by aligning reads to the reference transcriptome; reads that do not align are typically discarded from further analysis. We performed alignments on the read dataset with hisat2 [32] using parameters `-reorder -no-hd -n-ceil L,0,0.001 -no-sq -k 1 -p 4` and with the default hisat2 *Homo sapiens* GRCh38 genome index with transcripts and SNPs, available at https://genome-idx.s3.amazonaws.com/hisat/grch38_snptran.tar.gz. Whether or not each read was successfully aligned is indicated in Figure 3C. We observe that many of the reads with low probability under both the pPCA BEAR model and the reference model are unaligned. We also observed a cluster with a large number of unaligned reads, with high probability under the pPCA BEAR model and relatively low probability under the reference model. We focused on a subset of this cluster with particularly high probabilities under the pPCA BEAR model for follow-up visualization (black box in Figure 3C).

#### Visualization

The pPCA model provides a latent embedding of kmers in a *D* = 20 dimensional continuous space. We sought to visualize the representation of each sequence’s kmers in a low dimensional space. To compare two sequences *X,X*′, we defined a measure of dissimilarity,

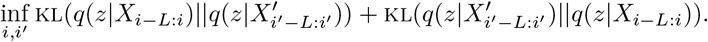

where *i* > *L* and *i*′ > *L* index positions in *X* and *X*′ respectively. This dissimilarity measure was used to define a distance matrix over reads in the Hodgkin’s lymphoma dataset, which was passed to tSNE [63] to obtain a low-dimensional visualization (Figure 3D).

#### Annotation

Observing the clusters in Figure 3D, we sought to determine where the reads in each cluster likely originated from, and, by implication, what the reference transcriptome model had trouble explaining in the data. We started by using NCBI’s BLAST tool [8] to search for likely sources, and found hits against the mitochondrial genome and the transcript of the gene *JUND*, part of the AP-1 early response transcription factor. We found that the mitochondrial reads are from a nonreference haplotype, which explains why the reference model gave them low probability. The low likelihood of the *JUND* reads under the reference was due to a TG repeat region in the 3’ UTR; similar repeats are present in many variations in different transcripts, thus the particular kmer-base transitions in this case become less likely. We also observed that many reads were chimeric, consisting of fusions of sequences from various parts of the transcriptome with some portion of the sequence CTGTCTCTTAT-ACACATCTCTGAACGGGCTGGCAAGGCAGACCG. The prefix CTGTCTCTTATACACATCT is a standard Illumina Nextera adapter sequence https://support-docs.illumina.com/SHARE/AdapterSeq/illumina-adapter-sequences.pdf, and the remainder of the sequence is presumably part of the primer. The adapter is an experimental artifact (presumably left in the read data due to inaccurate read trimming and quality control), and so is not part of the reference human transcriptome.

We used the same dissimilarity measure as above to compare reads to the mitochondria reference genome (Datasets.xlsx) and to the adapter sequence CTGTCTCTTATACACATCTCTGAACGGGCTGGCAAGGCAGACCG (Figure S16). (The distance to each of these sequences was taken to be the minimum of the distance to the forward and reverse complements.) Figure S16, along with the BLAST results for *JUND*, were the basis for the annotations in Figure 3D.

## O Hypothesis tests details

Here we provide details on the results reported in the **Testing hypotheses** subsection of the results (Section 6).

### O.1 Kidney transplant metagenomics

The Schreiber et al. [55] data is available for public download, as detailed in Datasets.xlsx. The read data was pre-sorted into viral and non-viral reads, but we pooled each of these to reconstruct the full sequencing experiment. We compared the day zero timepoint, i.e. before transplant, to the 4-6 week timepoint, i.e. after transplant, for each patient for which samples from both were available (note this did not include all patients in the study). We used the BEAR two-sample test, with the Jeffreys prior on *υ*, and a truncated uniform prior over lags 1 ≤ *L* ≤ 20. We cross referenced our two-sample test results with whether Schreiber et al. [55] determined there to be likely JC polyomavirus (JCPyV) transmission.

The results are shown in Table S6, and suggest that JCPyV transmission is associated with an overall shift in the patient microbiome at the sequence level. Patients indicated with an asterisk were diagnosed as having JCPyV before receiving the transplant, and thus the determination of whether the transplant transmitted JCPyV is less certain; for patient wdk036, phylogenetic analysis suggested that the transplant did transmit JCPyV, while for jns976 phylogenetic analysis suggested that it did not. Although the two-sample test results show close correlation with whether or not there was transmission, we caveat them by noting that for very small lags the Bayes factor rejects the null hypothesis for all patients; the question of the most “biologically relevant” prior on the lag *L* is an open question.

### O.2 *A. thaliana* hypothesis tests

#### Goodness-of-fit test

We trained reference-based AR models (described in Section L.1) via maximum likelihood on each *A. thaliana* sequencing dataset (the full dataset, with train/test subsets combined). We used *L* =17 in the AR model for all three datasets (corresponding the vanilla BEAR maximum marginal likelihood lag for two datasets, see Table S4). We embedded each trained AR model into a BEAR model to construct a goodness-of-fit test (i.e. we used the learned *f*(*θ*)). We fixed *L* =17 in the BEAR model (i.e. a deterministic prior over *L*) to determine if there was misspecification at the same resolution as the AR model. Figure 3E plots the Bayes factor as a function of *h*.

#### Two-sample tests

We simulated sequencing reads based on the *A. thaliana* reference genome (Datasets.xlsx) using the ART Illumina [29] simulator with parameters `-ss HS20 -p -l 100 -m 200 -s 10 -f 30`. We simulated roughly the same number of reads as was in each real dataset. We examined the Bayes factor 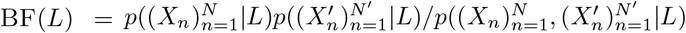, computed using vanilla BEAR models for each term (Figure 3F). As control experiments, we cut each dataset (and the simulated data) in half, and compared each of these halves to each other using the same two-sample test; as shown by the dotted lines in Figure 3F, the two-sample test correctly accepts the null hypothesis in these cases.

**Table S6:**
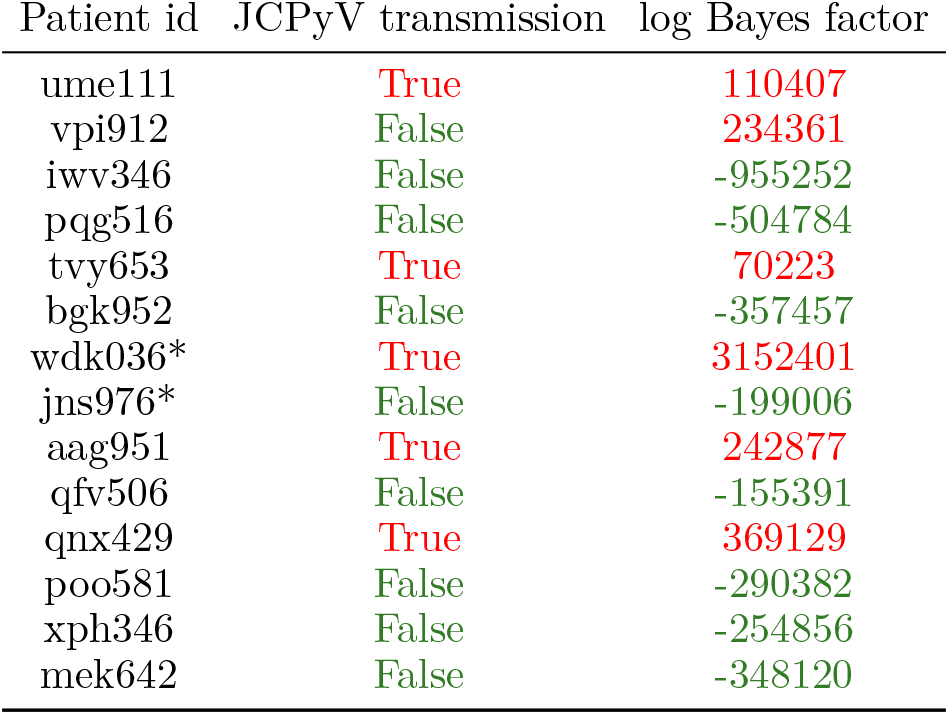
BEAR two-sample test results, performed on patient metagenome samples from before and after kidney transplant. Bayes factors that reject the null hypothesis are colored red, for easy comparison with whether or not JC polyomavirus (JCPyV) transmission was detected. Asterisks * indicate patients that were already infected with JCPyV before the transplant occurred.

#### Individual log likelihood ratio

To understand in detail the differences between the real and simulated data, we computed the conditional individual Bayes factor 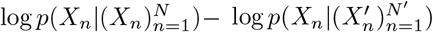 where 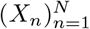 is the real data and 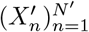 the simulated data. We approximated the log likelihood using the maximum *a posteriori* value of the transition parameter *υ* under the vanilla BEAR model, and fixed *L* = 17. Computing this likelihood efficiently for each read requires retrieving counts #(*k*, ·) for each kmer *k* in the read, which we accomplished using the Jellyfish kmer indexing package [42]. Histograms of the log likelihood ratio of each read *X_n_* in two of the *A. thaliana* datasets are shown in Figure 3G (gray).

#### Annotation

Observing the distinct peaks in Figure 3G, we sought to determine where the reads in each originated from. We discovered that many reads in the outlier peak from *A. thaliana* 1 matched *Bacillus cereus*, using NCBI’s BLAST tool [8]. To annotate the clusters further, we aligned the reads to reference sequences for centromeres, chloroplasts, and *B. cereus*, as well as (if the read did not align to one of these) the reference *A. thaliana* genome (reference sequences are listed in Datasets.xlsx). Alignments were performed using `hisat2` on paired end read data using parameters `-reorder -no-hd -n-ceil L,0,0.001 -no-sq -k 1 -p 4` to facilitate subsequent analysis and remove reads with ambiguous bases. The alignment to the centromere included the parameter `-mp 1,1` to allow lower quality alignments. Histograms of the set of reads that align to each reference are shown (stacked on top of one another, not overlayed) in Figure 3G.

## Footnotes

2 We do not need to assume 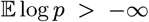 as we may define in this case 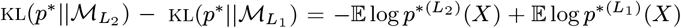 which we will see is bounded by the moment bound assumption 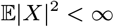.

3 It is not crucial that maximizers of the marginal likelihood exist for any of the result below: the results below hold assuming only that *h_N_, θ_N_* are approximate maximizers, i.e. 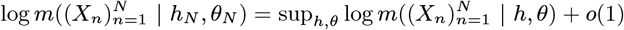 or in slightly altered form swapping the *o*(1) for *OP*(1).

4 Since 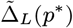 is compact, the set of polynomials with rational coefficients, 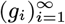 is dense in the space of continuous functions under the infinite norm. Pick 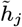 to have 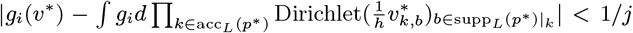 for all *i* ≤ *j* and then 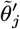 to have 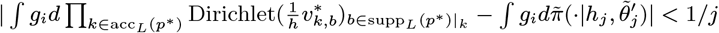 for all *i* ≤ *j*.

5 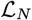 is stochastic due to *r_N_*, but since *h_N_N^β^* → ∞ for any *β* > 0, using the expansion in equation 20, one may show that the #*k* in *r_N_* can be replaced with 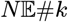 incurring only a penalty of *O_P_*(*N*^−1/2+*ϵ*^).

6 Note that the KL term in 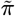 can be dominated by 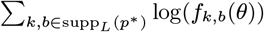.

## References

[1] 1001 Genomes Consortium. 1,135 genomes reveal the global pattern of polymorphism in arabidopsis thaliana. Cell, 166(2):481–491, July 2016.

[2] M. Abadi, P. Barham, J. Chen, Z. Chen, A. Davis, J. Dean, M. Devin, S. Ghemawat, G. Irving, M. Isard, and Others. Tensorflow: A system for large-scale machine learning. In 12th USENIX symposium on operating systems design and implementation (OSDI 16), pages 265–283. usenix.org, 2016.

[3] E. B. Alsop and J. Raymond. Resolving Prokaryotic Taxonomy without rRNA: Longer Oligonucleotide Word Lengths Improve Genome and Metagenome Taxonomic Classification. PLoS ONE, 8(7), 2013.

[4] J. L. Ba, J. R. Kiros, and G. E. Hinton. Layer normalization. July 2016.

[5] A. Bankevich, S. Nurk, D. Antipov, A. A. Gurevich, M. Dvorkin, A. S. Kulikov, V. M. Lesin, S. I. Nikolenko, S. Pham, A. D. Prjibelski, A. V. Pyshkin, A. V. Sirotkin, N. Vyahhi, G. Tesler, M. A. Alekseyev, and P. A. Pevzner. SPAdes: a new genome assembly algorithm and its applications to single-cell sequencing. J. Comput. Biol., 19(5):455–477, May 2012.

[6] J. O. Berger and A. Guglielmi. Bayesian and conditional frequentist testing of a parametric model versus nonparametric alternatives. Journal of the American Statistical Association, 96(453):174–184, 2001.

[7] S. Biswas, G. Khimulya, E. C. Alley, K. M. Esvelt, and G. M. Church. Low-N protein engineering with data-efficient deep learning. Nat. Methods, 18(4):389–396, 2021.

[8] G. M. Boratyn, C. Camacho, P. S. Cooper, G. Coulouris, A. Fong, N. Ma, T. L. Madden, W. T. Matten, S. D. McGinnis, Y. Merezhuk, Y. Raytselis, E. W. Sayers, T. Tao, J. Ye, and I. Zaretskaya. BLAST: a more efficient report with usability improvements. Nucleic Acids Res., 41(Web Server issue):W29–33, July 2013.

[9] S. F. Chen and J. Goodman. An empirical study of smoothing techniques for language modeling. Comput. Speech Lang., 13(4):359–394, Oct. 1999.

[10] P. J. A. Cock, T. Antao, J. T. Chang, B. A. Chapman, C. J. Cox, A. Dalke, I. Friedberg, T. Hamelryck, F. Kauff, B. Wilczynski, and M. J. L. de Hoon. Biopython: freely available python tools for computational molecular biology and bioinformatics. Bioinformatics, 25(11):1422–1423, June 2009.

[11] P. E. C. Compeau, P. A. Pevzner, and G. Tesler. How to apply de bruijn graphs to genome assembly. Nat. Biotechnol., 29(11):987–991, Nov. 2011.

[12] I. Cortés-Ciriano, J. J.-K. Lee, R. Xi, D. Jain, Y. L. Jung, L. Yang, D. Gordenin, L. J. Klimczak, C.-Z. Zhang, D. S. Pellman, PCAWG Structural Variation Working Group, P. J. Park, and PCAWG Consortium. Comprehensive analysis of chromothripsis in 2,658 human cancers using whole-genome sequencing. Nat. Genet., 52(3):331–341, Mar. 2020.

[13] S. Darmanis, S. A. Sloan, D. Croote, M. Mignardi, S. Chernikova, P. Samghababi, Y. Zhang, N. Neff, M. Kowarsky, C. Caneda, G. Li, S. D. Chang, I. D. Connolly, Y. Li, B. A. Barres, M. H. Gephart, and S. R. Quake. Single-Cell RNA-Seq analysis of infiltrating neoplastic cells at the migrating front of human glioblastoma. Cell Rep., 21 (5):1399–1410, Oct. 2017.

[14] A. P. Dawid. Posterior model probabilities. In P. S. Bandyopadhyay and M. R. Forster, editors, Philosophy of Statistics, volume 7, pages 607–630. North-Holland, Amsterdam, Jan. 2011.

[15] J. V. Dillon, I. Langmore, D. Tran, E. Brevdo, S. Vasudevan, D. Moore, B. Patton, A. Alemi, M. Hoffman, and R. A. Saurous. TensorFlow distributions. Nov. 2017.

[16] V. B. Dubinkina, D. S. Ischenko, V. I. Ulyantsev, A. V. Tyakht, and D. G. Alexeev. Assessment of k-mer spectrum applicability for metagenomic dissimilarity analysis. BMC Bioinformatics, 17:38, Jan. 2016.

[17] R. A. Dunstan, D. Pickard, S. Dougan, D. Goulding, C. Cormie, J. Hardy, F. Li, R. Grin-ter, K. Harcourt, L. Yu, J. Song, F. Schreiber, J. Choudhary, S. Clare, F. Coulibaly, R. A. Strugnell, G. Dougan, and T. Lithgow. The flagellotropic bacteriophage YSD1 targets salmonella typhi with a chi-like protein tail fibre. Mol. Microbiol., 112(6):1831–1846, Dec. 2019.

[18] R. Durbin, S. Eddy, A. Krogh, and A. Mitchison. Biological Sequence Analysis. 1998.

[19] J. Frazer, P. Notin, M. Dias, A. Gomez, K. Brock, Y. Gal, and D. Marks. Large-scale clinical interpretation of genetic variants using evolutionary data and deep learning. Dec. 2020.

[20] S. Geman and C.-R. Hwang. Nonparametric maximum likelihood estimation by the method of sieves. The Annals of Statistics, 10(2):401–414, 1982.

[21] S. Ghosal, J. K. Ghosh, and A. W. van der Vaart. Convergence rates of posterior distributions. Ann. Stat., 28(2):500–531, 2000.

[22] J. Ghosh and R. Ramamoorthi. Bayesian Nonparametrics. 2003.

[23] P. Gopalan, W. Hao, D. M. Blei, and J. D. Storey. Scaling probabilistic models of genetic variation to millions of humans. Nat. Genet., 48(12):1587–1590, Dec. 2016.

[24] R. M. Gray. Entropy and Information Theory. Springer Science & Business Media, Jan. 2011.

[25] A. Gretton, K. M. Borgwardt, M. J. Rasch, B. Schölkopf, and A. Smola. A kernel two-sample test. J. Mach. Learn. Res., 13:723–773, 2012.

[26] C. R. Harris, K. J. Millman, S. J. van der Walt, R. Gommers, P. Virtanen, D. Cournapeau, E. Wieser, J. Taylor, S. Berg, N. J. Smith, R. Kern, M. Picus, S. Hoyer, M. H. van Kerkwijk, M. Brett, A. Haldane, J. F. del Río, M. Wiebe, P. Peterson, P. Gérard-Marchant, K. Sheppard, T. Reddy, W. Weckesser, H. Abbasi, C. Gohlke, and T. E. Oliphant. Array programming with NumPy. Nature, 585(7825):357–362, Sept. 2020. doi: 10.1038/s41586-020-2649-2. URL https://doi.org/10.1038/s41586-020-2649-2.

[27] C. C. Holmes, F. Caron, J. E. Griffin, and D. A. Stephens. Two-sample bayesian nonparametric hypothesis testing. Bayesian Anal., 10(2):297–320, June 2015.

[28] T. A. Hopf, J. B. Ingraham, F. J. Poelwijk, C. P. I. Schärfe, M. Springer, C. Sander, and D. S. Marks. Mutation effects predicted from sequence co-variation. Nat. Biotechnol., 35(2):128–135, Feb. 2017.

[29] W. Huang, L. Li, J. R. Myers, and G. T. Marth. ART: a next-generation sequencing read simulator. Bioinformatics, 28(4):593–594, Feb. 2012.

[30] Z. Iqbal, M. Caccamo, I. Turner, P. Flicek, and G. McVean. De novo assembly and genotyping of variants using colored de bruijn graphs. Nat. Genet., 44(2):226–232, Jan. 2012.

[31] P. E. Jacob, L. M. Murray, C. C. Holmes, and C. P. Robert. Better together? statistical learning in models made of modules. Aug. 2017.

[32] D. Kim, J. M. Paggi, C. Park, C. Bennett, and S. L. Salzberg. Graph-based genome alignment and genotyping with HISAT2 and HISAT-genotype. Nat. Biotechnol., 37(8): 907–915, Aug. 2019.

[33] D. P. Kingma and J. Ba. Adam: A method for stochastic optimization. In ICLR, 2015.

[34] D. P. Kingma and M. Welling. Auto-Encoding variational bayes. Dec. 2013.

[35] M. Kokot, M. Dlugosz, and S. Deorowicz. KMC 3: counting and manipulating k-mer statistics. Bioinformatics, 33(17):2759–2761, Sept. 2017.

[36] W. Kool, H. van Hoof, and M. Welling. Stochastic beams and where to find them: The Gumbel-Top-k trick for sampling sequences without replacement. In International Conference on Machine Learning, pages 3499–3508. PMLR, 2019.

[37] A. Kucukelbir and D. M. Blei. Population empirical bayes. In Uncertainty in Artificial Intelligence, 2015.

[38] A. Kucukelbir, D. Tran, R. Ranganath, A. Gelman, and D. M. Blei. Automatic differentiation variational inference. J. Mach. Learn. Res., 18(14):1–45, Jan. 2017.

[39] Y. Li, K. Swersky, and R. Zemel. Generative moment matching networks. In International Conference on Machine Learning, pages 1718–1727. PMLR, 2015.

[40] J. R. Lloyd and Z. Ghahramani. Statistical model criticism using kernel two sample tests. In Advances in Neural Information Processing Systems, pages 829–837, 2015.

[41] J. Lloyd-Price, A. Mahurkar, G. Rahnavard, J. Crabtree, J. Orvis, A. B. Hall, A. Brady, H. H. Creasy, C. McCracken, M. G. Giglio, D. McDonald, E. A. Franzosa, R. Knight, O. White, and C. Huttenhower. Strains, functions and dynamics in the expanded human microbiome project. Nature, 550(7674):61–66, Oct. 2017.

[42] G. Marçais and C. Kingsford. A fast, lock-free approach for efficient parallel counting of occurrences of k-mers. Bioinformatics, 27(6):764–770, Mar. 2011.

[43] J. W. Miller. Asymptotic normality, concentration, and coverage of generalized posteriors. July 2019.

[44] S. Mohamed and B. Lakshminarayanan. Learning in implicit generative models. Oct. 2016.

[45] R. E. Mukamel, R. E. Handsaker, M. A. Sherman, A. R. Barton, Y. Zheng, S. A. McCarroll, and P.-R. Loh. Protein-coding repeat polymorphisms strongly shape diverse human phenotypes. Jan. 2021.

[46] N. A. O’Leary, M. W. Wright, J. R. Brister, S. Ciufo, D. Haddad, R. McVeigh, B. Rajput, B. Robbertse, B. Smith-White, D. Ako-Adjei, A. Astashyn, A. Badretdin, Y. Bao, O. Blinkova, V. Brover, V. Chetvernin, J. Choi, E. Cox, O. Ermolaeva, C. M. Farrell, T. Goldfarb, T. Gupta, D. Haft, E. Hatcher, W. Hlavina, V. S. Joardar, V. K. Kodali, W. Li, D. Maglott, P. Masterson, K. M. McGarvey, M. R. Murphy, K. O’Neill, S. Pujar, S. H. Rangwala, D. Rausch, L. D. Riddick, C. Schoch, A. Shkeda, S. S. Storz, H. Sun, F. Thibaud-Nissen, I. Tolstoy, R. E. Tully, A. R. Vatsan, C. Wallin, D. Webb, W. Wu, M. J. Landrum, A. Kimchi, T. Tatusova, M. DiCuccio, P. Kitts, T. D. Murphy, and K. D. Pruitt. Reference sequence (RefSeq) database at NCBI: current status, taxonomic expansion, and functional annotation. Nucleic Acids Res., 44(D1):D733–45, Jan. 2016.

[47] K. Papineni, S. Roukos, T. Ward, and W.-J. Zhu. BLEU: a method for automatic evaluation of machine translation. In Annual Meeting of the Association for Computational Linguistics, pages 311–318, 2002.

[48] S. Petrone, J. Rousseau, and C. Scricciolo. Bayes and empirical bayes: do they merge? Biometrika, 101(2):285–302, 2014.

[49] J. K. Pritchard, M. Stephens, and P. Donnelly. Inference of population structure using multilocus genotype data. Genetics, 155(2):945–959, June 2000.

[50] D. J. Rezende, S. Mohamed, and D. Wierstra. Stochastic backpropagation and approximate inference in deep generative models. In Proceedings of the 31st International Conference on Machine Learning, 2014.

[51] A. J. Riesselman, J. B. Ingraham, and D. S. Marks. Deep generative models of genetic variation capture the effects of mutations. Nat. Methods, 15(10):816–822, Oct. 2018.

[52] J. Rousseau. On the Frequentist Properties of Bayesian Nonparametric Methods. Annual Review of Statistics and Its Application, 3:211–231, 2016.

[53] J. Rousseau and K. Mengersen. Asymptotic behaviour of the posterior distribution in overfitted mixture models. J. R. Stat. Soc. Series B Stat. Methodol., 73(5):689–710, Nov. 2011.

[54] W. P. Russ, M. Figliuzzi, C. Stocker, P. Barrat-Charlaix, M. Socolich, P. Kast, D. Hilvert, R. Monasson, S. Cocco, M. Weigt, and R. Ranganathan. An evolution-based model for designing chorismate mutase enzymes. Science, 369:440–445, 2020.

[55] P. W. Schreiber, V. Kufner, K. Hübel, S. Schmutz, O. Zagordi, A. Kaur, C. Bayard, M. Greiner, A. Zbinden, R. Capaul, J. Böni, H. H. Hirsch, T. F. Mueller, N. J. Mueller, A. Trkola, and M. Huber. Metagenomic virome sequencing in living donor and recipient kidney transplant pairs revealed JC polyomavirus transmission. Clin. Infect. Dis., 69 (6):987–994, Aug. 2019.

[56] J.-E. Shin, A. J. Riesselman, A. W. Kollasch, C. McMahon, E. Simon, C. Sander, A. Manglik, A. C. Kruse, and D. S. Marks. Protein design and variant prediction using autoregressive generative models. Nat. Commun., 12(1):2403, Apr. 2021.

[57] J. T. Simpson, K. Wong, S. D. Jackman, J. E. Schein, S. J. M. Jones, and I. Birol. ABySS: a parallel assembler for short read sequence data. Genome Res., 19(6):1117–1123, June 2009.

[58] P. J. Stephens, C. D. Greenman, B. Fu, F. Yang, G. R. Bignell, L. J. Mudie, E. D. Pleasance, K. W. Lau, D. Beare, L. A. Stebbings, S. McLaren, M.-L. Lin, D. J. McBride, I. Varela, S. Nik-Zainal, C. Leroy, M. Jia, A. Menzies, A. P. Butler, J. W. Teague, M. A. Quail, J. Burton, H. Swerdlow, N. P. Carter, L. A. Morsberger, C. Iacobuzio-Donahue, G. A. Follows, A. R. Green, A. M. Flanagan, M. R. Stratton, P. A. Futreal, and P. J. Campbell. Massive genomic rearrangement acquired in a single catastrophic event during cancer development. Cell, 144(1):27–40, Jan. 2011.

[59] D. J. Sutherland, H. Y. Tung, H. Strathmann, S. De, A. Ramdas, A. Smola, and A. Gretton. Generative models and model criticism via optimized maximum mean discrepancy. In International Conference on Learning Representations. arxiv.org, 2017.

[60] M. E. Tipping and C. M. Bishop. Probabilistic principal component analysis. J. R. Stat. Soc. Series B Stat. Methodol., 61(3):611–622, Aug. 1999.

[61] D. Tran, M. Hoffman, D. Moore, C. Suter, S. Vasudevan, A. Radul, M. Johnson, and R. A. Saurous. Simple, distributed, and accelerated probabilistic programming. In Neural Information Processing Systems, 2018.

[62] A. van den Oord, S. Dieleman, H. Zen, K. Simonyan, O. Vinyals, A. Graves, N. Kalch-brenner, A. Senior, and K. Kavukcuoglu. WaveNet: A generative model for raw audio. Sept. 2016.

[63] L. van der Maaten. Visualizing data using t-SNE. J. Mach. Learn. Res., 9:2579–2605, 2008.

[64] A. W. van der Vaart. Asymptotic Statistics. 1998.

[65] R. Vershynin. High-Dimensional Probability: An Introduction with Applications in Data Science. 2020.

[66] P. Virtanen, R. Gommers, T. E. Oliphant, M. Haberland, T. Reddy, D. Cournapeau, E. Burovski, P. Peterson, W. Weckesser, J. Bright, S. J. van der Walt, M. Brett, J. Wilson, K. J. Millman, N. Mayorov, A. R. J. Nelson, E. Jones, R. Kern, E. Larson, C. J. Carey, Ì. Polat, Y. Feng, E. W. Moore, J. VanderPlas, D. Laxalde, J. Perktold, R. Cimrman, I. Henriksen, E. A. Quintero, C. R. Harris, A. M. Archibald, A. H. Ribeiro, F. Pedregosa, P. van Mulbregt, and SciPy 1.0 Contributors. SciPy 1.0: Fundamental Algorithms for Scientific Computing in Python. Nature Methods, 17:261–272, 2020. doi: 10.1038/s41592-019-0686-2.

[67] Y. Voichek and D. Weigel. Identifying genetic variants underlying phenotypic variation in plants without complete genomes. Nature Genetics, 52(5):534–540, 2020.

[68] E. N. Weinstein and D. S. Marks. A structured observation distribution for generative biological sequence prediction and forecasting. Feb. 2021.

